# Ultra-Fast Bioorthogonal Spin-Labeling and Distance Measurements in Mammalian Cells Using Small, Genetically Encoded Tetrazine Amino Acids

**DOI:** 10.1101/2023.01.26.525763

**Authors:** Subhashis Jana, Eric G. B. Evans, Hyo Sang Jang, Shuyang Zhang, Hui Zhang, Andrzej Rajca, Sharona E. Gordon, William N. Zagotta, Stefan Stoll, Ryan A. Mehl

## Abstract

Studying protein structures and dynamics directly in the cellular environments in which they function is essential to fully understand the molecular mechanisms underlying cellular processes. Site-directed spin-labeling (SDSL)—in combination with double electron–electron resonance (DEER) spectroscopy—has emerged as a powerful technique for determining both the structural states and the conformational equilibria of biomacromolecules. In-cell DEER spectroscopy on proteins in mammalian cells has thus far not been possible due to the notable challenges of spin-labeling in live cells. In-cell SDSL requires exquisite biorthogonality, high labeling reaction rates and low background signal from unreacted residual spin label. While the bioorthogonal reaction must be highly specific and proceed under physiological conditions, many spin labels display time-dependent instability in the reducing cellular environment. Additionally, high concentrations of spin label can be toxic. Thus, an exceptionally fast bioorthogonal reaction is required that can allow for complete labeling with low concentrations of spin-label prior to loss of signal. Here we utilized genetic code expansion to site-specifically encode a novel family of small, tetrazine-bearing non-canonical amino acids (Tet-v4.0) at multiple sites in green fluorescent protein (GFP) and maltose binding protein (MBP) expressed both in *E. coli* and in human HEK293T cells. We achieved specific and quantitative spin-labeling of Tet-v4.0-containing proteins by developing a series of strained *trans*-cyclooctene (sTCO)-functionalized nitroxides—including a *gem*-diethyl-substituted nitroxide with enhanced stability in cells—with rate constants that can exceed 10^6^ M^−1^ s^−1^. The remarkable speed of the Tet-v4.0/sTCO reaction allowed efficient spin-labeling of proteins in live HEK293T cells within minutes, requiring only sub-micromolar concentrations of sTCO–nitroxide added directly to the culture medium. DEER recorded from intact cells revealed distance distributions in good agreement with those measured from proteins purified and labeled *in vitro*. Furthermore, DEER was able to resolve the maltose-dependent conformational change of Tet-v4.0-incorporated and spin-labeled MBP *in vitro* and successfully discerned the conformational state of MBP within HEK293T cells. We anticipate the exceptional reaction rates of this system, combined with the relatively short and rigid side chains of the resulting spin labels, will enable structure/function studies of proteins directly in cells, without any requirements for protein purification.

**TOC:** 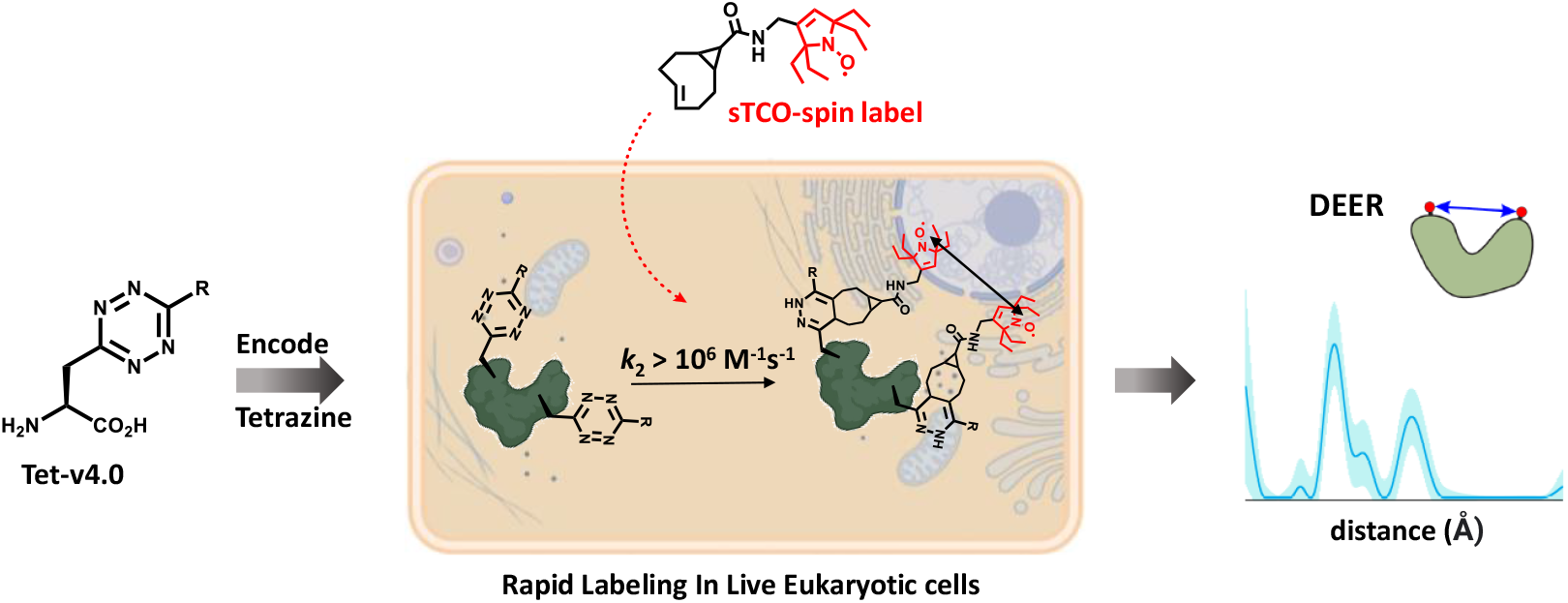

## Introduction

Measuring the structures and dynamics of proteins directly within cells has been a long-standing challenge in structural biology. The complex and crowded environment of the cell can impact the energetic landscape of proteins and may contribute significantly to the molecular mechanisms of protein function and dysfunction.^1–6^ However, few biophysical techniques are capable of residue-specific structural information in the cellular environment.^7^ Site-directed spin-labeling (SDSL), in combination with electron paramagnetic resonance (EPR) spectroscopy, holds considerable promise for elucidating structural dynamics in a wide range of proteins and macromolecular complexes in the cellular context.^8,9^ In particular, double electron–electron resonance (DEER) spectroscopy can inform on both the structural states and the conformational equilibria of biomacromolecules. By measuring the distance-dependent magnetic dipole interaction between pairs of spin labels introduced site-specifically into the protein sequence, DEER provides the full distribution of distances between the labels in a protein ensemble.^10,11^ SDSL of proteins typically relies on cysteine conjugation using thiosulfonate-, maleimide-, or iodoacetamide-functionalized spin labels; however, site-specific conjugation within cells requires labeling reactions and reagents that are bioorthogonal to avoid off-target labeling of cellular components. Consequently, most in-cell EPR studies to date have relied on the delivery of *in vitro* purified and spin-labeled proteins back into cells by microinjection,^12–16^ electroporation,^6,17,18^ osmotic shock,^19^ or other permeation techniques.^20–23^

The site-specific incorporation of non-canonical amino acids (ncAAs) by genetic code expansion (GCE) is a powerful method that has been utilized to introduce novel bioorthogonal chemistries into proteins.^24^ Previous studies using GCE for spin-labeling have employed ketone condensations,^25–29^ 3+2 azide-alkyne cycloadditions,^17,30,31^ Suzuki‒Miyaura couplings,^32^ quinone methide Michael additions,^33^ and inverse electron demand Diels‒Alder (IEDDA) cycloadditions.^34–36^ In addition, ncAAs that themselves include the EPR-active spin probe have been developed, in principle eliminating the need for subsequent spin-labeling.^37–39^ Despite significant progress in bioorthogonal spin-labeling methods over the past decade, their use in in-cell structural studies has been primarily limited to prokaryotic expression systems and has suffered from low yields, slow reaction rates, requirements for harsh reaction conditions or catalysts, and the inherent instability of nitroxide ncAAs or spin labels in the reducing intracellular environment.^37,40,41^

The advancement of site-directed spin-labeling methodologies for EPR-based structural studies within mammalian cells is reliant on several prerequisites, namely: (1) extremely rapid bioorthogonal coupling reactions that can access the high labeling efficiency needed at the typically low protein concentrations (nM – low-μM) in mammalian cells and the low spin-labeling reagent concentrations needed to avoid cellular toxicity, (2) high bioorthogonal reaction selectivity to minimize off-target labeling and background interference from unreacted label, (3) structurally short and rigid connectivity between the attached spin label and the protein peptide backbone to minimize broad distance distributions that can negatively impact structural interpretation in DEER spectroscopy. In the case of the most commonly employed spin labels, nitroxides, the experimental time window from spin label addition to EPR measurement is also a key experimental parameter, as nitroxides undergo a time-dependent chemical reduction in cellular environments.^42,43^ As such, the bioorthogonal coupling reaction of nitroxide labels within mammalian cells must be sufficiently fast to outcompete cellular nitroxide reduction.

The strain-promoted IEDDA reaction between tetrazines and cyclic alkenes has recently emerged as a promising approach for in-cell labeling owing to its fast reaction kinetics and high chemo-selectivity.^44–47^ The reaction proceeds readily in aqueous solvent in the absence of catalysts, forming highly stable adducts with the sole by-product being molecular nitrogen. In previous work, we developed the first eukaryotic compatible tetrazine ncAA (Tet-v3.0) GCE encoding system and demonstrated that these Tet-proteins can react rapidly and selectively with cyclopropane-fused, strained *trans*-cyclooctene (sTCO) reagents with rates as high as 8 × 10^4^ M^−1^ s^−1^.^48–50^

Here, we develop a GCE system with faster labeling rates and shorter linkage between protein and spin label, enabling SDSL-EPR studies of proteins directly within mammalian cells. We generated a family of small tetrazine amino acids, which we call Tet-v4.0 (Tet4), with the reactive tetrazine moiety appended directly to the amino acid β-carbon. We then evolved orthogonal aminoacyl tRNA synthetase (RS)/tRNA_CUA_ pairs capable of site-specifically incorporating Tet4 into proteins using both prokaryotic and eukaryotic expression systems. The resulting Tet4-incorporated proteins were found to react ~ 2‒15-fold faster with sTCO reagents than our previously reported tetrazine ncAAs, generating the first site-specific bioorthogonal protein labeling reaction with rates exceeding 10^6^ M^−1^ s^−1^.

We exploited the remarkable kinetics of the Tet4/sTCO reaction for SDSL-EPR by synthesizing three new sTCO‒nitroxide spin labels, including a *gem*-diethyl-substituted spin label that provided increased stability toward reduction in the cellular environment. These spin labels reacted rapidly and specifically with Tet4-incorporated proteins *in vitro*, in whole-cell lysates, and within live human embryonic kidney (HEK293T) cells. Quantitative labeling in HEK293T cells was achieved in minutes with sub-micromolar concentrations of spin label added to the growth medium. DEER on doubly spin-labeled Tet4 constructs of green fluorescent protein (GFP) and maltose-binding protein (MBP) expressed, labeled, and measured directly within HEK293T cells agree well with measurements from their *in vitro* purified counterparts. We further demonstrate that DEER can resolve the maltose-dependent open and closed conformations of Tet4-labeled MBP *in vitro*, and that it can identify the conformational state of a binding-impaired mutant of MBP directly within cultured mammalian cells.

## Results and Discussion

### Tet-v4.0 ncAA and sTCO-spin label design and synthesis

To achieve highly efficient spin-labeling of expressed proteins in live cells, we sought to develop an encodable, bioorthogonal system with the fastest possible reaction kinetics. Moreover, we sought to limit the size and flexibility of the resulting spin-labeled side chains, as spin labels with long and flexible linkers are known to broaden the distance distributions obtained with DEER spectroscopy and negatively impact structural interpretation. In previous work, we demonstrated that tetrazine amino acids can be site-specifically encoded into proteins where they are highly stable and show specific, rapid, and irreversible reactivity with sTCO reagents.^48,49,51^ We therefore chose to build on this initial work to design the Tet4 family of ncAAs (Figure 1A), in which the reactive tetrazine group is directly appended to the amino acid β-carbon. We generated three ncAAs—differing only in the functional group at the tetrazine 6-position—that we predicted would be stable in cells and display fast reaction kinetics with sTCO reagents. 6-methyl and 6-phenyl groups were chosen for their stability over a 6-position hydrogen, and the electron-withdrawing pyridyl variant was additionally included as it was predicted to display increased IEDDA reaction rates.^44,52^ All Tet4 ncAAs were synthesized from a common Boc-protected β-cyano alanine intermediate, generated in 4 steps from serine (Scheme S1). Substituted Tet4 ncAAs were obtained in 30-55% overall yield via nickel triflate-catalyzed nitrile coupling, using commercially available nitriles, followed by oxidation and deprotection.

**Figure 1.**
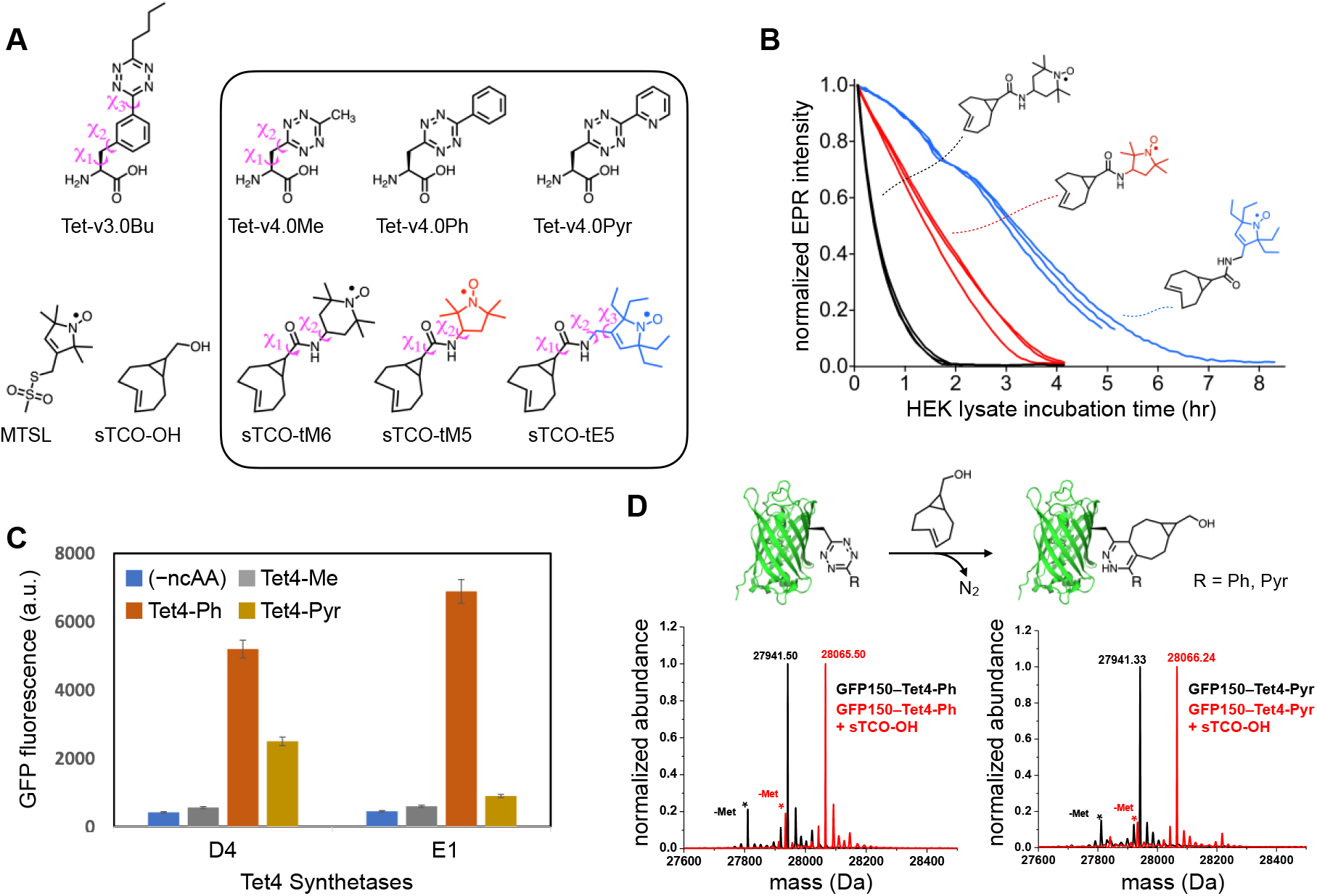
Tet-v4.0 ncAAs and sTCO-nitroxides: Spin-label Stability, Genetic Encoding, and Reactivity. (A) Tetrazine amino acid and sTCO-spin label structures. Rotatable side-chain dihedral angles preceding the tetrazine are illustrated for Tet-v3.0Bu and Tet-v4.0Me, as well as for sTCO-spin labels (B) Stability of sTCO-nitroxides (12 μM) in diluted HEK293T cytosolic extract at room temperature. EPR signal intensity was measured as the peak-to-trough amplitude of the X-band CW EPR spectrum. Each trace is an individual time course from triplicate measurements for each of the three sTCO-spin labels. (C) Tet4 incorporation efficiency measured with GFP fluorescence of *E. coli* co-transformed with GFP150‒TAG and evolved Mb-Pyl-tRNA/RS pairs D4 or E1 in the presence and absence of Tet4 ncAAs (0.5 mM). (D) Reactivity of GFP150‒Tet4-Ph/Pyr monitored by ESI-Q-TOF mass spectrometry. Schematic representation of the IEDDA reaction between GFP‒Tet4 and sTCO-OH (*top*). Deconvoluted mass spectra of purified GFP150‒Tet4-Ph (*bottom left*) and GFP150‒Tet4-Pyr (*bottom right*) before (black) and after (red) reaction with sTCO-OH. Cal. Mass of GFP‒wt: 27827.02 Da (avg); GFP150‒Tet4-Ph (observed: 27941.5 Da, expected: 27940.1Da); GFP150‒Tet4-Ph + sTCO-OH (observed: 28065.5 Da, expected: 28064.1 Da); GFP150‒Tet4-Pyr (observed: 27941.3 Da, expected: 27941.1Da); GFP150‒Tet4-Pyr + sTCO-OH (observed: 28066.2 Da, expected: 28065.1 Da). Asterisks (*) mark peaks corresponding to the loss of N-terminal methionine. Low intensity peaks at higher mass are sodium and potassium adducts.

As expected, Tet4 ncAAs reacted readily with sTCO reagents in aqueous buffer. Reaction rates with sTCO–alcohol (sTCO-OH)—determined by monitoring the loss of tetrazine absorbance at 270 nm with stopped-flow absorbance spectroscopy—yielded second-order rate constants of 1.1, 2.4, and 8.6 × 10^4^ M^−1^ s^−1^ for methyl-, phenyl-, and pyridyl-substituted Tet4 ncAAs, respectively (Table 1 and Figure S1). As predicted, the strongly electron-withdrawing pyridyl group of Tet4-Pyr yielded an approximately 4-fold increase in reaction rate relative to the 6-methyl and 6-phenyl Tet4 ncAAs, as well as to the best-performing Tet-v3.0 ncAA, Tet-v3.0-butyl (Tet3-Bu) (Table 1).

**Table 1.** Second-order reaction rate constants (k_2_) of tetrazine ncAAs with sTCO-OH

| Tetrazine Amino Acid | Reaction Rate ($\times 10^4 \text{ M}^{-1}\text{s}^{-1}$)<sup>†</sup> | |
| --- | --- | --- |
|  | Free amino acid | GFP-incorporated (TAG site) |
| Tet-v3.0Bu | $2.1 \pm 0.06$ | $7.8 \pm 0.30$ (N150) |
| Tet-v4.0Me | $1.1 \pm 0.14$ | --- |
| Tet-v4.0Ph | $2.4 \pm 0.20$ | $22 \pm 1.3$ (N150)<br>$20 \pm 1.1$ (L222) |
| Tet-v4.0Pyr | $8.6 \pm 1.2$ | $120 \pm 7.3$ (N150) |

<sup>†</sup> Second-order rate constants ( $k_2$ ) were obtained from plots of measured, pseudo-first order rate constants ( $k'$ ) against sTCO-OH concentration (see Figures S1 and S6).

In addition to the extraordinarily fast and selective reaction kinetics with tetrazines, sTCO reagents have also demonstrated good bioavailability while showing minimal degradation during the in-cell reaction timescale.^53,54^ To exploit these properties for use in SDSL-EPR, we synthesized three new sTCO-nitroxide spin labels (Figure 1A and Scheme S3). Aiming to minimize the linker length of the spin labels, we directly coupled sTCO-carboxylic acid to amino-functionalized nitroxides in which the amino group was situated as close to the nitroxide ring as possible (Scheme S3). We chose to explore three different nitroxide head groups: the 6-membered “TEMPO” nitroxide (tM6), and two 5-membered pyrroline/pyrrolidine (PROXYL) groups—one having the standard tetramethyl groups flanking the nitroxide (tM5) and the other containing tetraethyl substitutions (tE5). The tetraethyl—also known as *gem*-diethyl—substitutions have been shown to protect the nitroxide from chemical reduction to the EPR-silent hydroxylamine, which can occur in reducing environments such as those inside cells.^13,42,43,55–58^ Indeed, incubation of these sTCO-nitroxides in cytoplasmic extract from HEK293T cells revealed a strongly protective effect of the tetraethyl substitutions, with a 2-fold prolonged half-life of sTCO-tE5 compared with tetramethyl-substituted sTCO-tM5, and a nearly 8-fold improvement in comparison to the 6-membered ring sTCO-tM6 (Figure 1B).

### Synthetase selection for GCE with Tet-v4.0 ncAAs

To engineer a system for the genetic incorporation of Tet4 that would be compatible with both prokaryotic and eukaryotic expression systems, we focused on the pyrrolysyl-tRNA_CUA_/amino-acyl-tRNA synthetase pair (Pyl-tRNA/RS) from *Methanosarcina barkeri* (Mb), as this system demonstrated efficient encoding of our previously developed Tet-v3.0 family of ncAAs.^49^ Two Mb-Pyl-RS libraries—with 5 sites mutated to all 20 amino acids (3 × 10^6^ total variants)—were evaluated for the ability to accommodate Tet4-Me and Tet4-Ph (See SI for details). Standard life/death selection procedures involving two rounds of positive selection in the presence of 0.5 mM Tet4 ncAA and negative selection in the absence of ncAA were performed. The surviving synthetase variants were further evaluated for the ability to suppress an amber codon substituted at residue N150 of green fluorescent protein (GFP150-TAG). In total, four unique synthetases were identified that could express full-length GFP in the presence—but not in the absence—of Tet4-Ph, with two synthetase variants, D4 and E1, demonstrating high-fidelity incorporation and good yields for GFP150-TAG (~ 55-60 mg/L) (Figure 1C and Figure S2). Our screen yielded no synthetases capable of efficiently encoding Tet4-Me.

To incorporate Tet4-Pyr, we performed permissibility tests using the same four unique synthetases identified in our Tet4-Ph screen and found that only synthetase D4 was able to selectively incorporate Tet4-Pyr into GFP150-TAG with reasonable yields (~ 40 mg/L) (Figure 1C, Figure S3). The site-specific incorporation efficiency and fidelity of the E1 synthetase (for Tet4-Ph) and D4 synthetase (for Tet4-Pyr) were further verified by mass spectrometry of C-terminally 6×His-tagged GFP150-TAG constructs purified from *E. coli*. Deconvolutional analysis of the ESI-Q-TOF mass spectra yielded the expected masses confirming substitution of the native asparagine with Tet4-Ph or Tet4-Pyr, respectively (Figure 1D and SI Table S1). These results collectively demonstrate that evolved synthetases E1 and D4 can efficiently encode Tet4-Ph and Tet4-Pyr ncAAs, respectively, in *E. coli.* We therefore employed these two synthetases for all subsequent work.

### Reactivity of protein-encoded Tet-v4.0

To test the accessibility and reactivity of Tet4 ncAAs toward sTCO reagents once incorporated into proteins, we incubated purified samples of GFP150‒Tet4-Ph/Pyr with sTCO-OH and monitored the labeling reaction by mass spectrometry. Analysis of the ESI-Q-TOF data revealed apparently quantitative labeling as evidenced by mass increases of 124 Da for both GFP150‒Tet4 proteins, consistent with the expected sTCO-OH cycloaddition product and concomitant loss of N_2_ (Figure 1D and Table S1). We also verified labeling of GFP‒Tet4-Ph/Pyr using a more general SDS-PAGE mobility shift assay based on reaction with PEGylated sTCO reagents (sTCO-PEG5k/10k).^51^ GFP150‒Tet4-Ph displayed a near complete mass shift of ~ 5 kDa upon 10-minute incubation with 10-fold excess sTCO-PEG5k, indicating that virtually all protein contained the tetrazine and was quantitatively conjugated with sTCO-PEG5k (Figure S4A). A similar shift was observed for GFP150‒Tet4-Pyr but with a small percentage of un-reacted protein remaining, suggesting that some full-length GFP either did not contain Tet4-Pyr at residue 150—most likely from insertion of a canonical amino acid by way of near-cognate suppression^59^—or were otherwise unreactive, for example, due to tetrazine degradation. However, unreacted or degraded proteins were not detected by mass spectrometry, suggesting their abundance is low. We suspect that further synthetase optimization through more rigorous selections in the presence of Tet4-Pyr would generate a more robust synthetase and eliminate potential problems arising from near-cognate suppression.

DEER studies on proteins lacking intrinsic spin centers generally require the introduction of spin labels at two sites in the primary sequence. To verify that we could successfully encode Tet4 at two sites, we generated GFP constructs containing dual amber codon (TAG) substitutions at residues N150 and L222. Mass spectral analysis of purified GFP150/222‒Tet4-Ph confirmed the correct mass for doubly Tet4-Ph-incorporated GFP at residues 150 and 222, and subsequent reaction with sTCO-tE5 spin label revealed a mass increase of 690 Da, corresponding to the addition of two sTCO-tE5 spin labels minus two equivalents of molecular nitrogen (Figure S5A and Table S1). Dual labeling of GFP150/222‒Tet4-Ph was additionally confirmed by SDS-PAGE gel-band shift after reaction with sTCO-PEG5k (Figure S4A). Together these experiments demonstrate near-quantitative reactivity of singly and doubly Tet4-incorporated proteins.

Next, we determined the IEDDA reaction rates of Tet4 ncAAs incorporated at residue 150 of GFP. Tetrazine amino acids are known to partially quench GFP fluorescence when incorporated near the protein chromophore.^48,49,60^ Subsequent reaction of the tetrazine to form the 1,4-dihydropyridazine product removes this quenching effect and provides a sensitive measure of the reaction progress. We measured the time-dependent dequenching (increase in fluorescence) of purified GFP150–Tet4-Ph/Pyr upon reaction with sTCO-OH. Plots of pseudo-first order rate constants (*k*ʹ) against sTCO-OH concentration revealed 2^nd^-order rate constants (*k*_2_) of 2.2 × 10^5^ and 1.2 × 10^6^ M^−1^ s^−1^ for phenyl and pyridyl variants of Tet4, respectively (Table 1 and Figure S6). These represent the fastest reaction rates reported to date for genetically encoded tetrazine amino acids***Table 1***. We then measured rate constants (*k*_2_) for all three sTCO-nitroxides (Figure 1A) in reaction with GFP150‒Tet4-Ph. The observed reaction rates for sTCO-tM6, sTCO-tM5, and sTCO-tE5 were ~ 2‒4 × 10^5^ M^−1^ s^−1^, consistent with rates measured for sTCO-OH (Table 2 and Figure S7). The magnitude of these reaction rates reveal that spin-labeling is complete in seconds to minutes, even at sub-micromolar concentrations, and suggest that rapid and complete labeling of dilute protein solutions should be possible using stoichiometric amounts of sTCO-spin label.

**Table 2.** Reaction rates of GFP150-Tet-v4.0Ph with sTCO-nitroxides

| sTCO-Label | Reaction Rate ($\times 10^4 \text{ M}^{-1}\text{s}^{-1}$)<sup>†</sup> |
| --- | --- |
| sTCO-tM6 | $23 \pm 2.0$ |
| sTCO-tM5 | $24 \pm 1.3$ |
| sTCO-tE5 | $43 \pm 1.8$ |
| sTCO-OH | $22 \pm 1.3$ |

<sup>†</sup> Second-order rate constants ( $k_2$ ) were obtained from plots of the measured, pseudo-first order rate constants ( $k'$ ) against sTCO-OH or sTCO-nitroxide concentration (see Figures S6 and S7).

As we previously described, tetrazine amino acids incorporated into proteins react approximately three times faster with sTCO reagents than they do as free amino acids, presumably owing to the greater hydrophobicity of the protein surface relative to buffer.^48,49,61,62^ Our results on GFP150‒ Tet4 indicate a much larger rate increase of roughly 10-fold between protein-incorporated and free Tet4 (Table 1 and Figures S1 and S6). We hypothesize that the increased relative rate for the shorter Tet4 ncAAs compared to earlier Tet-v2.0 and Tet-v3.0 ncAAs could be a general effect arising from the closer proximity of the tetrazine group to the hydrophobic environment of the protein surface. However, it is also possible that the increased difference in relative rate observed for Tet4 stems from specific interactions dictated by the local environment. To test for site-specific effects on the Tet4 reaction rate, Tet4-Ph was encoded at position L222 of GFP which, in contrast to the positively charged environment surrounding GFP150, has a largely neutral electrostatic environment (Figure S8). Reaction of GFP222‒Tet4-Ph with varying concentrations of sTCO-OH yielded a 2^nd^-order rate constant of 2.0 × 10^5^ M^−1^ s^−1^, very similar to that of GFP150‒Tet4-Ph (2.2 × 10^5^ M^−1^ s^−1^), suggesting that local electrostatics do not play a significant role in the rate enhancements observed for protein-incorporated Tet4 ncAAs (Table 1). Taken together, our results demonstrate that the smaller Tet4 ncAAs can be efficiently encoded into proteins expressed in *E. coli*, either at a single site of interest or at two protein sites, where they can be selectively and quantitatively spin-labeled with sTCO-nitroxides. Moreover, the spin-labeling reaction proceeds with unprecedented speed, an important prerequisite for *in situ* labeling of proteins in mammalian cells.

### *In vitro* EPR and DEER

To investigate the utility of Tet4 incorporation for site-directed spin-labeling, we chose maltose binding protein (MBP) as a model. MBP undergoes a well-characterized clamshell closure that is stabilized by the binding of maltose and other maltodextrin ligands. The structures and energetics of this conformational change have been subject to extensive study using numerous biophysical techniques, including DEER,^63,64^ NMR,^65,66^ FRET,^67–70^ and X-ray crystallography.^71,72^ We selected four residues in the MBP sequence (S211, E278, K295, and E322) for mutation either to cysteine— for labeling with the thiol-specific spin label MTSL—or to Tet4 ncAA for labeling with sTCO-spin labels. In addition, we generated double-mutants with site-pairs 211/295 and 278/322 to probe the maltose-dependent conformational change of MBP using DEER (Figure 2A). Owing to the superior expression yields, we chose the Tet4-Ph ncAA incorporated with synthetase Mb-PylRS E1 for use in all spin-labeling studies.

**Figure 2.**
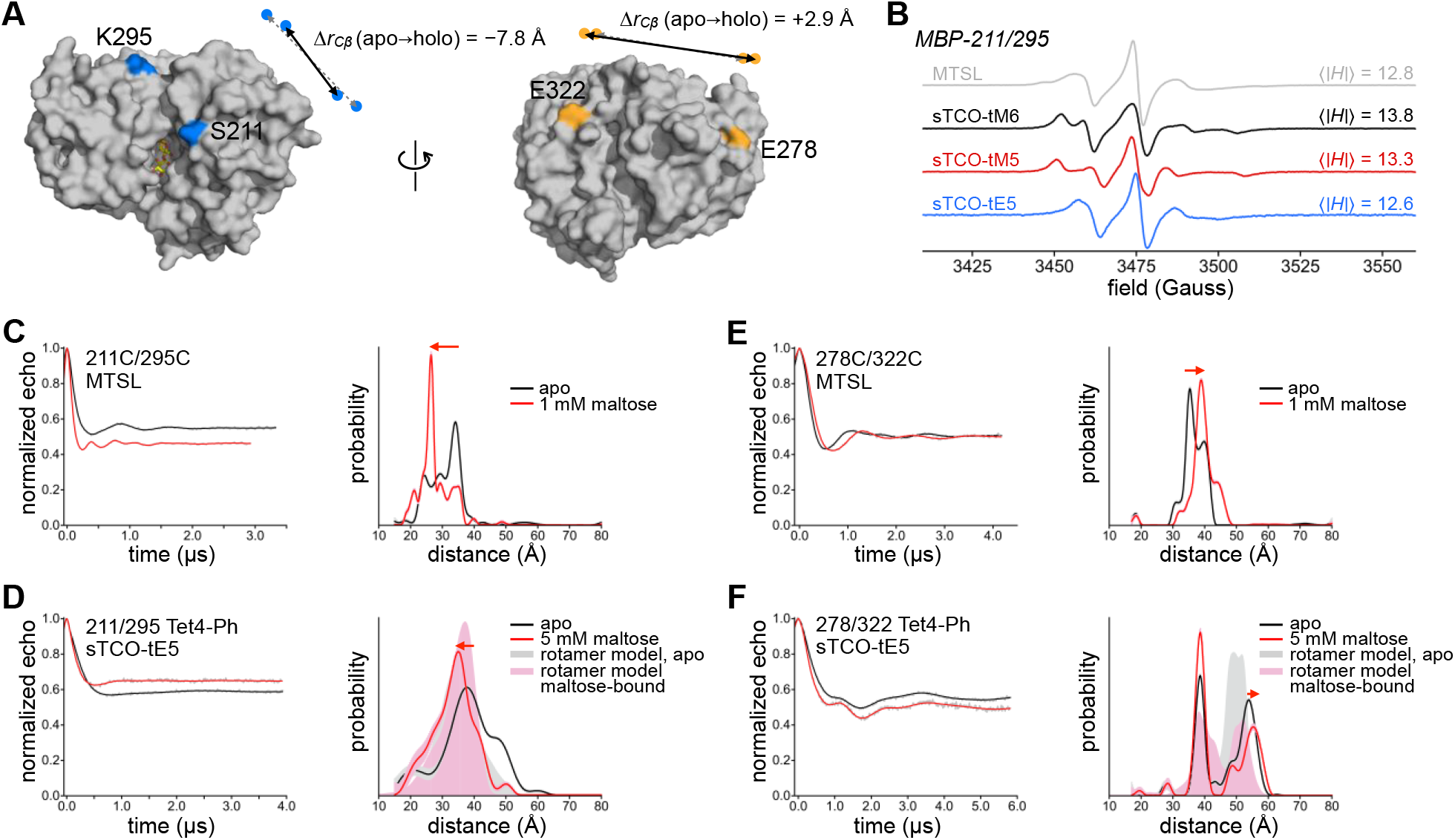
Tet-v4.0 SDSL-EPR and DEER on Maltose Binding Protein. (A) Surface rendering of maltose-bound MBP (1anf) with labeling sites indicated in color. For doubly-labeled constructs 211/295 (blue) and 278/322 (orange), the change in C_β_ distance between apo (dashed gray arrows) and holo (solid black arrows) conformations is indicated. (B) Room temperature X-band CW EPR spectra of purified MBP 211/295-Cys spin-labeled with MTSL (gray) and MBP 211/295-Tet4-Ph spin-labeled with sTCO-tM6 (black), -tM5 (red), or -tE5 (blue). (C‒F) Background-corrected DEER time traces and distance distributions for doubly spin-labeled MBP constructs 211/295 and 278/322. Maltose-free data are shown in black and data from samples recorded in the presence of 1 mM or 5 mM maltose are shown in red. Cysteine mutant constructs (C,E) were labeled with MTSL and Tet4-Ph constructs (D,F) were labeled with sTCO-tE5. Shaded distributions are predictions from *in silico* rotameric modeling with the chiLife software package in Python based on the available structures of apo (pdb 1omp; gray) and maltose-bound (pdb 1anf; pink) MBP.

Single and double cysteine and Tet4-Ph mutants of MBP were purified from *E. coli* and labeled with MTSL or sTCO-labels, respectively. Reaction of doubly Tet4-Ph-incorporated MBP constructs with sTCO-tE5 showed quantitative dual labeling by mass spectroscopic analysis (Figure S5B and Table S1). Likewise, reaction with sTCO-PEG10k showed two-site labeling efficiencies of 85-90% by SDS-PAGE mobility shift assay (Figure S4B and S9). In corroboration, labeling efficiencies of sTCO-nitroxides—as determined by double integration of the continuous-wave (CW) EPR spectra—were estimated to be ≥ 75% for all MBP‒Tet4-Ph sites studied (Figure S10). CW EPR spectra recorded at room temperature revealed that the side chains of MBP‒Tet4-Ph proteins labeled with sTCO-tM6 or -tM5 are significantly more rigid than are cysteines labeled with MTSL at equivalent residues, as evidenced by their larger spectral widths and correspondingly larger absolute-value first moments, 〈|Δ*H*|〉 (Figure 2B and Figure S11).^73^ This reduced rotational mobility is not surprising given the relatively bulky macrocyclic sidechain of spin-labeled Tet4 and the small number of rotatable bonds. MBP‒Tet4-Ph sites labeled with sTCO-tE5 display more mobile EPR spectra, similar to MTSL-labeled MBP (Figure 2B). We attribute this increased mobility relative to sTCO-tM6 and -tM5 spin labels to the additional methylene group—and hence, the extra rotatable bond—between the sTCO and pyrroline rings of sTCO-tE5 stemming from the amino-tE5 starting material used in our synthesis (Scheme S3).^58^

Next, we examined the ability of doubly spin-labeled MBP‒Tet4 constructs to report intramolecular distances and ligand-dependent conformational changes in MBP with DEER spectroscopy. Crystal structures of apo and maltose-bound MBP indicate a decrease in C_β_‒C_β_ distance between residues S211 and K295 of ~ 8 Å upon binding of maltose and closure of the clamshell. Conversely, the C_β_‒C_β_ distance between E278 and E322—located on the opposite surface of MBP (backside of the clamshell)—increases slightly (~ 3 Å) in the maltose-bound structure (Figure 2A). Indeed, 4-pulse DEER on MTSL-labeled MBP211C/295C clearly revealed the expected decrease in intramolecular distance in the presence of 1 mM maltose relative to the maltose-free sample (Figure 2C). Likewise, DEER revealed the small increase in distance expected upon maltose binding for MBP278C/322C labeled with MTSL (Figure 2E). To test if our Tet4-based spin-labeling system was capable of discerning the maltose-driven clamshell conformational change, we recorded DEER on doubly Tet4-Ph-encoded MBP constructs spin-labeled with sTCO-nitroxides, both in the presence and in the absence of maltose (Figure 2D,F and Figure S12). Distance distributions obtained for apo and maltose-bound MBP211/295‒Tet4-Ph labeled with sTCO-tE5 were in reasonable agreement with predictions from *in silico* rotameric modeling and displayed significantly broader distributions compared to the MTSL-labeled constructs (Figure 2D). Nevertheless, a clear shift toward shorter inter-spin distances was observed in the sample containing 5 mM maltose, consistent with the maltose-induced closure of MBP.

Surprisingly, DEER on sTCO-tE5-labeled MBP278/322‒Tet4-Ph revealed a pronounced bimodal distance distribution both in the presence and absence of maltose, with only one distance mode displaying the expected increase in inter-spin distance in response to maltose (Figure 2F). *In silico* spin label modeling using apo and maltose-bound MBP structures suggests that this multi-modality stems from restricted conformational sampling of the spin labels owing to the relatively bulky and rigid macrocyclic adduct being close to the protein surface. Our rotameric models suggest that steric clashes with neighboring residues may “trap” spin labels in distinct clusters of closely related conformations, giving rise to multimodal distance distributions (Figure S13). Indeed, multimodal distance distributions were also observed for tM6- and tM5-labeled MBP‒Tet4-Ph constructs (Figure S12). Altogether, these results show that DEER on *in vitro* spin-labeled Tet4-encoded proteins can reveal ligand-induced conformational changes giving rise to DEER distance changes as small as a few ångstroms; however, multimodal DEER distributions, likely arising from spin label rotameric restrictions, may be problematic at some protein sites and potentially complicate structural interpretation.

### Tet-v4.0 encoding and spin-labeling in mammalian cells

Next, we explored the compatibility of the Tet4 system with mammalian cells. Specifically, we examined the ability of Tet4 to be incorporated into proteins expressed in HEK293T cells, an immortalized human embryonic kidney cell line. As with previously reported Tet-v3.0,^49^ we found that Tet4-Ph and Tet4-Pyr ncAAs were well-tolerated by HEK293T cells when supplemented into the growth medium at concentrations up to 0.3 mM, whereas higher concentrations resulted in cytotoxicity (Figure S14). We cloned the genes for Tet4 E1 and D4 synthetases with a terminal nuclear export sequence into the eukaryotic expression Pyl-tRNA/RS vector pAcBac1, as described previously.^49^ Pyl-tRNA/RS pairs were then co-transfected with pAcBac1‒GFP150‒ TAG into HEK293T cells and incubated either in the absence of ncAA, or with various concentrations of Tet4 ncAAs, and suppression efficiencies were assessed using flow cytometry (Figure 3A and Figure S15-S18). Tet4-Ph and Tet4-Pyr were incorporated into GFP ~50% as well as the most efficient Tet-v3.0 system, Tet3-Bu. As with Tet4 incorporation into *E. coli* expressed proteins, the highest suppression efficiencies in HEK293T cells were obtained with Mb-PylRS synthetase E1 and Tet4-Ph ncAA. Selective incorporation and reactivity of GFP150‒Tet4 proteins were verified in whole-cell HEK293T lysates using sTCO-PEG5k-induced SDS-PAGE mobility shifts (Figure 3B and Figure S19), and in purified proteins by mass spectrometry (Figure S20 and Table S1).

**Figure 3.**
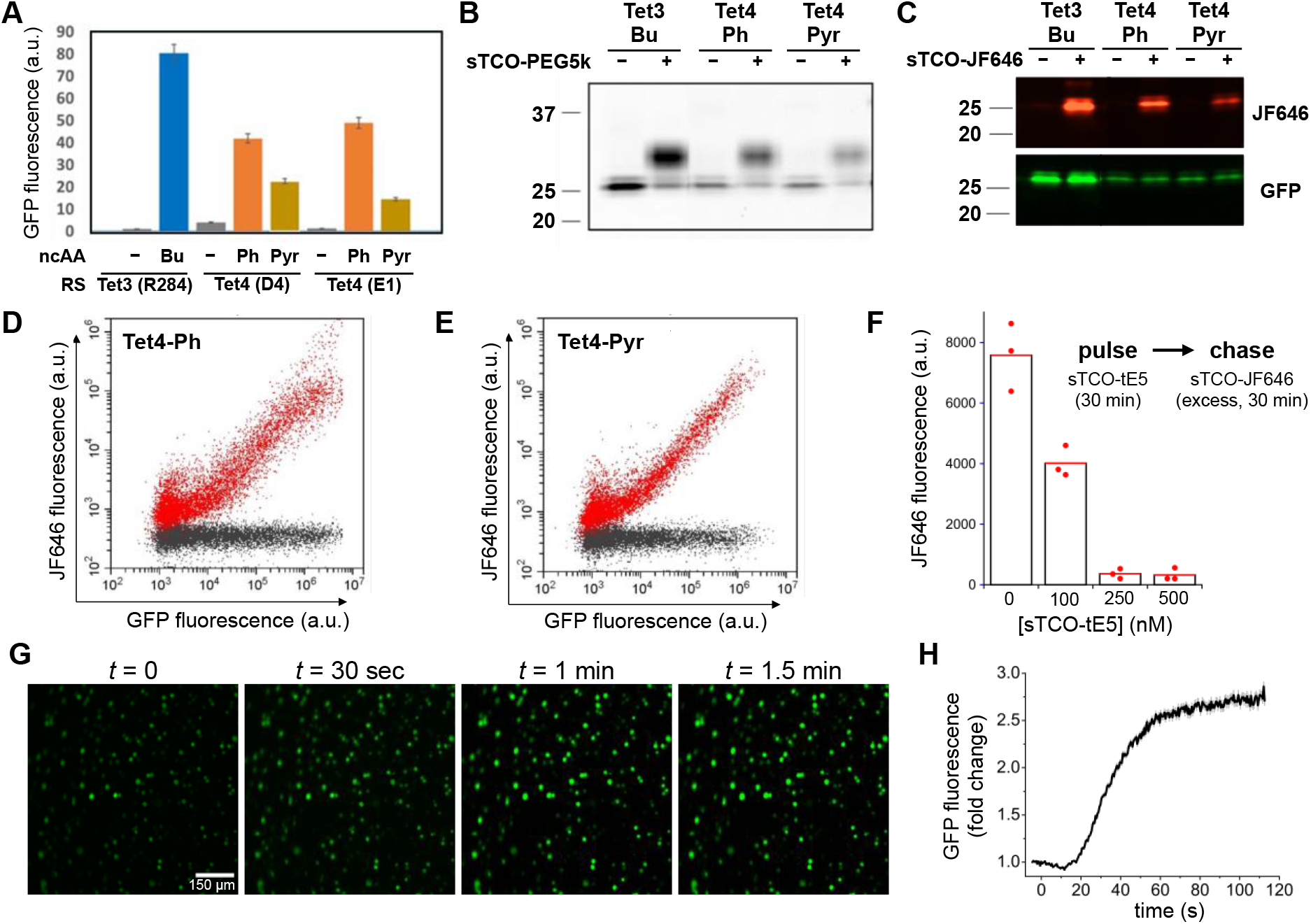
Encoding and reactivity of Tet-v4.0 in mammalian cells. (A) Suppression efficiency of GFP150-TAG with Tet4-Ph and Tet4-Pyr amino acids measured by flow cytometry of cultured HEK293T cells, in comparison with Tet3-Bu. (B) Reactivity of GFP150‒Tet4 in whole-cell HEK293T lysate verified by mobility shift of GFP detected by SDS-PAGE upon incubation with sTCO-PEG5k. (C) In-cell labeling of GFP150‒Tet4-Ph/Pyr in HEK293T cells with sTCO-JF646 as monitored by fluorescent imaging of SDS-PAGE gel. (D,E) Quantitative analysis of GFP150‒ Tet4-Ph (D) and GFP150‒Tet4-Pyr (E) labeling in live HEK293T cells with sTCO-JF646 using 2D single-cell fluorescent flow cytometry (red: 100 nM sTCO-JF646; black: 0.1% DMSO; 30 mins.). (F) Concentration dependence of in-cell spin-labeling of HEK293T cells expressing MBP322‒Tet4-Ph with sTCO-TEP assessed by pulse-chase with sTCO-JF646. MBP322‒Tet4-Ph molecules not labeled with sTCO-tE5 but subsequently labeled with excess sTCO-JF646 were quantified by in-gel fluorescence of whole-cell lysates. Points indicate individual experiments with mean JF646 fluorescence from triplicate experiments given by bars. (G,H) Live-HEK293T cell labeling of GFP150‒Tet4-Ph with sTCO-tE5 spin label. (G) Epifluorescence micrographs of HEK293T cells expressing GFP150‒Tet4-Ph at various times before and after perfusion with 1 μM sTCO-tE5. (H) Time course of GFP fluorescence increase of HEK293T cells expressing GFP150‒Tet4-Ph upon perfusion (at *t* = 0) with 1 μM sTCO-tE5 in DMEM + 10% FBS. Data are mean cell fluorescence divided by mean fluorescence at *t* = 0. Gray bars are mean ± standard error of the mean.

We then explored the prospect of *in situ* spin-labeling Tet4-incorporated proteins in living HEK293T cells. Several sTCO-functionalized fluorescent probes have previously been shown to permeate cell membranes and successfully label intracellular tetrazine-bearing proteins,^48,49,51^ and we reasoned that our small sTCO-nitroxides might also be cell-permeable. We first tested the reactivity of Tet4-encoded proteins in HEK293T cells by exposing cells expressing GFP150‒Tet4-Ph or Tet4-Pyr to the cell-permeable fluorophore sTCO-JF646.^51^ Cells were then washed, lysed, and analyzed by SDS-PAGE with fluorescence detection of both GFP and JF646 (Figure 3C and Figure S22). These results verified conjugation of JF646 exclusively at GFP, with undetectable labeling of off-target proteins. We further explored the reactivity of GFP150‒Tet4 in HEK293T cells by exposing cultured cells to sTCO-JF646 at two different concentrations—100 nM and 1 μM—for 30 minutes before quantifying GFP and JF646 fluorescence of individual cells using flow cytometry. The strong linear relationship between GFP and JF646 signals indicate that efficient, GFP-specific in-cell labeling occurred at 100 nM sTCO-JF646 with very little non-specific labeling (Figure 3D, E). At 1 μM sTCO-JF646, efficient GFP labeling was also achieved; however, significant background JF646 fluorescence was observed which suggests off-target association of the label with other cellular components (Figures S23 and S24). These experiments demonstrate that the fast reaction kinetics of Tet4-encoded proteins with sTCO reagents permits stoichiometric labeling in live eukaryotic cells, even at low concentrations of both label and protein.

Next, we exploited the in-cell labeling properties of sTCO-JF646 to assess the permeability and reactivity of our sTCO‒spin labels in live HEK293T cells using a pulse-chase assay. HEK293T cells expressing MBP322‒Tet4-Ph were incubated with various concentrations of the reduction-resistant spin label sTCO-tE5 for 30 minutes, after which the reaction was quenched with excess sTCO-JF646 (0.5 μM, 30 min, r.t.). Cells were then lysed and lysates analyzed using SDS-PAGE with fluorescence detection (Figure 3F and Figure S25). 100 nM sTCO-tE5 reduced the JF646 fluorescence 2-fold, indicating that roughly half of the available Tet4-Ph-incorporated MBP had been spin-labeled, whereas application of 250 nM sTCO-tE5 or higher led to complete spin-labeling of all available MBP‒Tet4 in the cell.

To further examine the time course of sTCO-tE5 labeling of HEK293T cells expressing GFP150‒ Tet4-Ph, we incubated the cells in culture medium containing 1 μM sTCO-tE5 while imaging GFP fluorescence with epifluorescence microscopy. As sTCO-tE5 entered cells and reacted with GFP150‒Tet4-Ph, the quenching effect of Tet4 on GFP fluorescence was relieved and an increase in fluorescence intensity was observed (Figure 3G,H). Under these conditions, the in-cell spin-labeling reaction appeared complete within 2 minutes. It should be noted that the labeling kinetics observed in this experiment are a product not only of the concentration-dependent IEDDA reaction rate, but also of the perfusion and mixing times, as well as the time required for sTCO-tE5 to diffuse across the cell membrane. These experiments suggest that intracellular Tet4-incorporated proteins can be quantitatively spin-labeled with our sTCO-nitroxides—directly within living cells—in a matter of minutes, using sub-micromolar concentrations of spin-label.

### In-cell EPR and DEER

To assess the possibility of detecting spin-labeled Tet4 proteins in mammalian cells by EPR, we incubated HEK293T cells expressing GFP150‒Tet4-Ph in medium supplemented with 200 nM sTCO-tE5 for 10 minutes, after which we removed the labeling medium, transferred the cells into a quartz EPR tube, and recorded CW EPR spectra of the pelleted cells at ambient temperature (Figure 4A, *top*). The EPR spectrum, recorded 1 hour after first exposure to spin label, reveals a slow-motional spectrum very similar to that of GFP150/222‒Tet4-Ph purified from *E. coli* and spin-labeled with sTCO-tE5 *in vitro* (Figure S26). This EPR signal was significantly reduced in spin-labeled cells in which the plasmid encoding the PylRS/tRNA pair was omitted from the transfection or where expression was carried out in the absence of Tet4-Ph ncAA (Figure 4A, *middle* and *bottom*). Similar synthetase-dependent in-cell CW EPR signals were also obtained from sTCO-tE5-labeled MBP295‒Tet4-Ph and MBP322‒Tet4-Ph (Figure S27). Although greatly reduced, there was still a notable slow-motion EPR spectrum in spin-labeled cells grown in the presence of Tet4-Ph, but lacking the plasmid encoding the PylRS/tRNA pair (see Figure 4A, *middle*). This likely stems from residual free Tet4-Ph ncAA in the cells being conjugated with sTCO-tE5 and highlights the importance of thoroughly washing free ncAA from the cells prior to exposure to sTCO reagents. Together these results demonstrate the ability of the reduction-resistant nitroxide sTCO-tE5 to site-specifically label proteins harboring Tet4 within living mammalian cells, generating EPR signals that persist for at least 1 hour at ambient temperatures with minimal background labeling.

**Figure 4.**
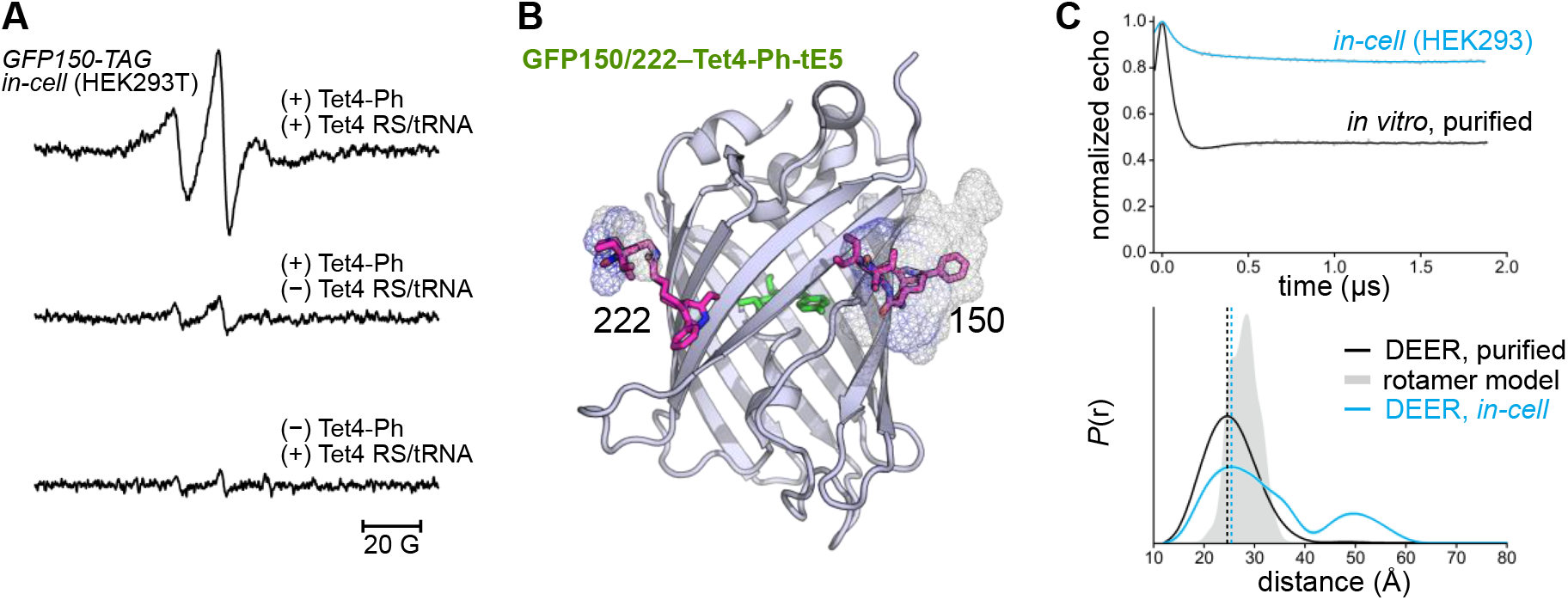
CW EPR and DEER in HEK293 cells. (A) Room temperature X-band EPR spectra of HEK293T cells transfected with GFP150-TAG and Tet4 Mb-PylRS/tRNA in the presence of 150 μM Tet4-Ph ncAA (*top*). Cells were spin-labeled with 200 nM sTCO-tE5 (10 min, r.t.) and spectra were recorded 1 h post-labeling. EPR spectra of labeled cells lacking either the Mb-PylRS/tRNA pair (*middle*) or Tet4-Ph ncAA (*bottom*) are displayed on identical scales. (B) Structural model of GFP150/222‒Tet4-Ph labeled with sTCO-tE5 displaying estimated rotameric distributions as mesh surface. Spin-labeled side chains were modeled with the chiLife package in Python. (C) 4-pulse DEER time traces (*top*) and distance distributions (*bottom*) for GFP150/222‒Tet4-Ph-tE5 recorded from *in vitro*-purified protein (black) and in intact HEK293T cells (blue). Error bands on the distributions estimated from LongDistances are shown but are similar to the linewidth. The simulated distance distribution from rotameric modeling is shown in gray shade.

To test whether accurate intramolecular protein distances could be measured by DEER in mammalian cells using the bioorthogonal Tet4/sTCO-nitroxide system, we generated GFP constructs for both bacterial and mammalian cell expression containing dual amber codon (TAG) substitutions at residues N150 and L222 (Figure 4B). We chose GFP for our initial in-cell DEER experiments because it is a small, well-structured protein devoid of large-scale conformational changes, thus allowing interpretation of the distance distribution without interference from conformational heterogeneity contributed by protein dynamics. Indeed, 4-pulse DEER of doubly Tet4-Ph-incorporated GFP purified from *E. coli* and subsequently spin-labeled with sTCO-tE5 yielded a mono-modal distance distribution centered at ~ 25 Å, in reasonable agreement with the simulated distance distribution predicted by rotameric modeling (Figure 4C). The large modulation depth (~ 50%) observed in the DEER time-domain data is again indicative of highly efficient spin label conjugation at both Tet4 sites.

Next, we expressed GFP150/222‒Tet4-Ph in HEK293T cells. After incubation with 250 nM sTCO-tE5 for 10 minutes, cells were washed once with PBS buffer before loading into an EPR tube and snap-freezing in liquid nitrogen for DEER measurement. The echo modulation obtained for the in-cell sample was significantly smaller than for the *in vitro*-purified sample, indicating a high percentage of singly spin-labeled molecules that do not contribute to echo modulation. These lone spins likely result from several sources: (1) chemical reduction of the nitroxides in the cell will increase the proportion of proteins having only one or zero active spin centers; (2) proteins truncated at the second amber codon will produce peptides having only one Tet4-Ph side chain available for spin-labeling; and (3) spin labels either free in solution or conjugated to residual tRNA-loaded or unincorporated Tet4-Ph will also result in a non-modulated DEER echo, decreasing the experimental modulation depth. Nevertheless, the resulting distance distribution obtained from intact HEK293T cells is very similar to the distance distribution obtained from purified protein, with distances of maximum probability—24.4 Å (*in vitro*) and 25.5 Å (in-cell)— differing by only 1.1 Å (Figure 4C). The in-cell DEER distribution does, however, contain a lower probability peak centered around 50 Å that is not present in the *in vitro* distribution. Given that the time-domain data were only collected out to ~ 1.8 μs, probabilities in this distance range are often unreliable and are likely attributable to uncertainties in the DEER background separation. However, we cannot rule out GFP dimerization, higher-order oligomerization, or partial unfolding in the cellular environment as sources of this distance component.

Having shown that accurate distance distributions could be obtained in HEK293T cells with the Tet4/sTCO-tE5 system, we asked whether we could use Tet4 incorporation and *in vivo* spin-labeling to determine the conformational state of a multi-state protein directly within the cellular environment. Here we turned again to MBP as a model protein; however, precise control of the cellular dextrin content—and, by extension, the MBP open-to-closed equilibrium—was not feasible. Moreover, FRET studies have suggested that an unidentified component of the HEK293T cytoplasm, perhaps glycogen, can bind MBP and bias the equilibrium toward the closed conformation.^68^ We therefore chose to introduce a mutation (W340A) within the maltose binding pocket of MBP known to lower the affinity for maltodextrin ligands by about two orders of magnitude (Figure 5A).^74^ We reasoned that this mutation would preclude the binding of endogenous dextrins or other small molecules from the HEK293T cytoplasm and drive the conformational equilibrium of MBP toward the fully open structure.

**Figure 5.**
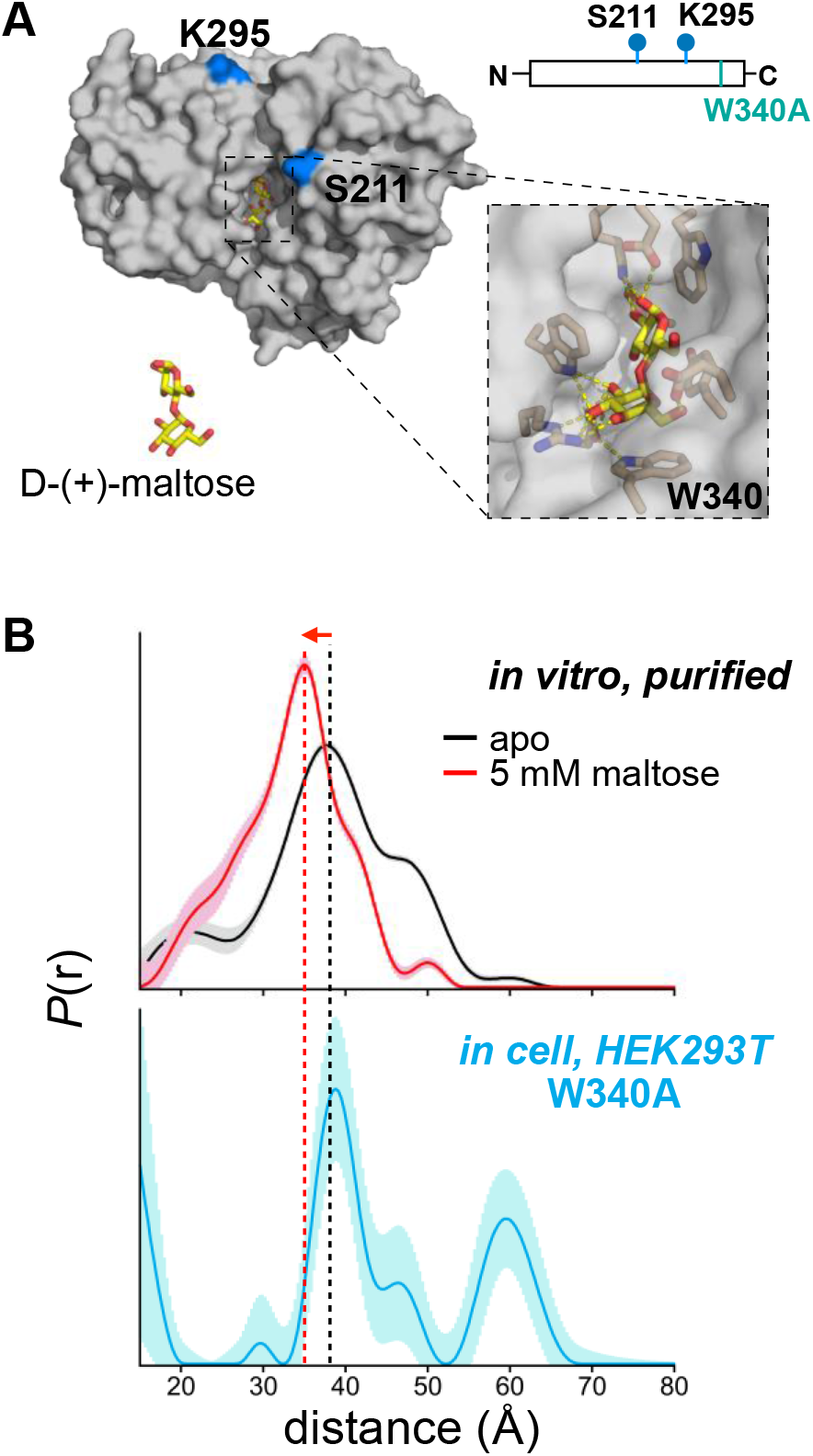
Conformation of MBP W340A in mammalian cells by DEER. (A) Structural model and schematic of the MBP construct used for in-cell DEER. Residues substituted with Tet4-Ph are indicated in blue. The maltose-binding pocket with Trp-340 highlighted is shown as zoomed inset. (B) Distance distributions obtained by DEER for MBP211/295‒Tet4-Ph-tE5 *in vitro* (*top*) and for MBP(340A)211/295‒Tet4-Ph-tE5 in HEK293T cells (*bottom*) with uncertainties represented as shaded error bands. Dashed lines centered at the maximum of the *in vitro* apo (black) and maltose-bound (red) distributions are shown for ease of comparison to the in-cell distribution.

Our *in vitro* DEER experiments on doubly sTCO-tE5-labeled MBP211/295 showed a clear shift in the distance distributions between apo and maltose-bound samples (see Figure 2D), and we therefore chose to introduce the 211TAG and 295TAG mutations—along with W340A—into our mammalian MBP expression construct for in-cell DEER measurements. MBP211/295‒TAG (W340A) was co-transfected with PylRS/tRNA E1 into HEK293T cells in the presence of 100 μM Tet4-Ph ncAA and the cells were spin-labeled with sTCO-tE5 in the same manner as described for the GFP DEER construct. The distance distribution obtained from 4-pulse DEER of the intact cells revealed a primary peak centered at ~ 39 Å, consistent with the apo (clamshell open) conformation of MBP measured from *in vitro*-purified protein (Figure 5B and Figure S28). While there was once again a long-distance peak at ~ 60 Å in the in-cell data, there was no component to the distance distribution that would suggest the presence of the closed conformation of MBP, in agreement with the expected effect of the W340A mutation. This result provides a proof-of-principle example that genetically encoded Tet4—in combination with *in situ* spin-labeling with sTCO-nitroxides— can identify protein conformations in their native environment within mammalian cells, without any protein purification or delivery steps. In-cell DEER using this system can potentially report on protein conformational equilibria, protein–protein interactions, and protein oligomerization in the native cellular environment and may prove useful in mechanistic studies of various cellular processes—including, but not limited to—receptor activation, developmental and cell-cycle-dependent mechanisms, and stress- or mutation-induced protein dysregulation. We note that all delivery of genetic components in this work was achieved by transient HEK293T cell transfection. We fully anticipate that more efficient methods of gene delivery, including stable cell line development and viral transduction, will greatly improve the sensitivity of in-cell DEER using the Tet4 system.

## Conclusions

In summary, we have developed a new family of tetrazine amino acids that are efficiently encoded into proteins using both prokaryotic and eukaryotic expression systems. The relatively small size and limited flexibility of the Tet-v4.0 family are desirable attributes for label-based distance measurement techniques such as DEER and FRET, where excessive probe length and flexibility can complicate structural interpretation and potentially obscure detection of protein conformational changes. We found that Tet4 ncAAs incorporated into proteins are site-specifically labeled by sTCO reagents with unprecedented speed. The reaction of GFP150‒Tet4-Pyr with sTCO-OH, for instance, was found to proceed with a second-order rate constant of 1.2 × 10^6^ M^−1^ s^−1^, which is—to our knowledge—the fastest genetically encoded bioorthogonal protein conjugation reaction reported to date. The tetrazine/sTCO IEDDA reaction is known to be accelerated by hydrophobic effects,^61,62^ and the rate enhancements we observe here for Tet4 relative to previously developed tetrazine-bearing ncAAs is likely due to the closer proximity of the Tet4 tetrazine group to the protein surface.

We harnessed the remarkable reaction kinetics of the Tet4/sTCO reaction for use in SDSL-EPR by developing a series of sTCO-functionalized nitroxides—all of which displayed rapid and specific spin-labeling of Tet4.0-incorporated GFP and MBP. DEER spectroscopy demonstrated that although distance distributions obtained from sTCO-spin-labeled proteins were significantly broadened compared with the gold standard cysteine-reactive spin label MTSL, Tet4-based spin labels were nevertheless able to report ligand-induced conformational rearrangements in MBP. We observed efficient incorporation and labeling at all six GFP and MBP residues studied— encompassing both *α*‒helical and β‒sheet structures—indicating that Tet4 should be broadly applicable for bioorthogonal spin-labeling of surface-exposed protein residues. Although multi-modal distance distributions were observed in several data sets that appeared to depend on the specific spin label and mutation site, we showed that rotameric modeling was able to predict this multi-modality in some cases, suggesting that computational modeling may be useful for selecting favorable sites for Tet4 incorporation. We anticipate that further explorations in spin label and tetrazine ncAA design will also help to understand the basis for multi-modal distance distributions and lead to improved ncAA/spin label pairs.

Lastly, we established rapid and selective spin-labeling of Tet4-incorporated proteins within live HEK293T cells using sub-micromolar concentrations of sTCO‒nitroxides applied directly to the cell culture medium. In-cell DEER measurements revealed inter-spin distance distributions of doubly labeled GFP and MBP Tet4 constructs and successfully reported on the in-cell conformation of a binding-site mutant of MBP. We anticipate that SDSL-EPR studies—and DEER spectroscopy in particular—using the Tet4/sTCO‒nitroxide system will help to uncover the nanoscale structures and conformational equilibria of proteins in their native cellular environments. In addition, this approach may prove useful for targeted dynamic nuclear polarization (DNP)-enhanced NMR studies and may help provide increased sensitivity of in-cell NMR measurements in the solid state.^75,76^

## Acknowledgements

This research was funded in part by the GCE4All Biomedical Technology Development and Dissemination Center supported by the National Institute of General Medical Science grant RM1-GM144227 as well as National Institutes of Health grant 1R01GM131168-01 (to R.A.M.) and National Science Foundation grant NSF-2054824 (to R.A.M). Additional National Institutes of Health and National Science Foundation research support came from NIGMS grant GM125753 and NSF grant CHE-2154302 (to S.S.), NIGMS grant R35GM145225 (to S.E.G), National Eye Institute grant R01EY010329 and NIGMS grant R01GM127325 (to W.N.Z.), NIGMS grant R01GM124310-01 and NSF grant CHE-1955349 (to A.R.), and NIH training grant T32-EY007031 (to E.G.B.E). We would like to thank Drs. Galen Flynn and Maxx Tessmer (University of Washington) for invaluable technical support and advice. Protein mass spectrometry was carried out at the Oregon State University Mass Spectrometry Center on the Waters Ion Mobility ToF Mass Spectrometer supported by NIH grant 1S10RR025628-01.

## Notes

### Competing Interest Statement

The authors have declared no competing interest.

### Summary of Updates

Updated the authorship order with the author information added below- Subhashis Jana,a⊥ Eric G. B. Evans,b c⊥ Hyo Sang Jang,a Shuyang Zhang,d Hui Zhang,d Andrzej Rajca,d Sharona E. Gordon,c William N. Zagotta,c Stefan Stoll,✽,b and Ryan A. Mehl ✽,a a Department of Biochemistry and Biophysics, Oregon State University, Corvallis, Oregon 97331, United States. b Department of Chemistry, University of Washington, Seattle, WA 98195, United States. c Department of Physiology & Biophysics, University of Washington, Seattle, WA 98195, United States. d Department of Chemistry, University of Nebraska, Lincoln, NE 68588-0304, United States. ⊥ Equal contributors ✽ To whom correspondence should be addressed

