## Supplementary Material for "Ultra-Fast Bioorthogonal Spin-Labeling and Distance Measurements in Mammalian Cells Using Small, Genetically Encoded Tetrazine Amino Acids"

### This PDF file includes:

Materials and Methods
Schemes S1–S3
Figures S1 to S28
Tables S1
$^{1}H$ and $^{13}C$ NMR spectra
ESI-MS spectrum

References 1–12

### Materials and Methods

#### General Synthetic Methods:

All purchased chemicals were used without further purification. 3-Amino-PROXYL was purchased from Toronto Research Chemicals and 3-aminomethyl-2,2,5,5-tetraethyl-1-pyrrolinyloxy free radical was synthesized according to the published procedure.[1] Anhydrous dichloromethane (DCM) and dimethyl sulfoxide were used after overnight stirring with calcium hydride and distillation under argon atmosphere. Thin layer chromatography (TLC) was performed on silica 60F-254 plates. The TLC spots of alkene were identified by potassium permanganate staining. Flash chromatographic purification was performed using silica gel 60 (230-400 mesh size). $^{1}$H NMR spectra were recorded on Bruker at 400MHz and 700 MHz and $^{13}$C NMR spectra were recorded at 175 MHz. The chemical shifts were shown in ppm and are referenced to the residual non-deuterated solvent peak $CDCl_3$ ($\delta$ =7.26 in $^{1}$H NMR, $\delta$ = 77.23 in $^{13}$C NMR), $CD_3OD$ ($\delta$ =3.31 in $^{1}$H NMR, $\delta$ = 49.2 in $^{13}$C NMR), $d_6$-DMSO ($\delta$ =2.5 in $^{1}$H NMR, $\delta$ = 39.5 in $^{13}$C NMR) as an internal standard. Splitting patterns of protons are designated as follows: s-singlet, d-doublet, t-triplet, q-quartet, m-multiplet, bs- broad singlet, dd- doublet of doublets.

#### Synthetic Procedure

***Tert-butyl (S)-(3-hydroxy-1-(methoxy(methyl)amino)-1-oxopropan-2-yl) carbamate*:**[2] To a solution of N-boc protected L-serine (2 gm, 9.7 mmol) in a dry DCM (50 mL) N, O dimethyl

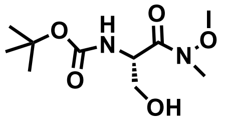

hydroxyl amine hydrochloride (1.4 gm, 10.7 mmol) was added, and the reaction mixture was stirred at -5 $^{o}$C (ice and salt mixture) under nitrogen atmosphere. Then the solution was treated with N-methyl morpholine (1.2 mL, 10.7 mmol) and 1-Ethyl-3-(3-dimethylaminopropyl) carbodiimide (2 gm, 10.7 mmol) was added in three portions over 30 minutes. After stirring at -5 $^{o}$C for 2 hours the reaction mixture was treated with 1.0 M aqueous HCl solution and immediately extracted with 200 mL dichloromethane. The organic layer was washed with saturated sodium bicarbonate, then brine solution and dried over $Na_2SO_4$. The solvent was removed by rotary vacuum and afforded white solid compound utilized for next step without purification (yield- 83%). $^{1}$H NMR ($CDCl_3$; 400 MHz) $\delta$ 5.64 (bs, 1H), 4.8 (bs, 1H), 3.84-3.79(m, 2H), 3.78 (s, 3H), 3.23 (s, 3H), 1.45 (s, 9H).

***Tert-butyl (S)-(1-(methoxy(methyl)amino)-3-(methylsulfonyl)-1-oxopropan-2-yl) carbamate* (4)** :[2] The N, C protected L-serine (1 gm, 4.1 mmol) was dissolved in a dry DCM under nitrogen

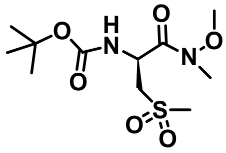

atmosphere, the temperature was reduced to 0 $^{o}$C, then mesyl chloride (380 μL, 4.92 mmol) and triethyl amine (680 μL, 4.92 mmol) were added. The reaction mixture was stirred until the starting material consumed as monitored by TLC (nearly an hour), then it was poured into water and DCM (50 ml each). The organic layer was washed with water two times, dried under sodium sulfate and then evaporated. The pure compound was isolated by silica gel flash column chromatography (40%

ethyl acetate in hexanes) as brown oil (Yield- 85%).$^1$H NMR ($CDCl_3$; 400 MHz) δ 5.34 (d, 1H), 4.95 (bs, 1H), 4.43-4.49 (m, 2H), 3.79 (s, 3H), 3.04 (s, 3H), 3.26 (s, 3H), 1.46 (s, 9H).

***(S)-2-((tert-butoxycarbonyl)amino)-3-cyanopropanoic acid*** **(3)**: To a solution of the N, C protected mesylated serine (1 gm, 3.1 mmol) in dry DMSO (20 ml), sodium cyanide (380 mg, 7.8 mmol) was added and stirrer for 24 hrs. at 55 °C. Next, 20 mL water was added to reaction mixture and extracted with ethyl acetate 3 times (20 mL each). The combined organic layer was washed

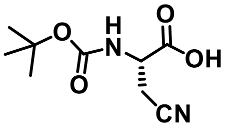

with brine solution 3 times to remove the DMSO, then dried under anhydrous sodium sulfate and evaporated. The synthesized N, C protected beta cyano-alanine was isolated by silica gel flash column chromatography (hexane: ethyl acetate, 3:1), with a yield of 65%. Next, the C-terminal Weinreb amide was deprotected using 1.5 eqv. lithium hydroxide in (tetrahydrofuran) THF- water (10:1) system at 0 °C. After completion of the reaction, the THF was removed by a rotary evaporator. Then 10 mL water was added to the reaction mixture and it was cooled to 0 °C, then acidified (pH 3 to 4) with diluted acetic acid. The reaction mixture was extracted with DCM and the combined organic layers were dried over anhydrous sodium sulfate. The N-boc protected beta cyano-alanine was isolated by silica gel flash column chromatography (10 % methanol in DCM) with a yield of 96%. $^1$H NMR (700 MHz, $CDCl_3$) δ 5.77 (bs, 1H), 4.44 (bs, 1H), 2.99 (d, 1H), 2.91 (d, 1H); 1.41 (s, 9H). $^{13}$C NMR (175MHz, $CDCl_3$- $CD_3OD$ mix.) δ 171.1, 155.3, 116.8, 81.1, 50.1, 28.3, 21.8.

***(S)-2-((tert-butoxycarbonyl)amino)-3-(6-methyl-1,2,4,5-tetrazin-3-yl)propanoic acid*** **(2a):** A flame dried, 15 mL heavy walled reaction tube was charged with boc-protected β-cyano L-alanine

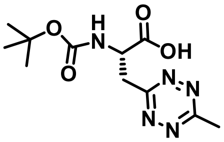

(200 mg, 0.934 mmol), nickel(II) trifluoromethanesulfonate ($Ni(OTf)_2$) (166 mg, 0.467 mmol) and acetonitrile (0.5 mL, 9.3 mmol) under argon atmosphere. Under argon anhydrous hydrazine (1.4 mL, 46.7 mmol) was added to the reaction mixture with stirring then the reaction vessel was purged with argon for 10 minutes. The sealed reaction mixture was heated to 35-37 °C with stirring for 32 hrs. The reaction mixture was cooled to room temperature, opened slowly, and 10 eqv. of aqueous 2 M sodium nitrite ($NaNO_2$) and 10 mL water were added with stirring. Next, the reaction mixture was washed with 20 mL ethyl acetate to remove the homocoupled product. The aqueous phase was acidified with 4 M HCl (pH~2) under ice cold conditions then was extracted with ethyl acetate (3 times). The combined organic layer was dried with sodium sulfate and concentrated under reduced pressure. Silica gel flash column chromatography (30% ethyl acetate in hexanes with 1% acetic acid) of the resin yielded 143 mg of **Boc-Tet4-Me** (0.51 mmol, 54 %) in the form of a pinkish red gummy material. $^1$H NMR (400MHz, $CDCl_3$) δ 5.62 (d, 1H), 4.54 (bs, 1H), 3.84 (dd, 2H), 3.07 (s, 3H), 1.47 (s, 9H). $^{13}$C NMR (175MHz, $CDCl_3$) δ 172.1, 167.7, 166.3, 155.4, 81.2, 50.2, 37.1, 28.2, 21.6.

***(S)-2-amino-3-(6-methyl-1,2,4,5-tetrazin-3-yl)propanoic acid trifluoro acetate salt* (1a):** In a dry round-bottom flask, boc-protected Tet4-Me amino acid (120 mg, 0.423 mmol) was dissolved in

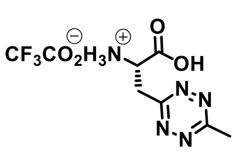

1:1 mixture of dry dichloromethane and trifluoroacetic acid (TFA) by volume (1.0 ml total) under argon. The reaction mixture was allowed to stir at room temperature for 1 hr. It was then concentrated under reduced pressure, and dissolved in DCM to drive off TFA. This process was repeated twice prior to drying completely under high vacuum affording a pinkish red color TFA salt of **Tet4-Me**, with quantitative yield. $^{1}$H NMR (700MHz, $CD_3OD$) δ 4.48 (bs, 1H), 3.98 (dd, 1H), 3.83 (dd, 1H), 3.04 (s, 3H). $^{13}$C NMR (175MHz, $CD_3OD$) δ 169.7, 167.7, 163.5, 51.9, 36.7, 21.3. ESI-MS calculated for $C_6H_{10}N_5O_2$ ($[M + H]^+$) 184.0829, found 184.0834.

***(S)-2-((tert-butoxycarbonyl)amino)-3-(6-phenyl-1,2,4,5-tetrazin-3-yl)propanoic acid* (2b):** In a flame dried, 15 mL heavy walled reaction tube containing boc-protected β-cyano L-alanine (120 mg, 0.560 mmol) was charged with nickel(II) trifluoromethanesulfonate ($Ni(OTf)_2$) (98 mg, 0.28

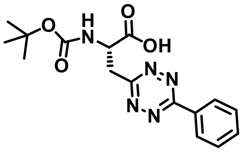

mmol) and benzonitrile (0.4 mL, 3.9 mmol) under an argon atmosphere. Then anhydrous hydrazine (1 mL, 28 mmol) and 0.3 mL ethanol (EtOH) were added to the reaction mixture and purged with argon for 10 minutes and the tube immediately sealed. The reaction mixture was heated to 42 $^{o}$C for 30 hrs. The reaction mixture was cooled to room temperature, opened slowly and 10 eqv. 2 M sodium nitrite aqueous solution and 10 mL water were added to the reaction mixture. Then, the reaction mixture was washed with ethyl acetate to remove the homocoupled product. The aqueous phase was acidified with 4 M HCl (pH~2) under ice cold conditions and subsequently extracted with ethyl acetate (3 times). The combined organic layer was dried with anhydrous sodium sulfate and concentrated under reduced pressure. Silica gel flash column chromatography (15 % ethyl acetate in hexanes with 1% acetic acid) yielded 62 mg of **Boc-Tet4-Ph** (0.18 mmol, 32 %) in the form of a reddish pink color gummy material. $^{1}$H NMR (400MHz, $CDCl_3$) δ 8.55 (d, 2H), 7.59-7.57 (m, 3H), 5.67 (d, 1H), 5.01 (bs, 1H), 3.96 (dd, 2H), 1.35 (s, 9H). $^{13}$C NMR (175MHz, $CDCl_3$) δ 174.8, 166.7, 164.6, 155.6, 133.1, 131.6, 129.5, 128.3, 80.9, 51.9, 37.4, 28.4.

***Hydrochloride salt (S)-2-amino-3-(6-phenyl-1,2,4,5-tetrazin-3-yl)propanoic acid* (1b):** In a dry round-bottom flask, boc-protected Tet4-Ph amino acid (80 mg, 0.23 mmol) in 3 mL ethyl acetate was charged with 1 mL dioxane (saturated with HCl gas) under an argon atmosphere. The reaction

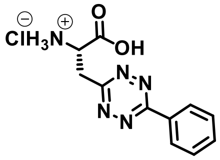

mixture was allowed to stir at room temperature for 2 hr. Then it was concentrated under reduced pressure, then dissolved in ethyl acetate. This process was repeated 2 times to remove excess HCl gas. Finally, 5 mL pentane was added and dried completely under high vacuum resulting in a solid radish-pink color chloride salt of **Tet4-Ph**, with quantitative yield (~ 97%). $^{1}$H NMR (700MHz, $CD_3OD$ ) δ 8.59 (d, 2H), 7.69 (t, 1H), 7.65 (t, 2H), 4.81 (dd, 1H), 4.07 (dd, 2H), 3.99 (dd, 2H). $^{13}$C NMR (175MHz, $CD_3OD$) δ 170.6, 167.2, 166.3, 134.2, 133.4, 130.6, 129.2, 52.2, 36.22. ESI-MS calculated for $C_{11}H_{12}N_5O_2$ ($[M + H]^+$) 246.0986, found 246.0994.

***(S)-2-((tert-butoxycarbonyl)amino)-3-(6-(pyridin-2-yl)-1,2,4,5-tetrazin-3-yl)propanoic acid*** **(2c):** Following the synthetic procedure for **2b**, the boc-protected β-cyano L-alanine (0.20 g, 0.93 mmol) and 2-pyridine carbonitrile (0.7 mL, 6.5 mmol) produced 0.145 g (0.42 mmol) of the title

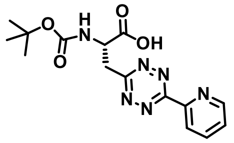

compound **Boc-Tet4-Pyr** as a pink color material with a yield 45%. $^{1}$H NMR (400MHz, $CDCl_3$) δ 8.96 (d, 1H), 8.68 (d, 1H), 8.05 (t, 1H), 7.62 (t, 1H), 5.71 (bs, 1H), 5.01 (bs, 1H), 4.03 (bs, 2H), 1.41 (s, 9H). $^{13}$C NMR (175MHz, $CDCl_3$) δ 176.4, 174.1, 167.9, 155.6, 150.5, 149.8, 138.2, 127.1, 124.4, 80.5, 52.2, 37.7, 28.4.

***Hydrochloride salt of (S)-2-amino-3-(6-(pyridin-2-yl)-1,2,4,5-tetrazin-3-yl) propanoic acid*** **(1c):** Following the boc-deprotection reaction for **1b,** the chloride salt of **Tet4-Pyr** was generated in a

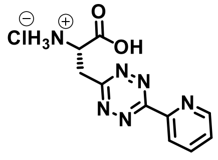

near quantitative yield (~ 97-98%). $^{1}$H NMR (700MHz, $CD_3OD$) δ 9.1 (d, 1H), 9.06 (d, 1H), 8.71 (t, 1H), 8.21 (t, 1H), 4.23(dd, 1H), 4.13(dd, 1H). $^{13}$C NMR (175MHz, $CD_3OD$) δ 170.3, 169.1, 162.5, 147.6, 147.1, 146.3, 130.7, 127.2, 52.1, 36.5. ESI-MS calculated for $C_{10}H_{11}N_6O_2$ ($[M + H]^+$) 247.0938, found 247.0932.

***4-Nitrophenyl sTCO*** **(6)**: Synthetic procedure Jang et al. and O'Brien et al.,[4, 5] $^{1}$H NMR (400MHz,

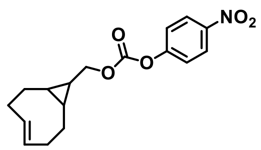

$CDCl_3$) *δ* 8.27 (2H, d), 7.37 (2H, d), 5.88-5.82 (1H, m), 5.18-5.14 (1H, m), 4.18 (2H, d), 2.43-2.39 (1H, m), 2.35-2.22 (3H, m), 1.96-1.90 (2H, m), 0.94-0.83 (1H, m), 0.69-0.64 (1H, m), 0.62-0.49 (3H, m).

***sTCO-amine*** **(7)**: In 3 mL of anhydrous dichloromethane, 100 mg of the 4-nitrophenyl sTCO (6) (0.315 mmol) and 100 µL (1.5 mmol) of ethylenediamine were added under an argon atmosphere.

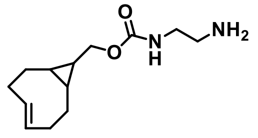

The reaction mixture was allowed to stir at room temperature for overnight. After that, the solvent was concentrated onto silica gel under reduced pressure and the product **7** (45 mg, 0.19 mmol) was purified by silica gel column chromatography (30-35% methanol in DCM). Yield 60%. $^{1}$H NMR (400MHz, $CDCl_3$) *δ* 5.83-5.75 (1H, m), 5.09-5.01 (1H, m), 3.85 (2H, d), 3.18 (2H, t), 2.77 (2H, t), 2.30-2.26 (1H, m), 2.21-2.12 (3H, m), 1.87-1.80 (2H, m), 0.79-0.74 (1H, m), 0.50-0.43 (2H, m), 0.37-0.34 (2H, m).

***sTCO-JF646*** **(8):** In a flame dried round-bottom flask, Dye JF-646-NHS (5 mg, 0.008 mmol) was dissolved in anhydrous DCM (2 mL). sTCO-amine **7** (3 mg, 0.012 mmol) then N,N-diisopropylethylamine (DIPEA) (20 µl, 0.15 mmol) was added to the solution and the reaction

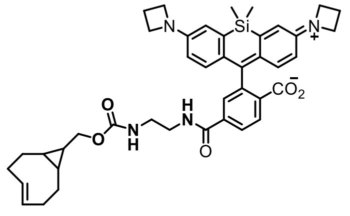

mixture was stirred overnight at room temperature. The solvent was concentrated onto silica gel under reduced pressure and purified the title compound **8** (4.5 mg, 0.006 µmol) by silica gel column chromatography (20% methanol in DCM). Yield 75%. $^{1}$H NMR (400MHz, $CD_3OD$) *δ* 8.023 (2H, d), 7.73 (1H, s), 6.74 (2H, d), 6.72

(1H, s), 6.70 (1H, s), 6.31 (2H, dd), 5.81-5.74 (1H, m), 5.09-5.02 (1H, m), 3.91 (8H, t), 3.81 (2H, d), 3.46 (2H, t), 2.39 (4H, q), 2.26 (1H, d), 2.19-2.15 (3H, m), 1.89-1.79 (2H, m), 0.89 (1H, t), 0.83-0.72 (1H, m), 0.63 (3H, s ), 0.55 (3H, s ), 0.51-0.42 (2H, m), 0.37-0.27 (2H, m).

***sTCO-PEG5000* (9):**[5] Amino-PEG5000 (130 mg, 0.026 mmol) and 4-nitrophenyl sTCO (6) (10 mg, 0.031 mmol) were dissolved in 3 mL of anhydrous DCM, then triethylamine (10 µL, 0.052 mmol) was added under an argon atmosphere. The reaction mixture was stirred at room temperature for 18 hours. The reaction mixture was concentrated onto silica gel under reduced pressure and purified by silica gel column chromatography (5% methanol in DCM) resulting in **9** (104 mg, yield 74%). $^1$H NMR (400MHz, $CD_3OD$) $\delta$ 5.90-5.82 (1H, m), 5.17-5.09 (1H, m), 3.91(2H, d, $J$ = 6.8 Hz), 3.82-3.79 (3H, m ), 3.63 (411H, bs), 3.54-3.50 (4H, m), 3.45 (2H, t, $J$ = 4.8 Hz), 3.35 (3H, s), 3.26 (2H, t, $J$ = 5.6 Hz), 3.16 (2H, q, $J$ = 7.6 Hz), 2.35 (1H, d, $J$ = 15.2 Hz), 2.27-2.15 (3H, m), 1.97-1.88 (2H, m), 0.95-0.84 (1H, m), 0.66-0.54 (2H, m), 0.48-0.41 (2H, m).

***(E)-bicyclo[6.1.0]non-4-ene-9-carboxylic acid* (10, sTCO-$CO_2$H):** Following the literature method[3], a continuous flow apparatus was used for photoisomerization with active removal of trans-isomer by silver nitrate impregnated silica gel. In a dry quartz flask, sCCO-$CO_2$H -cis isomer[4] (1.2 gm, 7.22 mmol) and methyl benzoate (2.2 mL, 18.06 mmol) were dissolved in 400 mL solvent (hexane:ether, 1:1). The flask was placed in a Raynot (R) reactor and connected via

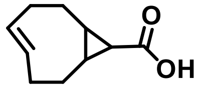

PTFE tubing to a column and FMI pump. The column was packed with dry silica (60 Å, 6 cm) and then silver impregnated silica (17 gm). Then the pump was turned on and the rate of circulation adjusted to 80 mL/min and 16 low pressure mercury lamps (2537 Å) turned on. The photolysis of the reaction mixture was continued for 7 hrs. The column was washed with additional 400 mL ether then silica was poured into a 500 mL erlenmeyer flask which was allowed to dry. Then the silica was stirred with saturated aqueous sodium chloride solution (150 mL) and methylene chloride (150 mL) for 15 minutes. After filtration of silica, the aqueous part was extracted with dichloromethane (3 times). The combined organic layers again washed with 100 mL water, dried with anhydrous sodium sulfate, and concentrated under reduced pressure to afford trans isomer–sTCO-$CO_2$H (**10**) as a white solid that was used without further purification. Yield 62%. $^1$H NMR (400MHz, $CDCl_3$) δ 5.91-5.83 (1H, m), 5.20-5.12 (1H, m), 2.40 (1H, d, $J$ = 4.8 Hz), 2.32-2.22 (3H, m), 2.03-1.92 (2H, m), 1.37-1.33 (1H, m), 1.28-1.21 (1H, m), 0.96-0.89 (2H, m), 0.65 (1H, q, $J$ = 12.8 Hz). $^{13}$C NMR (175MHz, $CD_3OD$) δ 178.8, 139.1, 132.7, 39.1, 34.4, 33.1, 28.5, 28.3, 28.3, 27.4.

***sTCO -2,2,5,5-tetramethyl-1-piperidinyloxy free radical* (sTCO-tM6, 11):** In dry DCM (3 mL), sTCO-$CO_2$H **10** (12 mg, 0.07 mmol), and 1-Ethyl-3-(3-dimethylaminopropyl) carbodiimide (EDC.HCl) (17 mg, 0.086 mmol) were added under an argon atmosphere and allowed to stir for 15 minutes under ice cold conditions. 4-Amino-TEMPO free radical (14 mg, 0.08 mmol) was

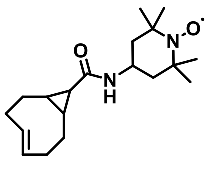

added followed by N,N-diisopropylethylamine (DIPEA) (10 µl, 0.4 mmol) to the reaction mixture. After 15 minutes, ice bath was removed and stirring was continued for another 16 hours at room temperature. The reaction was concentrated onto silica gel under reduced pressure and **11** (9 mg, 0.028 mmol) was purified by silica gel column chromatography (50% ethyl acetate in hexane). Yield 40%. ESI-MS calculated for $C_{19}H_{32}N_2O_2$ ([M + H]$^+$) 320.2464, found 320.2463.

***sTCO -2,2,5,5-tetramethyl-1-pyrrolidinyloxy free radical*** **(sTCO-tM5, 12):** Following the procedure for making **11**, 3-Amino-PROXYL was used to synthesize and purify **12**. Yield 44%. ESI-MS calculated for $C_{18}H_{30}N_2O_2$ ([M + H]$^+$) 306.2307, found 306.2310.

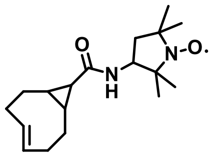

***sTCO -2,2,5,5-tetraethyl-1-pyrrolinyloxy free radical*** **(sTCO-tE5, 13):** Following the above procedure for making **11**, 3-aminomethyl-2,2,5,5-tetraethyl-1-pyrrolinyloxy free radical[1] was used to synthesize and purify **13**. Yield 42%. ESI-MS calculated for $C_{23}H_{38}N_2O_2$ ([M + H]$^+$) 374.2933, found 374.2910.

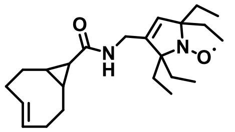

**Tetrazine amino acid (Tet4) rate constants measurement.** The solutions of tetrazine amino acids (0.2 mM) and sTCO-OH (1.0-6.0 mM) were made in PBS (137 mM NaCl, 2.7 mM KCl, 10 mM $Na_2HPO_4$, 1.8 mM $KH_2PO_4$, pH 7.4) with 2% methanol to ensure solubility of Tet ncAAs. The measured loss of tetrazine absorbance at 270 nm was used to determine the reaction rate of Tet ncAAs. Pseudo-first order conditions were employed with sTCO-OH in 5- to 60- fold excess of tetrazine amino acids at 25 $^oC$. All measurements were performed in triplicate and the resulted decay curves were fit to a single exponential equation. The mean values of pseudo first order rate constants ($k'$) were plotted against different concentration of sTCO-OH to obtain second order rate constants ($k_2$) from the slope of the plot (Figure S1).

**Selection of aminoacyl-tRNA synthetases for Tet4 incorporation**. To incorporate Tet4-Me and Tet4-Ph, a library of all amino acids at 5 sites (Asn311, Cys313, Val366, Trp382, and Gly 386) from *Methanosarcina barkeri* (Mb) system was chosen for its ability to incorporate large aromatic amino acids as described previously.[5, 6] The synthetase (RS) library was encoded on a kanamycin (Kn) resistant plasmid (pBK, 3000 bp) under control of the constitutive *Escherichia coli* GlnRS promoter and terminator (pBK-RS library). The pBK-RS library was mutated at the above-mentioned sites with NNK codons corresponding to all 20 natural amino acids where N is A, C, G, or T and K is G or T. The pBK-RS library, was transformed into DH10b cells containing a positive selection plasmid (pRep pylT) for "positive" selection steps and DH10b cells containing a negative selection plasmid (pYOBB2 pylT) for "negative" selection steps.

The positive selection plasmid, pRep pylT (10000 bp), encodes a mutant (*Mb*) pyrrolysyl-tRNA$_{CUA}$, an amber codon-disrupted chloramphenicol acetyltransferase, an amber codon-disrupted T7 RNA polymerase, a T7 promoter controlled GFP gene, and the tetracycline (Tcn) resistance marker. The negative selection plasmid, pYOBB2 pylT (7000 bp), encodes the mutant pyrrolysyl-tRNA$_{CUA}$, an amber codon-disrupted barnase under control of an arabinose promoter and rrnC terminator, and the chloramphenicol (Cm) resistance marker. pRep pylT electrocompetent cells and pYOBB2 pylT electrocompetent cells were made from DH10B cells carrying the respective plasmids and stored in 100 µL aliquots at −80 °C.

Following the general selection procedure, positive and negative selections were carried out on the pBK-library.[7] For positive selections, LB agar media plates containing 60 µg/mL chloramphenicol (Cm), 50 µg/mL kanamycin (Kn), 25 µg/mL tetracycline (Tcn) and 0.5 mM tetrazine amino acids were used. For negative selections, LB agar plates containing 25 µg/mL chloramphenicol (Cm), 50 µg/mL kanamycin (Kn) and 0.2% arabinose. The pBK-RS library was moved back and forth between positive and negative cell selections for two full rounds (positive 1, negative 1, positive 2, negative 2) The remaining pBK-RS library from the second negative selection transformed into DH10B cells containing the pALS to evaluate individual pBK-library members. The pALS plasmid contains the sfGFP reporter with a TAG codon at residue 150 as well as pyrrolysyl-tRNA$_{CUA}$. Individual colonies (96) were collected from the agar plate used to inoculate a 96-deep well plate containing 0.5 mL per well non-inducing media (NIM) containing 50 µg/mL Kn and 25 µg/mL Tcn. This block was cultured for 24 hrs. at 37 °C and 300 RPM. The non-inducing media saturated cells were used to inoculate into two 96-well plates containing 0.5 mL per well auto-inducing media (AIM) in presence and absence of Tet4 amino acids at 37 °C and 300 RPM. The cell culture and GFP expression was monitored by OD600 and fluorescence measurement at 24 hrs. and 48 hrs. This analysis led to four unique synthetases D4, E1, D10 and F2 which were able to selectively suppress the TAG interrupted GFP in presence of Tet4-Ph. None of the remaining pBK-library members were selective for encoding Tet4-Me.

**Efficiency and fidelity of selected synthetases.** The efficiency and fidelity of the selected synthetases were measured by expressing GFP150 in 50 mL AIM with and without 0.5 mM Tet4-Ph. DH10b cells containing the selected pBK-RSs and the pALS-GFP150TAG plasmids, were used to inoculate 5 mL of NIM containing kanamycin (50 µg/mL) and tetracycline (25 µg/mL). Cells were grown for 16 hours at 37 °C shaking at 250 rpm. The saturated NIM cultures (500 µL) were used to inoculate in 50 mL AIM containing kanamycin (50 µg/mL) and tetracycline (25 µg/mL) and measured fluorescence using a Turner Biosystems Picofluor fluorimeter diluting 100 µL cell culture in 1.9 mL water. Two synthetases D4 and E1 for Tet4-Ph were identified to have good efficiency and fidelity (Figure. S2A). The selected two synthetases are differ at two mutation sites C313A/G and W382I/V respectively (Figure. S2B).

**Permissivity screen for Tet4-Pyr with selected synthetases.** To assess if the top four selected RSs for Tet4-Ph could incorporate Tet4-Pyr, we evaluated expressions of GFP150TAG with Tet4-Pyr. Using the above efficiency and fidelity method, 5 mL AIM separate cultures were inoculated with cells containing the pBK-RSs/pALS-GFP150. Tet4-Pyr (0.5 mM) was added to cultures from a 50 mM DMF ncAA stock solution. Cultures were grown for 30-36 hours at 37 °C and 250 rpm. Fluorescence of the culture was measured by diluting 100 µL cell culture in 1.9 mL water. (Figure S3).

**Expression and purification of GFP150-TAG-Tet4.** Using expression conditions above for efficiency and fidelity measurements GFP150-Tet4 was expressed in 50 mL AIM and all cells were harvested after 36 hrs by centrifugation at 5000 rcf for 10 min. Media was removed, and cell pellets were stored at −80 °C. Cells were resuspended in wash buffer (NaCl 300 mM, $NaH_2PO_4$ 15.5 mM, $Na_2HPO_4$ 34.5 mM, imidazole 5 mM, pH 7.1). Cells were lysed using a Microfluidics M-110P microfluidizer (18,000 psi) and the lysate was collected in wash buffer. The lysate was clarified by centrifugation (21000 rcf, 30 mins.) and to the clarified supernatant TALON resin (300 µL bed volume) was added. Lysate was incubated with the resin for 1-2 hours gently rocking at 4° C. Resin and lysate were applied to a column and flow through was discarded. Resin was washed 5 times with 10 mL wash buffer. Protein was eluted with 250 µL elution buffer (NaCl 300 mM, $NaH_2PO_4$ 15.5 mM, $Na_2HPO_4$ 34.5 mM, imidazole 250 mM, pH 7.0). Protein concentration was determined by measuring absorbance at 280 nm. Protein purity was assessed using SDS-PAGE (Figure S4).

**Mobility Shift Assay.** The purified GFP-Wt, GFP150-Tet4-Ph/Pyr and MBP-Tet4-Ph variants were diluted to 50 µM in PBS. The protein was reacted with excess sTCO-PEG5k (500 µM) for 5-10 minutes in PBS at room temperature. Protein was denatured through the addition of Laemmli buffer and heated at 95 °C for 10 minutes. Samples were then analyzed using a 12% SDS-PAGE gel (Figure S4).

**Mass spectra of GFP-Tet4.** Purified GFP-TAG150-Tet4 was diluted to 50 µM and desalted using Zeba™ spin desalting column and analyzed using an FT LTQ mass spectrometer at the Oregon State University mass spectrometry facility. A 6545XT Q-TOF mass spectrometer interfaced with a 1290 Infinity II HPLC (Agilent Technologies, Santa Clara, CA) was used to verify reaction of purified GFP-TAG150-Tet4 with sTCO. Intact proteins were desalted and concentrated using a Poroshell 300SB-C18 column prior to electrospray. Spectra were deconvoluted using the Maximum Entropy deconvolution algorithm in BioConfirm. (Figure 1D, S5 and S20).

**Measuring reaction rates of GFP-Tet4-Ph/Pyr with sTCO-OH and sTCO-spin labels.** Fluorescence dequenching of pure GFP-Tet protein by reaction with sTCO-reagents was used to measure the rates of Tet reaction on proteins.[5] The fluorescence of GFP-Tet4-Ph and GFP-Tet4-Pyr in 3 mL of PBS (12 nmol) was measured (488 nm excitation, 509 nm emission, 5 points/second) for 50 seconds prior to the addition of 10 µL sTCO and sTCO-spin labels (0.03 -3 µmol sTCO reagent in PBS). Stock concentrations (10 - 900 µmol) of sTCO reagents were prepared in methanol. Reactions were monitored until the return of fluorescence stabilized. Curves were fitted to a single exponential equation using the curve-fitting program OriginPro 8.5 to determine kinetic constants. (Figure S6 and S7).

**Maltose binding protein (MBP)-Tet4-Ph expression and purification for spin labeling:**

**MBP cloning.** The wild type (Wt) and TAG variants (211, 278, 295, 322, 211-295 and 278-322) of the MBP genes with N-terminus His-tag were cloned in place of the GFP gene in the pALS plasmid. The MBP genes were amplified form the pETM-11 vectors[8] using the primers pALS-MBP-For and pALS-MBP-Rev and incorporated into the pALS plasmid at the cut sites XhoI and XbaI.

pALS-MBP-For (5'GTTTTTTGGGCTAACAGGAGGAATTAACATGAAACATCACCATCACCATCACCCCA-3')

pALS-MBP-Rev (5'GAGTTTTTGTTCGGGCNCAAGCTTCGCTCGAGTTATTATTTGGTGATGCGAGTCTGC-3').

***E. coli*** **MBP-Tet4-Ph expression and purification.** Single colonies of DH10B cells co-transformed with pBK-E1 and single/double TAG mutant pALS-MBP plasmids (pALS-MBP-Wt; pALS-MBP-211; pALS-MBP-278; pALS-MBP-295; pALS-MBP-322; pALS-MBP-211/295; pALS-MBP-278/322) were used to inoculate a 5 mL non-inducing culture containing 50 μg/mL Kn and 25 μg/mL Tet. The non-inducing culture was grown to saturation, shaking at 250 rpm and 37 °C. Autoinduction media (50 mL) with 50 μg/mL Kn and 25 μg/mL Tet, was inoculated with 0.7 mL of saturated non-inducing culture. When autoinducing cultures reached and OD of 0.5-0.7 0.3 mM Tet4-Ph was added. After 30 hours of shaking at 37 °C, cells were collected by centrifugation and stored at −80°C. The cell pellets were resuspended in TALON wash buffer (pH~7.4) with 5 mM imidazole and lysed using a microfluidizer (final volume 30 mL). Cell lysate was clarified by centrifugation and added to TALON cobalt resin (0.2 mL bed-volume) and incubated for 1 hr at 4 °C. Bound resin was washed with >50 mL volumes wash buffer. Protein was eluted by using 0.3 mL elution buffer containing 250 mM imidazole (pH ~ 7.1). Purified protein yields per liter of media were MBP-Wt (120 mg/L), MBP-211 (15 mg/L), MBP-295 (25 mg/L), MBP-278 (32 mg/L), MBP-322 (45 mg/L), MBP-211/295 (10 mg/L), MBP-278/322 (14 mg/L).

**In-vitro spin-labeling.** Purified single and double cysteine constructs of MBP were incubated at room temperature with 10 mM tris(2-carboxyethyl)phosphine for 10 minutes, concentrated with a 50-kDa molecular-weight cutoff (MWCO) spin concentrator (GE Healthcare) and purified by size exclusion chromatography (SEC) on a Superdex 200 10/300 GL column (GE Healthcare) equilibrated with EPR sample buffer (150 mM NaCl, 25 mM HEPES, pH 7.2). SEC fractions were immediately reacted with MTSL (Toronto Research Chemicals) at room temperature for ∼4 h, protected from light. Unreacted spin label was removed by desalting (PD-10) into EPR sample buffer and concentrated (50-kDa MWCO). Removal of residual free spin label and exchange into deuterated buffer (for DEER samples) was achieved by microdialysis (Pierce). Glycerol (CW samples) or d8-glycerol (DEER samples) was supplemented to a final concentration of 30% (vol/vol). For samples containing maltose, D-(+)-maltose was included at 1 – 5 mM, depending on the experiment. Samples for CW EPR were loaded into 1.0 mm outer diameter (OD), 0.7 mm inner diameter (ID) quartz capillaries (Sutter), sealed with wax, and maintained at room temperature until measurement. Samples for DEER spectroscopy were loaded into 1.5 mm OD/1.1 mm ID quartz tubes (Sutter) with flame-sealed bottoms and flash frozen in liquid nitrogen. Samples were stored at −80 °C until measurement.

For labeling of single or double GFP and MBP constructs containing Tet4-Ph, purified proteins were desalted into EPR sample buffer (PD-10) and reacted with 100-250 µM sTCO-tM6, sTCO-tM5, or sTCO-tE5 for 1 h at room temperature. Removal of free spin label and EPR sample preparation was performed in the same manner as described above for MTSL-labeled constructs.

**In-cell spin-labeling.** Free Tet4 ncAA was removed from HEK293T cells expressing Tet4-Ph constructs of GFP or MBP by exchanging the culture medium 3 times with fresh DMEM + 10% FBS over the course of 4 – 6 h with incubation at 37 °C, 5% $CO_2$. Following the final incubation, medium was removed and replaced with medium supplemented with 100 – 250 nM sTCO-tE5, depending on the experiment. Cells were incubated in labeling medium at room temperature for 10 – 15 minutes, after which the labeling medium was removed and the cells were washed once with PBS, detached from the surface with PBS + 5 mM EDTA and collected by centrifugation at 300 × g. Cell pellets were gently resuspended in a minimum volume of PBS. For CW EPR measurements, cells were loaded into 1.0 mm OD, 0.7 mm ID quartz capillaries, sealed with wax, briefly pelleted by centrifugation at 500 × g and immediately taken to the spectrometer for recording. For DEER experiments, resuspended cells were transferred to a 3 mm OD quartz EPR tube, pelleted by centrifugation at 300 × g and frozen in liquid $N_2$. DEER samples were stored at −80 °C until measurement.

**EPR spectroscopy.** Continuous-wave EPR spectra were recorded at room temperature on a Bruker EMX spectrometer operating at X-band frequency (~9.8 GHz) equipped with a Bruker ER 4123D dielectric resonator. Spectra were recorded with 100 kHz field modulation with a sweep rate of 1.8 G/s and a modulation amplitude of 2 G. CW EPR spectra were background subtracted and baseline corrected in LabVIEW™. Spin concentrations were calculated by double integration of the field-modulated spectrum and comparison to a standard curve of TEMPO free radical (Sigma). Labeling efficiencies were calculated as the spin concentration obtained by double integration divided by the total protein concentration estimated from optical absorbance at 280 nm using an extinction coefficient for 6xHis-MBP of 67,280 $M^{-1}\,cm^{-1}$. Reported labeling efficiencies are the average efficiencies between apo and maltose-bound samples for each construct.

Pulsed EPR experiments were performed at Q-band frequency (~34 GHz) using a Bruker EleXsys E580 spectrometer with an overcoupled Bruker EN 5107D2 (*in vitro* samples) or ER 5106-QT2 (in-cell samples) resonator. Pulses were generated with a Bruker SpinJet AWG and amplified with a 300 W TWT amplifier (Applied Systems Engineering). DEER experiments were performed at 45–50 K using a variable-temperature cryogen-free system (Bruker/ColdEdge). The deadtime-free, four-pulse DEER sequence $[(\pi/2)_{probe} — \tau_1 — (\pi)_{probe} — \tau_1 + t — (\pi)_{pump} — \tau_2 - t — (\pi)_{probe} — \tau_2 — (echo)]$ was employed with a 260 ns or 400 ns $\tau_1$ delay and $\tau_2$ delays ranging from 2–6 μs depending on the sample. For samples in deuterated buffer, $\tau_1$ delays were incremented by 16 ns over 8 steps to suppress deuterium ESEEM contributions to the DEER trace. The pump pulse was implemented as a 150 ns sech/tanh pulse with a frequency bandwidth of 80 MHz and a truncation parameter (β) of 10. The magnetic field was adjusted such that the pump pulse was centered near the maximum of the nitroxide field-swept spectrum. Probe pulses were 60 ns ($\pi/2$ and $\pi$) Gaussian-shaped pulses applied at a frequency 80 MHz below that of the pump pulse center frequency. All pulses were compensated for resonator bandwidth using the *pulse* function of EasySpin-v6.0-dev.[9] Raw time-domain DEER traces were background-corrected and distance distributions were calculated by Tikhonov regularization in LongDistances (by Christian Altenbach, available at www.biochemistry.ucla.edu/Faculty/Hubbell). Error bands in the distance distribution were calculated by stochastic addition of random noise to the DEER signal and variation of the background parameters followed by recalculation of the distance distribution using the error analysis feature of LongDistances with default values. Error bands are plotted as the mean ±1 standard deviation from 100 independent calculations with parameters varied as described above.

**In-silico spin label modeling.** Spin-labeled side chain ensembles of Tet4-Ph-tE5 were modeled onto MBP and GFP structures using chiLife, an open-source Python package for site-directed spin label modeling (https://github.com/StollLab/chiLife).[10] Tet4-Ph-tE5 ncAA with N-terminal acetylation and C-terminal amidation was constructed as the 1,4-dihydropyridazine product with *cis* fusion of the cyclopropane ring and the eight-membered ring set to the 'half-chair' conformation.[4] A rotamer library consisting of 601 low-energy structures was generated using the GFN-FF force field in CREST.[12] Rotamer ensembles and simulated DEER distance distributions for each site-pair were generated in chiLife using off-rotamer sampling of mobile dihedral angles using the default values.

### Eukaryotic expression materials and methods

**Toxicity Screen of Tetrazines**.[5] HEK293T cells were plated in a 96-well plate at approximately 10% confluency. Cells were incubated for 48 h with Tet4 ncAA or 1% DMSO, and the cell viability was measured using CellTiter Glo assay kit (Promega) according to the manufacturer's instruction. Briefly, 25 μl of CellTiter Glo reagent was added to each well and incubated for 10 min at RT. The signal was measured for 1 sec using TR717 microplate luminometer (Berthold, Germany) and WinGlow software version 1.25 (Berthold Technologies). The data was normalized to vehicle control and fit to a curve using non-linear regression method with GraphPad Prism 5. n = 3 ± 95% CI.

**Transfection of HEK293T cells and imaging method.** HEK293T cells were plated in a 24-well plate at the density of 40% confluency. Cells were transfected the next day using jetPRIME reagent according to the manufacturer's protocol. Briefly, 67 ng of pAcBac1-NES-D4-RS or pAcBac1-NES-E1-RS and 555 ng of pAcBac1-GFP-150TAG plasmid DNA were diluted with 50 uL of jetPRIME buffer. To the diluted DNA, 1.2 uL of jetPRIME reagent was added and vortexed. The complex was incubated for 10 min at room temperature. It was gently added to the cells. Tet4 ncAA or 0.1% DMSO were added to the cells immediately. Cells were incubated 18 h ~ 48 h before analysis. GFP expression was verified by fluorescence microscopy on EVOS FL imaging system.

**Flow Cytometry Assessment**. Following published protocol Jang et al.[5]

**In-gel Fluorescence Analysis of *in vivo* Protein Labeling.** Following published protocol Jang et al.[5]

**Live-cell spin-labeling – fluorescence microscopy.** HEK293T/17 cells expressing GFP150-Tet4-Ph were plated on circular (25 mm diameter) poly-(D)-lysine-coated microscope glass coverslips and incubated in DMEM + 10% FBS until the desired cell density was reached. Coverslips were then encased within a homebuilt perfusion device and bathed in DMEM + 10% FBS. Live cells were imaged using a Nikon Eclipse TE2000-E microscope with a 10× water immersion objective. GFP was excited using epifluorescence with a Lambda SC SmartShutter controller (Sutter Instruments) with 470/40 nm excitation and 515/30 nm emission filters. GFP fluorescence was recorded with 10 ms exposures using an Evolve 512 EMCCD camera (Photometrics) and MetaMorph software (Molecular Devices). Cells were manually perfused with DMEM + 10% FBS medium containing 1 μM sTCO-tE5 spin label, and cells were imaged every 500 ms for a total of 240 images. Mean cell fluorescence for each timepoint was calculated by first defining a region of interest (ROI) for each cell based on the final image in the time series using the particle analysis tool in ImageJ. Particle ROIs were then applied across all images in the time series. Particle intensities for each ROI were background subtracted and the mean particle intensity was calculated for each image. Mean fluorescence values were then normalized by the mean fluorescence value from the first image, recorded prior to application of spin label.

### Gene and Protein Sequence:

**GFP-wt (Protein):**
MVSKGEELFTGVVPILVELDGDVNGHKFSVRGEGEGDATNGKLTLKFICTTGKLPVPWPTLVTTLTYGVQC
FSRYPDHMKRHDFFKSAMPEGYVQERTISFKDDGTYKTRAEVKFEGDTLVNRIELKGIDFKEDGNILGHKL
EYNFNSHNVYITADKQKNGIKANFKIRHNVEDGSVQLADHYQQNTPIGDGPVLLPDNHYLSTQSVLSKDPN
EKRDHMVLLEFVTAAGITHGMDELYKGSHHHHHH

**GFP-wt (DNA):**
ATGGTTAGCAAAGGTGAAGAACTGTTTACCGGCGTTGTGCCGATTCTGGTGGAACTGGATGGTGATGT
GAATGGCCATAAATTTAGCGTTCGTGGCGAAGGCGAAGGTGATGCGACCAACGGTAAACTGACCCTG
AAATTTATTTGCACCACCGGTAAACTGCCGGTTCCGTGGCCGACCCTGGTGACCACCCTGACCTATGGC
GTTCAGTGCTTTAGCCGCTATCCGGATCATATGAAACGCCATGATTTCTTTAAAAGCGCGATGCCGGA
AGGCTATGTGCAGGAACGTACCATTAGCTTCAAAGATGATGGCACCTATAAAACCCGTGCGGAAGTTA
AATTTGAAGGCGATACCCTGGTGAACCGCATTGAACTGAAAGGTATTGATTTTAAAGAAGATGGCAAC
ATTCTGGGTCATAAACTGGAATATAATTTCAACAGCCATAATGTGTATATTACCGCCGATAAACAGAA
AAATGGCATCAAAGCGAACTTTAAAATCCGTCACAACGTGGAAGATGGTAGCGTGCAGCTGGCGGAT

CATTATCAGCAGAATACCCCGATTGGTGATGGCCCGGTGCTGCTGCCGGATAATCATTATCTGAGCAC
CCAGAGCGTTCTGAGCAAAGATCCGAATGAAAAACGTGATCATATGGTGCTGCTGGAATTTGTTACCG
CCGCGGGCATTACCCACGGTATGGATGAACTGTATAAAGGCAGCCACCATCATCATCACCATTAA

**E1-PylRS (Protein):**
MDKKPLDVLISATGLWMSRTGTLHKIKHHEVSRSKIYIEMACGDHLVVNNSRSCRTARAFRHHKYRKTCK
RCRVSDEDINNFLTRSTESKNSVKVRVVSAPKVKKAMPKSVSRAPKPLENSVSAKASTNTSRSVPSPAKSTP
NSSVPASAPAPSLTRSQLDRVEALLSPEDKISLNMAKPFRELEPELVTRRKNDFQRLYTNDREDYLGKLERD
ITKFFVDRGFLEIKSPILIPAEYVERMGINNDTELSKQIFRVDKNLCLRPMLAPTLYNYLRKLDRILPGPIKVF
EVGPCYRKESDGKEHLEEFTMVSFGQMGSGCTRENLEALIKEFLDYLEIDFEIVGDSCMVYGDTLDIMHGD
LELSSAHVGPVSLDREWGIDKPVIGAGFGLERLLKVMHGFKNIKRASRSESYYNGISTNL

**E1-PylRS (DNA):**
ATGGATAAAAAACCGCTGGATGTGCTGATTAGCGCGACCGGCCTGTGGATGAGCCGTACCGGCACCCT
GCATAAAATCAAACATCATGAAGTGAGCCGCAGCAAAATCTATATTGAAATGGCGTGCGGCGATCATC
TGGTGGTGAACAACAGCCGTAGCTGCCGTACCGCGCGTGCGTTTCGTCATCATAAATACCGCAAAACC
TGCAAACGTTGCCGTGTGAGCGATGAAGATATCAACAACTTTCTGACCCGTAGCACCGAAAGCAAAAA
CAGCGTGAAAGTGCGTGTGGTGAGCGCGCCGAAAGTGAAAAAAGCGATGCCGAAAAGCGTGAGCCGT
GCGCCGAAACCGCTGGAAAATAGCGTGAGCGCGAAAGCGAGCACCAACACCAGCCGTAGCGTTCCGA
GCCCGGCGAAAAGCACCCCGAACAGCAGCGTTCCGGCGTCTGCGCCGGCACCGAGCCTGACCCGCAG
CCAGCTGGATCGTGTGGAAGCGCTGCTGTCTCCGGAAGATAAAATTAGCCTGAACATGGCGAAACCGT
TTCGTGAACTGGAACCGGAACTGGTGACCCGTCGTAAAAACGATTTCAGCGCCTGTATACCAACGAT
CGTGAAGATTATCTGGGCAAACTGGAACGTGATATCACCAAATTTTTTGTGGATCGCGGCTTTCTGGA
AATTAAAAGCCCGATTCTGATTCCGGCGGAATATGTGGAACGTATGGGCATTAACAACGACACCGAAC
TGAGCAAACAAATTTTCCGCGTGGATAAAAACCTGTGCCTGCGTCCGATGCTGGCCCCGACCCTGTAT
AACTATCTGCGTAAACTGGATCGTATTCTGCCGGGTCCGATCAAAGTTTTTGAAGTGGGCCCGTGCTAT
CGCAAAGAAAGCGATGGCAAAGAACACCTGGAAGAATTCACCATGGTTTCGTTTGGTCAAATGGGCA
GCGGCTGCACCCGTGAAAACCTGGAAGCGCTGATCAAAGAATTCCTGGATTATCTGGAAATCGACTTC
GAAATTGTGGGCGATAGCTGCATGGTGTATGGCGATACCCTGGATATTATGCATGGCGATCTGGAACT
GAGCAGCGCGCATGTGGGTCCGGTTAGCCTGGATCGTGAATGGGGCATTGATAAACCGGTGATTGGCG
CGGGTTTTGGCCTGGAACGTCTGCTGAAAGTGATGCATGGCTTCAAAAACATTAAACGTGCGAGCCGT
AGCGAAAGCTACTATAACGGCATTAGCACGAACCTGTAA

**D4-PylRS (Protein):**
MDKKPLDVLISATGLWMSRTGTLHKIKHHEVSRSKIYIEMACGDHLVVNNSRSCRTARAFRHHKYRKTCK
RCRVSDEDINNFLTRSTESKNSVKVRVVSAPKVKKAMPKSVSRAPKPLENSVSAKASTNTSRSVPSPAKSTP
NSSVPASAPAPSLTRSQLDRVEALLSPEDKISLNMAKPFRELEPELVTRRKNDFQRLYTNDREDYLGKLERD
ITKFFVDRGFLEIKSPILIPAEYVERMGINNDTELSKQIFRVDKNLCLRPMLAPTLYNYLRKLDRILPGPIKIFE
VGPCYRKESDGKEHLEEFTMVSFAQMGSGCTRENLEALIKEFLDYLEIDFEIVGDSCMVYGDTLDIMHGDL
ELSSAHVGPVSLDREWGIDKPIIGAGFGLERLLKVMHGFKNIKRASRSESYYNGISTNL

**D4-PylRS (DNA):**
ATGGATAAAAAACCGCTGGATGTGCTGATTAGCGCGACCGGCCTGTGGATGAGCCGTACCGGCACCCT
GCATAAAATCAAACATCATGAAGTGAGCCGCAGCAAAATCTATATTGAAATGGCGTGCGGCGATCATC
TGGTGGTGAACAACAGCCGTAGCTGCCGTACCGCGCGTGCGTTTCGTCATCATAAATACCGCAAAACC
TGCAAACGTTGCCGTGTGAGCGATGAAGATATCAACAACTTTCTGACCCGTAGCACCGAAAGCAAAAA
CAGCGTGAAAGTGCGTGTGGTGAGCGCGCCGAAAGTGAAAAAAGCGATGCCGAAAAGCGTGAGCCGT
GCGCCGAAACCGCTGGAAAATAGCGTGAGCGCGAAAGCGAGCACCAACACCAGCCGTAGCGTTCCGA
GCCCGGCGAAAAGCACCCCGAACAGCAGCGTTCCGGCGTCTGCGCCGGCACCGAGCCTGACCCGCAG
CCAGCTGGATCGTGTGGAAGCGCTGCTGTCTCCGGAAGATAAAATTAGCCTGAACATGGCGAAACCGT

TTCGTGAACTGGAACCGGAACTGGTGACCCGTCGTAAAAACGATTTTCAGCGCCTGTATACCAACGAT
CGTGAAGATTATCTGGGCAAACTGGAACGTGATATCACCAAATTTTTTGTGGATCGCGGCTTTCTGGA
AATTAAAAGCCCGATTCTGATTCCGGCGGAATATGTGGAACGTATGGGCATTAACAACGACACCGAAC
TGAGCAAACAAATTTTCCGCGTGGATAAAAACCTGTGCCTGCGTCCGATGCTGGCCCCGACCCTGTAT
AACTATCTGCGTAAACTGGATCGTATTCTGCCGGGTCCGATCAAAATTTTTGAAGTGGGCCCGTGCTAT
CGCAAAGAAAGCGATGGCAAAGAACACCTGGAAGAATTCACCATGGTTTCGTTTGCTCAAATGGGCA
GCGGCTGCACCCGTGAAAACCTGGAAGCGCTGATCAAAGAATTCCTGGATTATCTGGAAATCGACTTC
GAAATTGTGGGCGATAGCTGCATGGTGTATGGCGATACCCTGGATATTATGCATGGCGATCTGGAACT
GAGCAGCGCGCATGTGGGTCCGGTTAGCCTGGATCGTGAATGGGGCATTGATAAACCGATTATTGGCG
CGGGTTTTGGCCTGGAACGTCTGCTGAAAGTGATGCATGGCTTCAAAAACATTAAACGTGCGAGCCGT
AGCGAAAGCTACTATAACGGCATTAGCACGAACCTGTAA

**MBP-wt (Protein):**
MKHHHHHHPMSDYDIPTTENLYFQGAMAKTEEGKLVIWINGDKGYNGLAEVGKKFEKDTGIKVTVEHPD
KLEEKFPQVAATGDGPDIIFWAHDRFGGYAQSGLLAEITPDKAFQDKLYPFTWDAVRYNGKLIAYPIAVEA
LSLIYNKDLLPNPPKTWEEIPALDKELKAKGKSALMFNLQEPYFTWPLIAADGGYAFKYENGKYDIKDVGV
DNAGAKAGLTFLVDLIKNKHMNADTDYSIAEAAFNKGETAMTINGPWAWSNIDTSKVNYGVTVLPTFKG
QPSKPFVGVLSAGINAASPNKELAKEFLENYLLTDEGLEAVNKDKPLGAVALKSYEEELAKDPRIAATMEN
AQKGEIMPNIPQMSAFWYAVRTAVINAASGRQTVDEALKDAQTRITK

**MBP-wt (DNA):**
ATGAAACATCACCATCACCATCACCCCATGAGCGATTACGACATCCCCACTACTGAGAATCTTTATTTT
CAGGGCGCCATGGCGAAAACTGAAGAAGGTAAACTGGTAATCTGGATTAACGGCGATAAAGGCTATA
ACGGTCTCGCTGAAGTCGGTAAGAAATTCGAGAAAGATACAGGAATTAAAGTCACCGTTGAGCATCC
GGATAAACTGGAAGAGAAATTCCCACAGGTTGCGGCAACTGGCGATGGCCCTGACATTATCTTCTGGG
CACACGACCGCTTTGGTGGCTACGCTCAATCTGGCCTGTTGGCTGAAATCACCCCGGACAAAGCGTTC
CAGGACAAGCTGTATCCGTTCACCTGGGATGCCGTACGTTACAACGGCAAGCTGATTGCTTACCCGAT
CGCTGTTGAAGCGTTATCGCTGATTTATAACAAAGATCTGCTGCCGAACCCGCCAAAAACCTGGGAAG
AGATCCCGGCGCTGGATAAAGAACTGAAAGCGAAAGGTAAGAGCGCGCTGATGTTCAACCTGCAAGA
ACCGTACTTCACCTGGCCGCTGATTGCTGCTGACGGGGGTTATGCGTTCAAGTATGAAAACGGCAAGT
ACGACATTAAAGACGTGGGCGTGGATAACGCTGGCGCGAAAGCGGGTCTGACCTTCCTGGTTGACCTG
ATTAAAAACAAACACATGAATGCAGACACCGATTACTCCATCGCAGAAGCTGCCTTTAATAAAGGCGA
AACAGCGATGACCATCAACGGCCCGTGGGCATGGTCCAACATCGACACCAGCAAAGTGAATTATGGT
GTAACGGTACTGCCGACCTTCAAGGGTCAACCATCCAAACCGTTCGTTGGCGTGCTGAGCGCAGGTAT
TAACGCCGCCAGTCCGAACAAAGAGCTGGCAAAAGAGTTCCTCGAAAACTATCTGCTGACTGATGAA
GGTCTGGAAGCGGTTAATAAAGACAAACCGCTGGGTGCCGTAGCGCTGAAGTCTTACGAGGAAGAGT
TGGCGAAAGATCCACGTATTGCCGCCACTATGGAAAACGCCCAGAAAGGTGAAATCATGCCGAACAT
CCCGCAGATGTCCGCTTTCTGGTATGCCGTGCGTACTGCGGTGATCAACGCCGCCAGCGGTCGTCAGA
CTGTCGATGAAGCCCTGAAAGACGCGCAGACTCGCATCACCAAATAA

### Scheme 1-3:

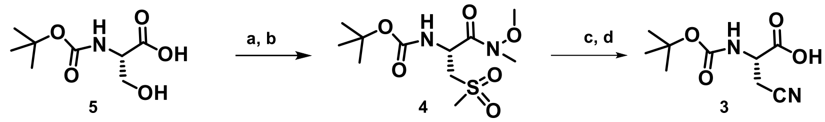

(a) EDC.HCl, N-Methylmorpholine, N,O dimethyl hydroxylamine hydrochloride, dry DCM, -5 - 0 °C,
(b) Methanesulfonyl chloride, triethyl amine, dry DCM, 0 °C (c) NaCN, dry DMSO, 55 °C (d) LiOH.H20, THF-H2O (5:1)

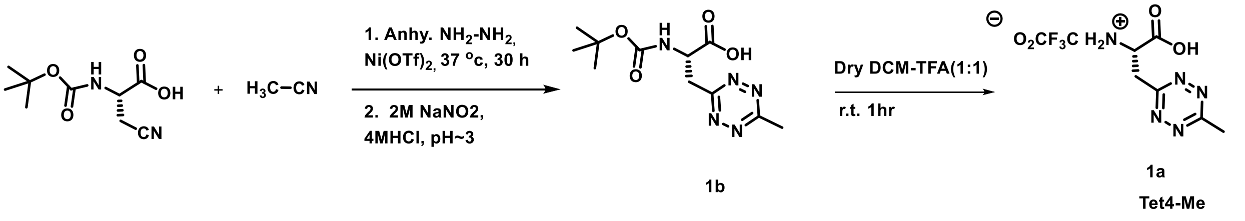

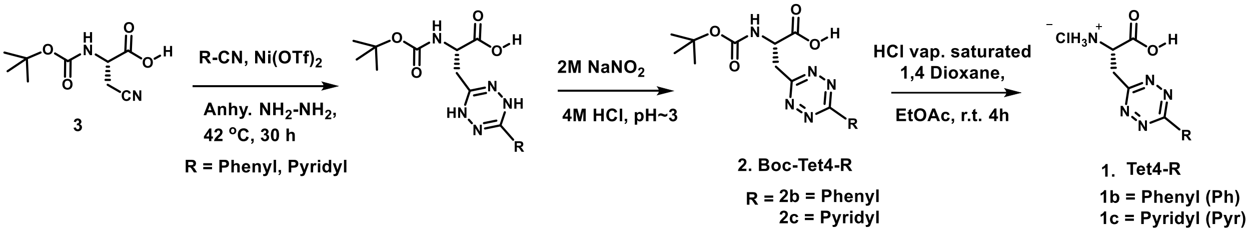

**Scheme S1.** Synthesis of s-tetrazine derivatives of alanine (Tet4-ncAAs).

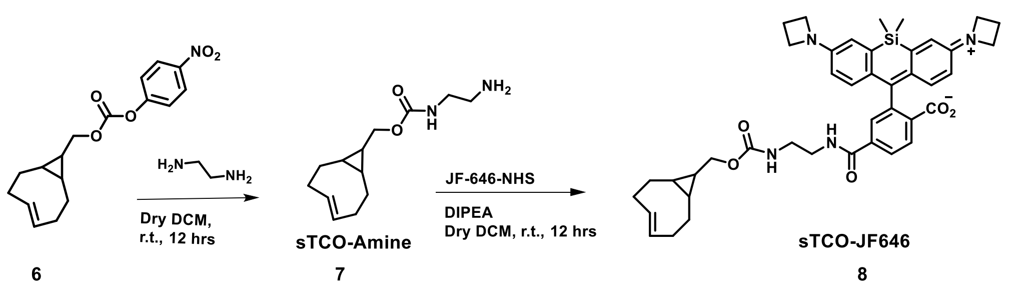

**Scheme S2. Synthesis of sTCO-JF646.**

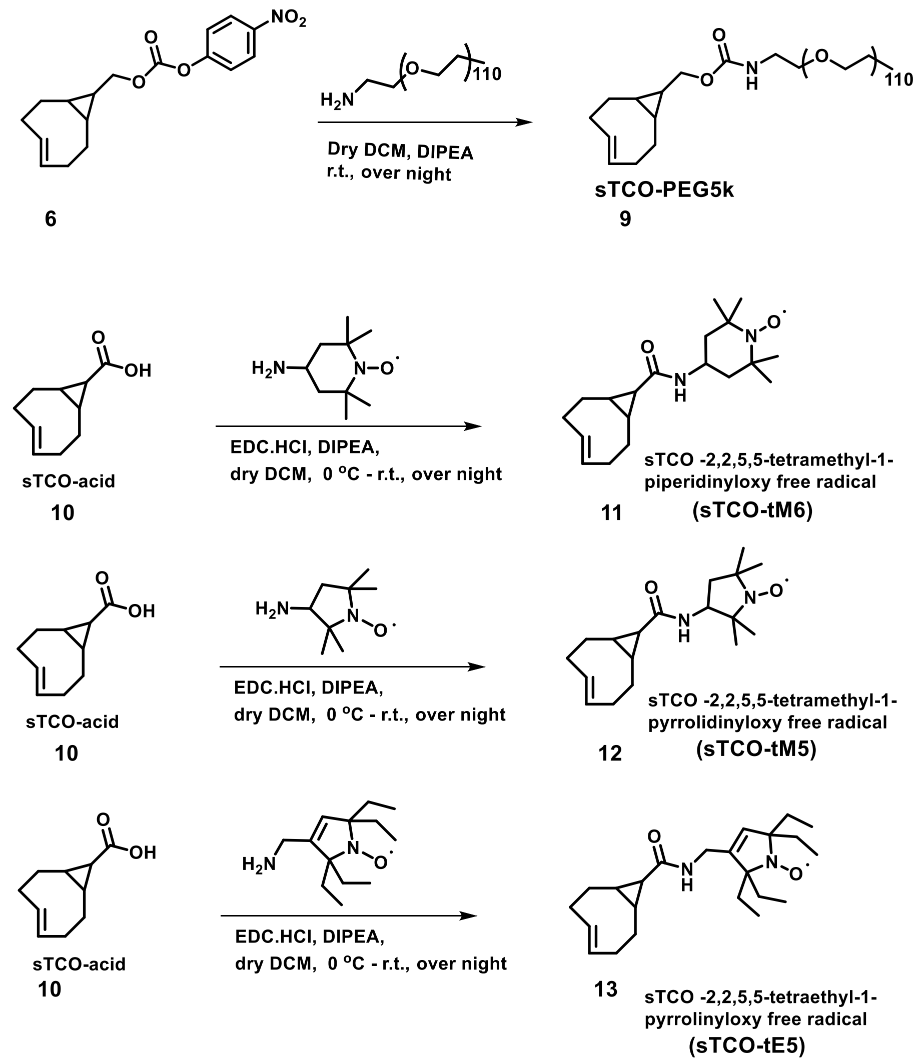

**Scheme S3.** Synthesis of sTCO-PEG5k and sTCO-tM6, sTCO-tM5, and sTCO-tE5 spin labels.

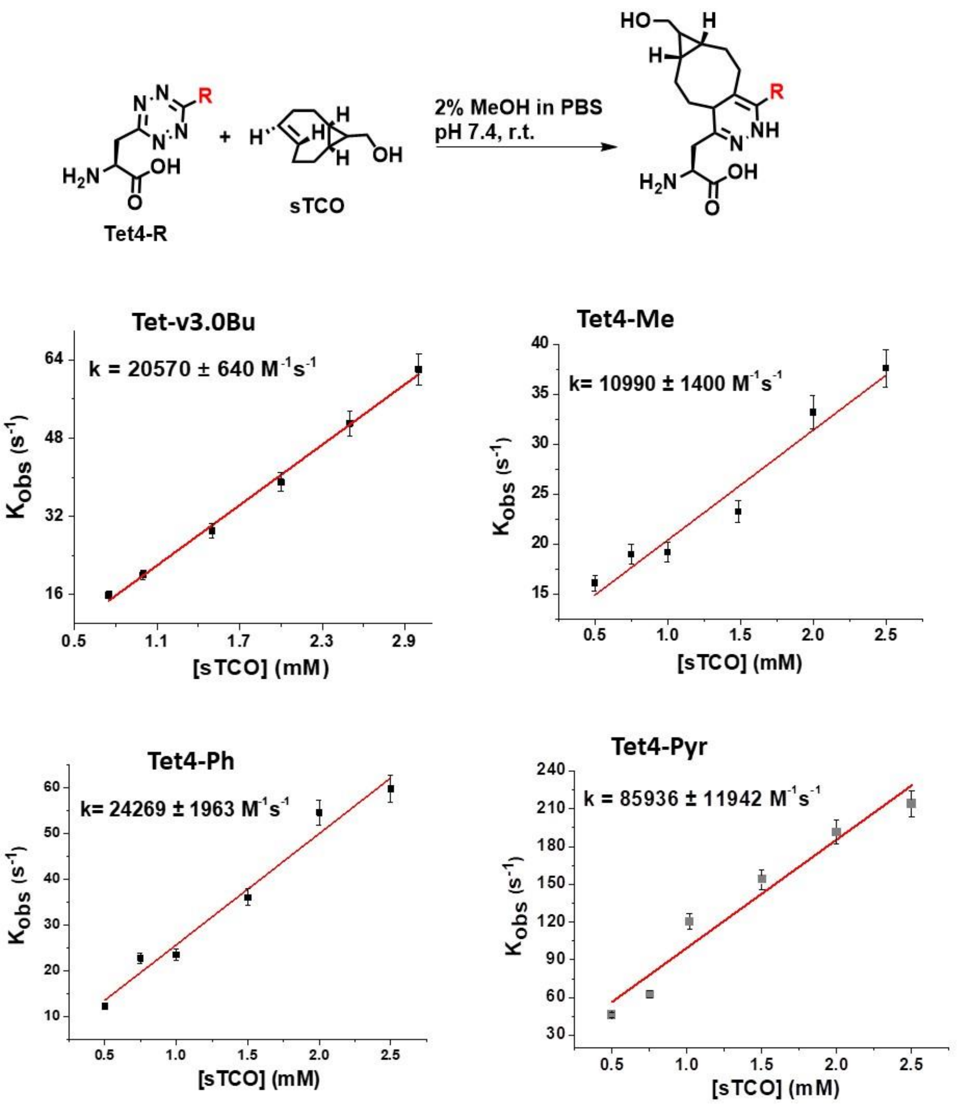

**Figure S1**. **Measurement of reaction rates for Tet4 amino acids with sTCO-OH.** The loss of tetrazine signal at 270 nm was measured over time upon the addition of sTCO-OH at different concentrations. Plot of pseudo first order rate constants ($k'$) against concentration of sTCO-OH to determine the second order rate constants ($k_2$) of cycloaddition reaction of Tet4 derivatives with cyclopropyl fused *trans*-cyclooctene alcohol (sTCO-OH). All measurements were performed in triplicate at 25 ºC in PBS.

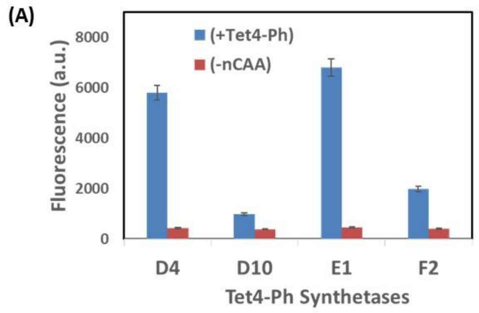

(B)

| Mutation sites | Asn (N) 202 | Ile (I) 287 | Glu (Q) 302 | Asn (N) 311 | Cys (C) 313 | Val (V) 366 | Trp (W) 382 |
|---|---|---|---|---|---|---|---|
| **D4** | --- | --- | --- | **Ser (S)** | **Ala (A)** | **His (H)** | **Ile (I)** |
| D10 | --- | --- | --- | Ser (S) | Gly (G) | Arg (R) | Leu (L) |
| **E1** | --- | **Val (V)** | --- | **Ser (S)** | **Gly (G)** | **His (H)** | **Val (V)** |
| F2 | --- | --- | --- | Ser (S) | Gly (G) | His (H) | Ile (I) |

**Figure S2:** (A) Expression efficiency and fidelity of the selected synthetase were measured in triplicate by quantifying the fluorescence of TAG suppressed GFP in the presence and absence of Tet4-Ph (0.5 mM). The RS-D4 and E1 resulted in expressed GFP yields of 55 - 60 mg/L culture. (B) Sequence differences at mutation sites of the selected top synthetase for Tet4-Ph incorporation.

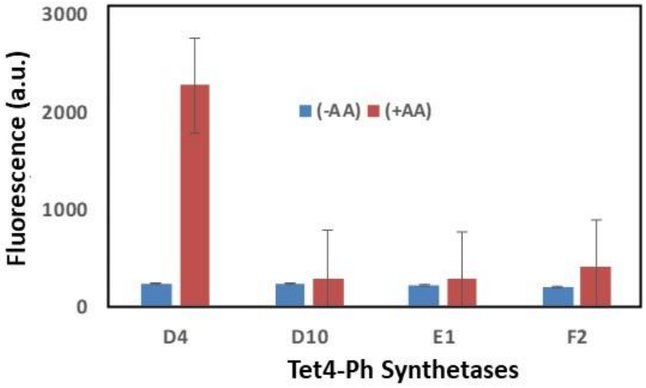

**Figure S3: Permissibility test for Tet4-Pyr.** Expression efficiency and fidelity of the selected synthetase were measured in triplicate by quantifying the fluorescence of TAG suppressed GFP in the presence and absence of Tet4-Pyr (0.5 mM). Only the synthetase D4 has incorporated the Tet4-Pyr. Yield ~ 40 mg/L.

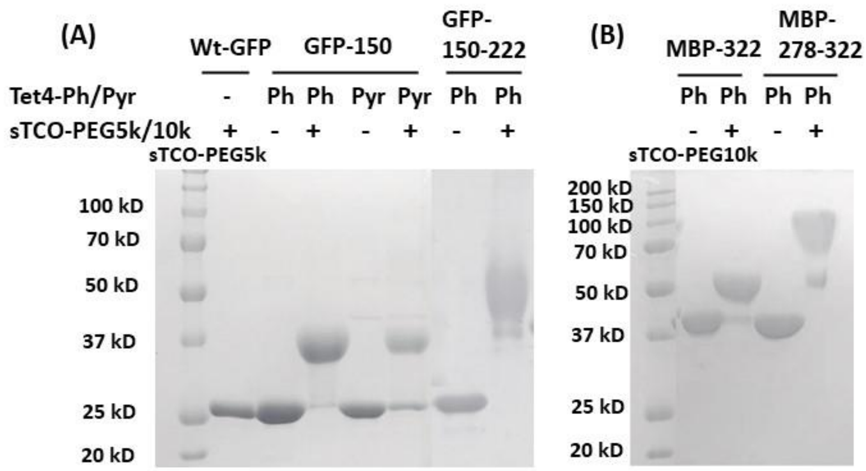

**Figure S4: SDS-PAGE mobility shift assay**. Labeling efficiency of (A) GFP-Tet4-Ph/Pyr and (B) MBP-Tet4-Ph verified by SDS-PAGE mobility shift upon reaction with sTCO-PEG5k/10k.

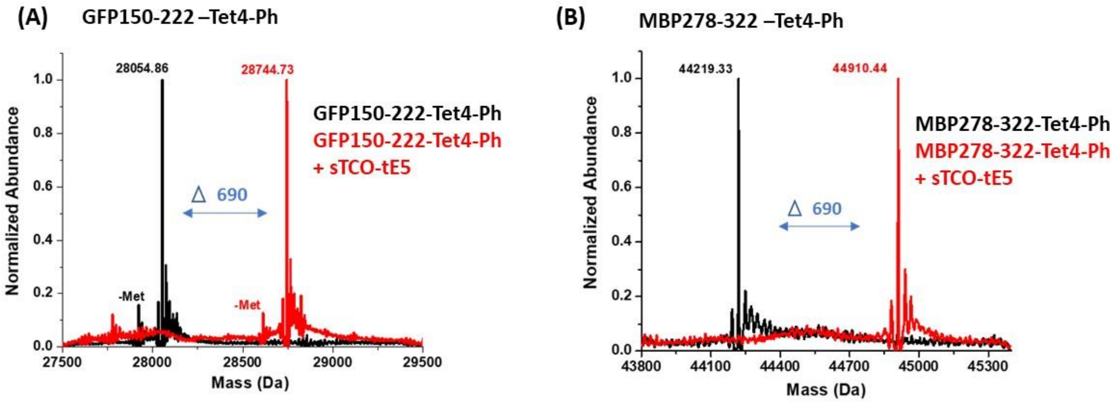

**Figure S5. ESI-Q-TOF mass spectrometry confirmed dual incorporation of Tet4-Ph into GFP and MBP and quantitative labeling with sTCO-spin label**. ESI mass spectroscopy analysis of purified GFP-150/222-Tet4-Ph and MBP-278/322-Tet4-Ph confirms high fidelity encoding of dual Tet-4.0Ph and quantitative double labeling with sTCO-tE5. Purified GFP150/222-Tet4-Ph and MBP-278/322-Tet4-Ph (black) and reacted with 5-fold molar excess of sTCO-tE5 for 10 minutes (red). (A) Cal. Mass of GFP-wt: 27827.02 Da avg; GFP-150/222-Tet4-Ph observed: 28054.9 Da avg, (expected: 28054.1 Da avg); GFP-150/222-Tet4-Ph + sTCO-tE5 observed: 28744.7 Da avg, (expected: 28744.5 Da avg). (B) Cal. Mass of MBP-wt: 44023.9 Da avg.; MBP-278/322-Tet4-Ph observed 44219.3 Da avg (expected: 44219.9 Da avg.); MBP-278/322-Tet4-Ph + sTCO-tE5 observed: 44910.4 Da avg. (expected: 44910.6 Da avg.). The lower mass peak labeled with '-Met' is a loss of n-terminal methionine and upper mass peaks are salt sodium and potassium adducts.

**Table S1**: Mass spectra analysis of *E. coli* and *HEK293T* cell expressed GFP-Tet4-Ph/Pyr and MBP-Tet4-Ph variants in presence and absence of sTCO reagents.

| ***E. coli* expressed proteins** | | | | | |
|---|---|---|---|---|---|
| Proteins | Residues at TAG sites | Theoretical Mass (Da avg.) | Observed Mass (Da avg.) | Difference (Da) | Modifications |
| Wt-GFP | Asn150 | 27827.3 | 27826.8 | -0.5 | - |
| GFP150-Tet4-Ph | Asn150/ Tet4-Ph | 27940.1 | 27941.5 | +1.4 | Tet4-Ph incorporation |
| | | | 27810.1 | -130 | -Met |
| GFP150-Tet4-Ph + sTCO-OH | Asn150/ Tet4-Ph | 28064.1 | 28065.5 | +1.4 | sTCO addition and $N_2$ loss |
| | | | 27934.1 | -130 | -Met |
| GFP150-Tet4-Pyr | Asn150/ Tet4-Pyr | 27941.1 | 27941.3 | +0.2 | Tet4-Pyr incorporation |
| | | | 27810.6 | -130.5 | -Met |
| GFP150-Tet4-Pyr + sTCO-OH | Asn150/ Tet4-Pyr | 28065.11 | 28066.2 | +1.1 | sTCO addition and $N_2$ loss |
| | | | 27933.8 | -131.3 | -Met |
| GFP150/222-Tet4-Ph | Asn150/ Tet4-Ph and Leu222/ Tet4-Ph | 28054.1 | 28054.9 | +0.8 | Dual Tet4-Ph incorporation |
| | | | 27922.5 | -132.4 | -Met |
| GFP150/222-Tet4-Ph + sTCO-tE5 | Asn150/ Tet4-Ph and Leu222/ Tet4-Ph | 28744.5 | 28744.7 | +0.2 | Double sTCO-tE5 addition and $N_2$ loss |
| | | | 28612.9 | -131.8 | -Met |
| Wt-MBP | Glu278-Glu322 | 44023.9 | 44022.7 | -1.2 | - |
| MBP278/222-Tet4-Ph | Glu278/ Tet4-Ph and Glu322/ Tet4-Ph | 44219.9 | 44219.3 | -0.6 | Dual Tet4-Ph incorporation |
| MBP278/222-Tet4-Ph+ sTCO-tE5 | Glu278/ Tet4-Ph and Glu322/ Tet4-Ph | 44910.6 | 44910.4 | -0.2 | Double sTCO-tE5 addition and $N_2$ loss |
| ***HEK293T* expressed proteins** | | | | | |
| wt-GFP | Asn150 | 29141.5 | 29141.8 | +0.3 | - |
| GFP150-Tet4-Ph | Asn150/ Tet4-Ph | 29254.4 | 29255.7 | +1.3 | Tet4-Ph incorporation |
| GFP150-Tet4-Ph + sTCO-OH | Asn150/ Tet4-Ph | 29378.3 | 29379.8 | +1.5 | sTCO addition and $N_2$ loss |
| GFP150-Tet4-Pyr | Asn150/ Tet4-Pyr | 29255.4 | 29255. 5 | +0.1 | Tet4-Pyr incorporation |
| | | | 29243.2 | -12.2 | Tetrazine degradation and oxadiazole formation |
| GFP150-Tet4-Pyr + sTCO-OH | Asn150/ Tet4-Pyr | 29379.3 | 29380.3 | +1.0 | sTCO addition and $N_2$ loss |
| | | | 29243.2 (Unreactive) | -136.1 | Tetrazine degradation (Fig. S21). |

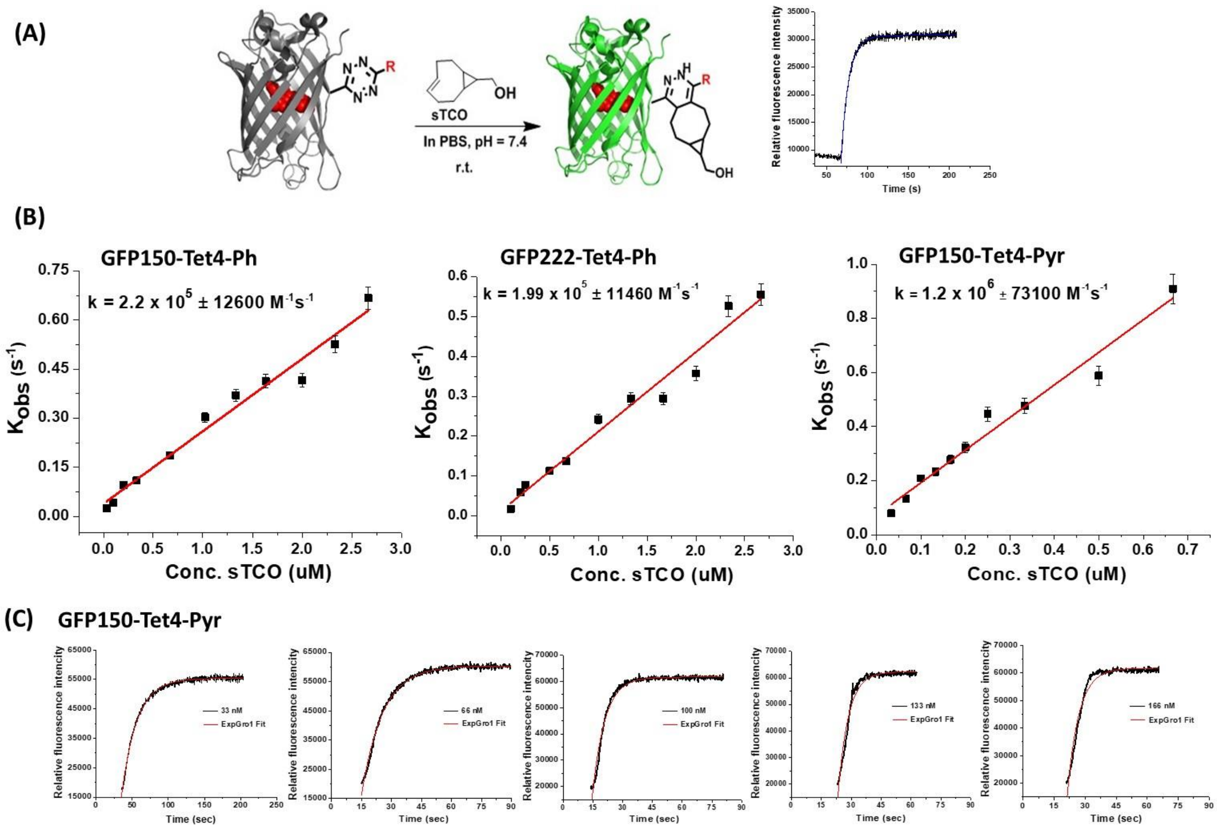

**Figure S6. Measurement of reaction rates for GFP-Tet4-Ph and GFP-Tet4-Pyr with sTCO-OH.** (A) Schematic representation of GFP fluorescence quenched by the tetrazine amino acids at site150 and 222 and the GFP fluorescence returned upon reaction with sTCO-OH. (B) Plots of pseudo first order rate constants ($k'$) against concentration of sTCO-OH to determine the second order rate constants ($k_2$) for reaction of GFP-Tet4 with sTCO-OH. (C) Representative measured fluorescence enhancement for reaction of GFP150-Tet4-Pyr (conc. 12 nM) with low sTCO concentrations (33 nM – 166 nM) with single exponential fits use in pseudo-first order reaction kinetics model.

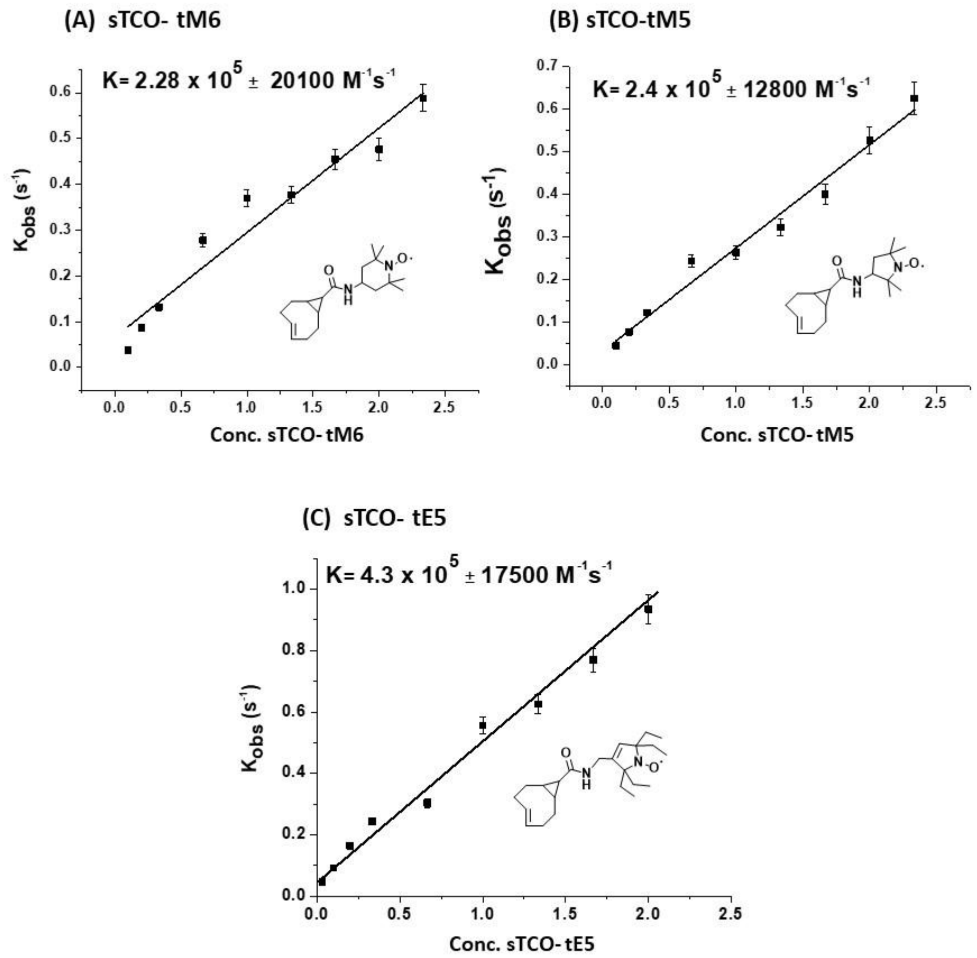

**Figure S7.** Plots of pseudo first-order rate constants (k′) against concentration of sTCO spin-labels to determine the second order rate constants ($k_2$) for the reaction of GFP150-Tet4-Ph with sTCO spin labels (A) sTCO-tM6 (B) sTCO-tM5 (C) sTCO-tE5.

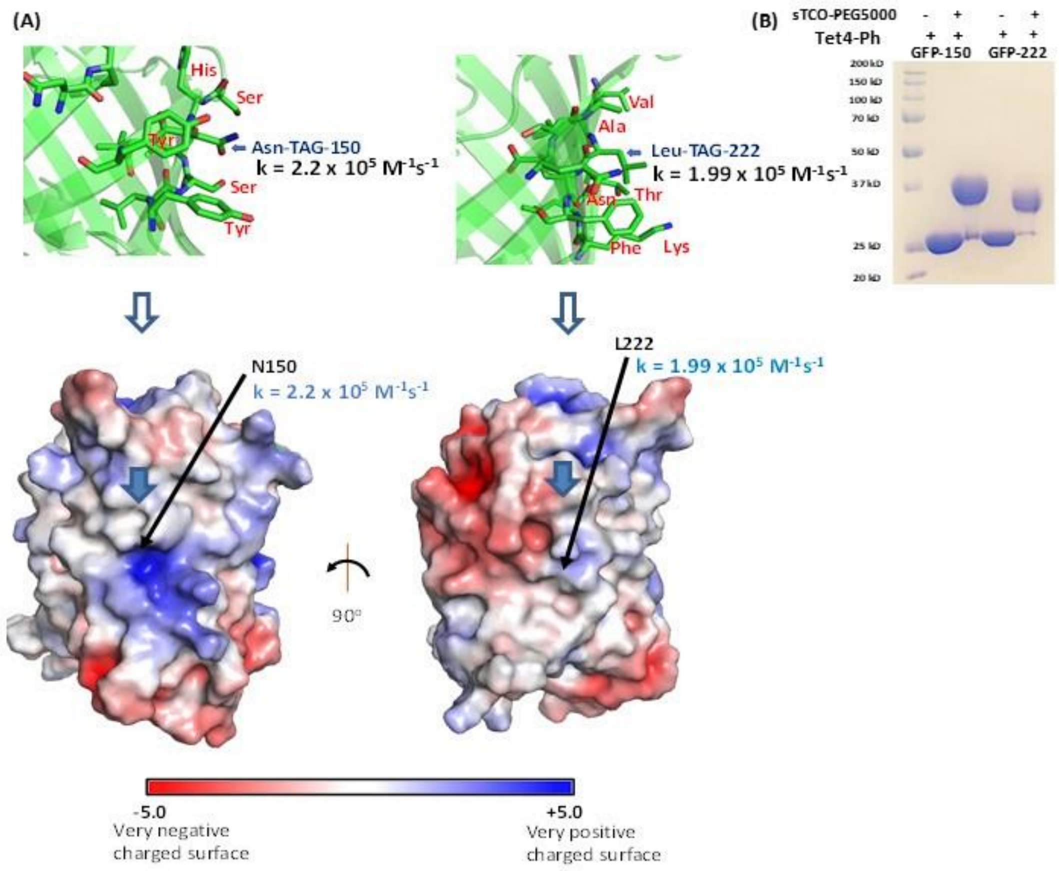

**Figure S8.** (A) Schematic presentation of electrostatic environments of GFP-TAG sites at 150 and 222. The GFP-Tet4-Ph shows very similar reactivity with sTCO-OH at both TAG sites 150 and 222 which are in different electrostatic environments. (B) The SDS-PAGE mobility shift in the presence and absence of sTCO-PEG5k verified reactivity of GFP-Tet4-Ph at both TAG sites150 and 222.

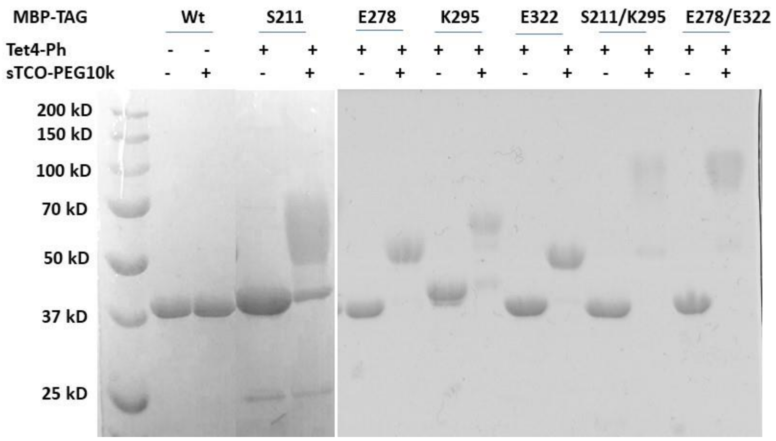

**Figure S9. Purity and reactivity estimates of MBP-Tet4-Ph constructs**. SDS-PAGE gel (Coomassie stain) shows mobility shift of MBP constructs in the presence and absence of sTCO-PEG10k (2 µg MBP-Tet4-Ph, 10 eqv. sTCO-PEG10k, in 25 µL PBS (pH~7.1), 10 minutes at r.t.).

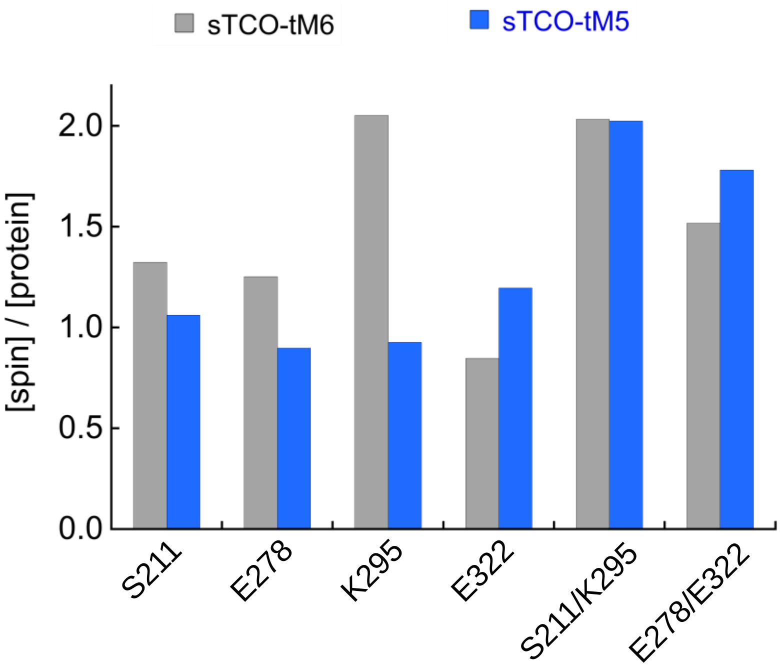

**Figure S10.** Spin-labeling efficiencies of sTCO-tM6 (gray) and sTCO-tM5 (blue) for single and double Tet4-Ph MBP constructs purified from *E. coli*. Spin concentrations were estimated by double integration of the CW EPR spectra (see methods) and total protein concentration was estimated from absorbance at 280 nm ($\varepsilon$ = 67,280 $M^{-1}cm^{-1}$).

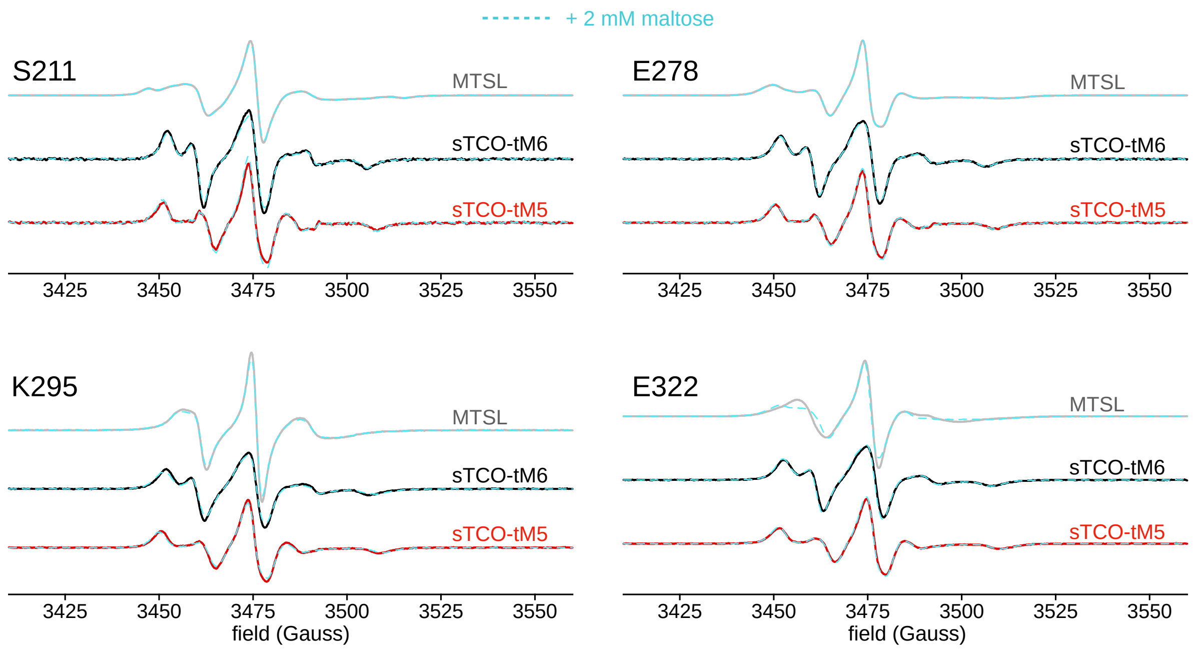

**Figure S11.** Room temperature X-band CW-EPR spectra of singly spin-labeled MBP constructs. *Gray* traces are spectra from cysteine mutants spin-labeled with MTSL whereas *black* and *red* spectra are from Tet4-Ph mutants spin-labeled with sTCO-tM6 and sTCO-tM5, respectively. Spectra for each construct were also recorded in the presence of 2 mM maltose (*dashed cyan*) and are shown overlayed on the apo spectra. All spectra are normalized by total spin concentration as determined by double integration of the baseline-corrected spectra.

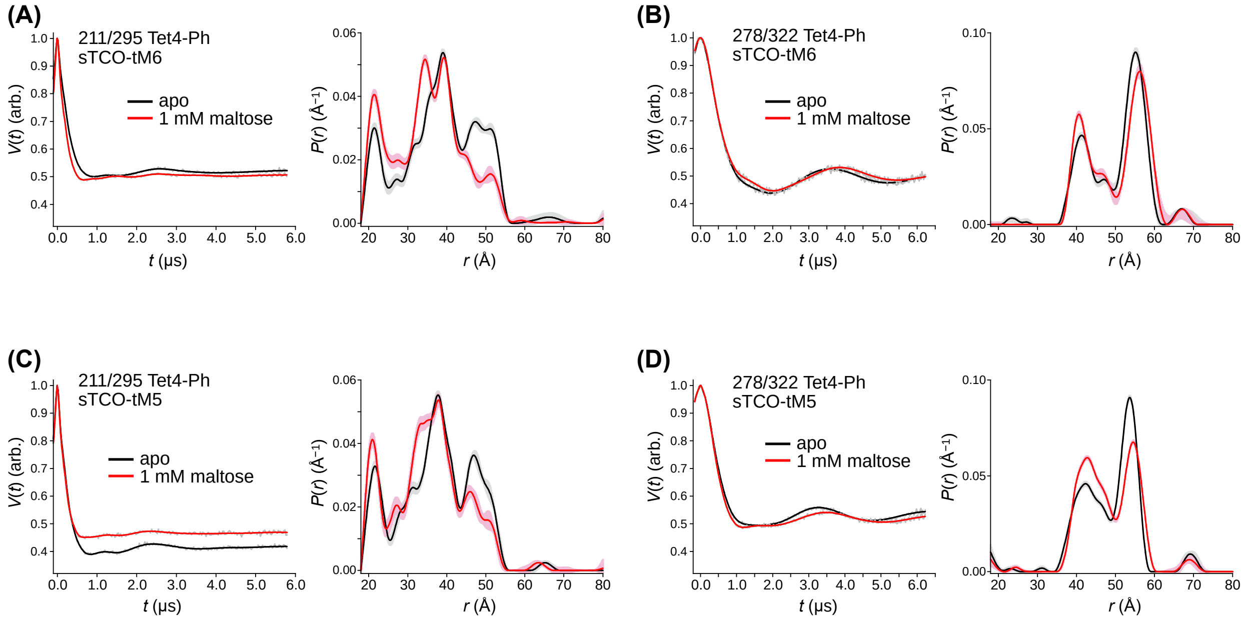

**Figure S12.** Background-corrected DEER time-traces and distance distributions for *in vitro*-purified and spin-labeled double Tet4-Ph MBP constructs in the absence of maltose (black) and in the presence of 1 mM maltose (red). Calculated distance distributions were obtained by non-parametric fitting of the time-domain signal with Tikhonov regularization using the LongDistances software package, written by Christian Altenbach and freely available from https://sites.google.com/site/altenbach/labview-programs/epr-programs/long-distances. Error bands on the distributions were generated by the Error function of LongDistances using default values.

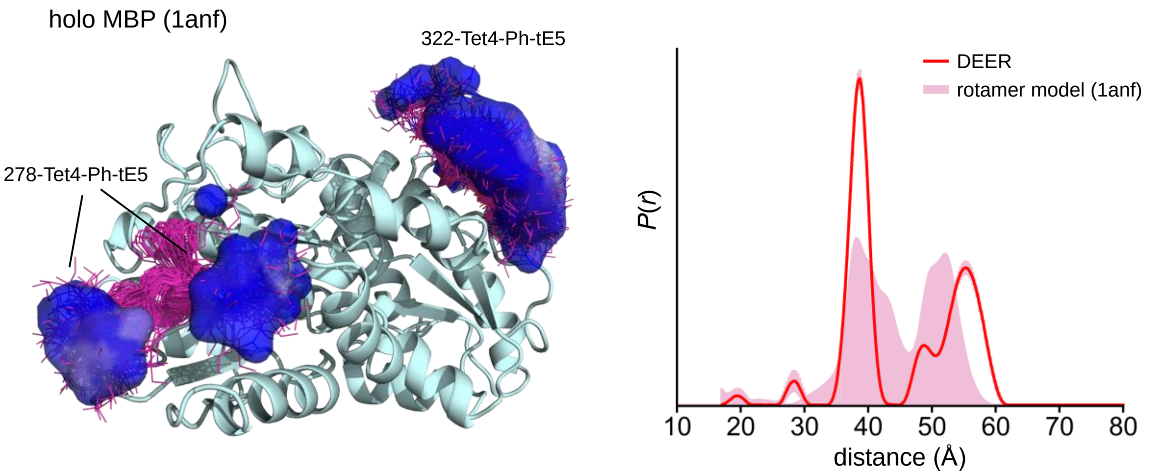

**Figure S13.** In silico modeling of sTCO-tE5 conjugated to Tet4-Ph at residues 278 and 322 of maltose-bound MBP (pdb accession 1anf) predicts that spin labels at 278 will predominantly reside in two separate rotameric clusters. The resulting simulated inter-spin distance distribution calculated from this rotameric model (pink filled) is distinctly bimodal and agrees well with the experimental distance distribution obtained by DEER (red trace). Spin label modeling was performed with the open source Python module chiLife,[10] available at https://github.com/StollLab/chiLife.

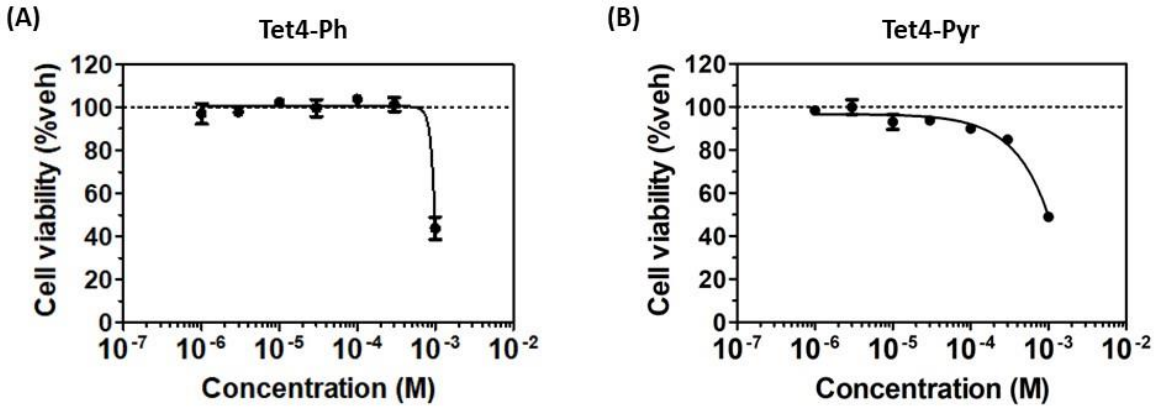

**Figure S14**. **The effect of Tet4 ncAA concentration on mammalian cell viability**. *HEK293T* cells were incubated for 48 h with (A) Tet4-Ph, (B) Tet4-Pyr, or 1% DMSO as indicated. Cell viability was compromised above 0.3 mM tetrazine amino acid. The cell viability was examined by CellTiter Glo a  ssay, and the data was normalized using a vehicle control.

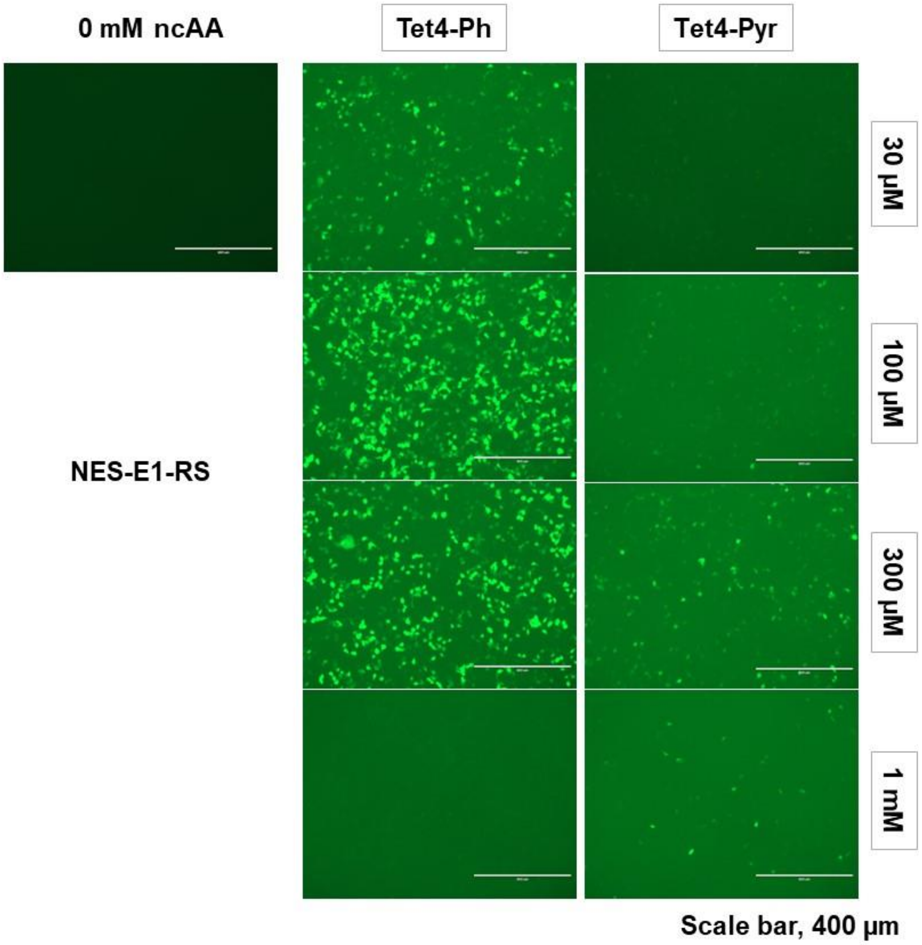

**Figure S15. The effect of Tet4 amino acids on GFP-TAG suppression in *HEK293T* cells.** The plasmid pAcBac1-E1-RS in presence of nuclear export sequences (NES) was used to transfect *HEK293T* cells along with pAcBac1-GFP-150TAG in the presence of 30 µM – 1 mM Tet4-Ph/Pyr or 0.1% DMSO. After 18 h, GFP fluorescence images were captured using EVOS FL imaging system.

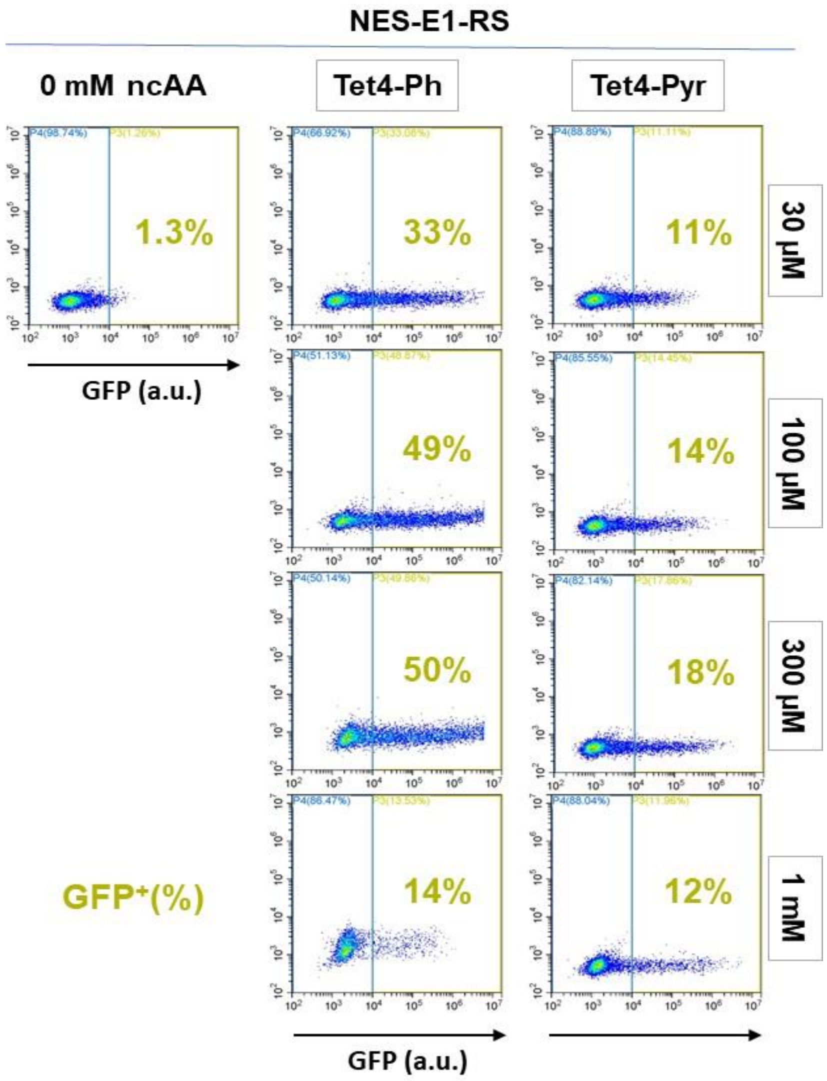

**Figure S16.** Flow cytometry analysis of the figure S15, GFP expression efficiency in presence of Tet4 in *HEK293T* cells with pAcBac1-NES-E1-RS at 30 µM – 1 mM concentrations.

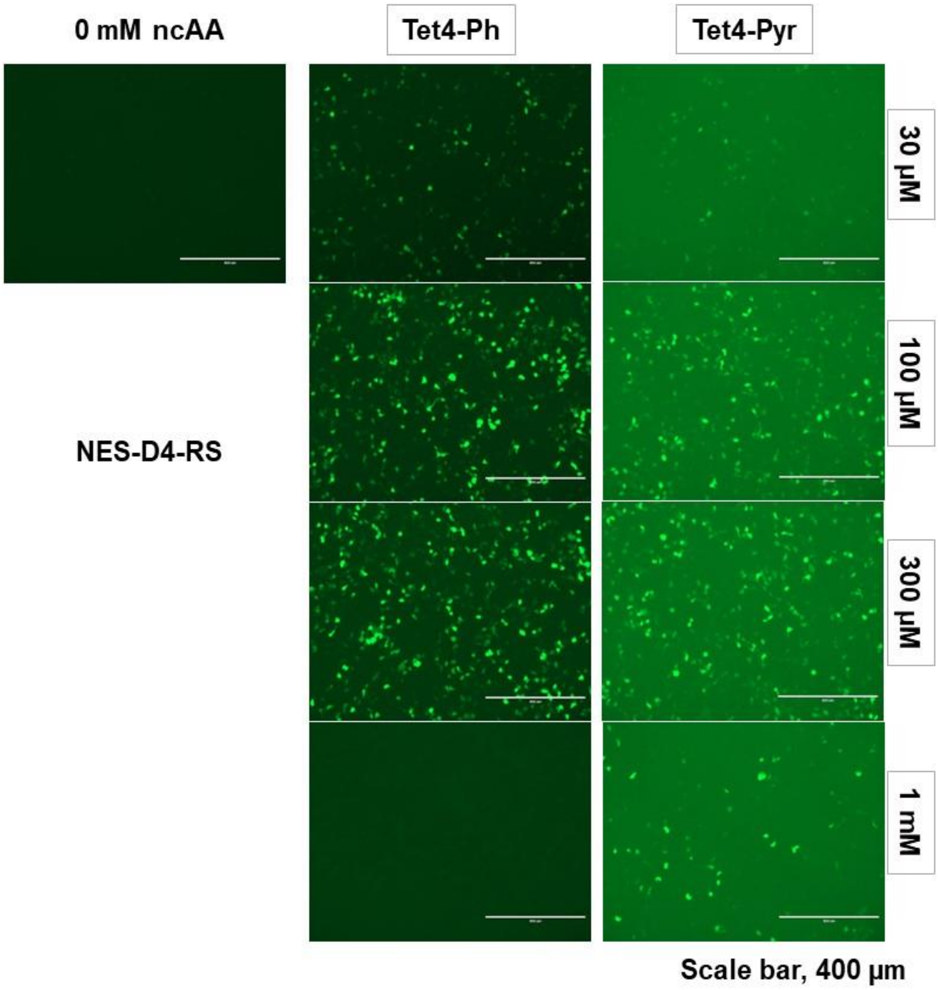

**Figure S17. The effect of Tet4 amino acids on GFP-TAG suppression in *HEK293T* cells.** The plasmid pAcBac1-D4-RS in presence of nuclear export sequences (NES) was used to transfect *HEK293T* cells were transfected with pAcBac1-D4-RS along with pAcBac1-GFP-150TAG in the presence of 30µM – 1mM Tet4-Ph/Pyr or 0.1% DMSO. After 18 h, GFP fluorescence images were captured using EVOS FL imaging system.

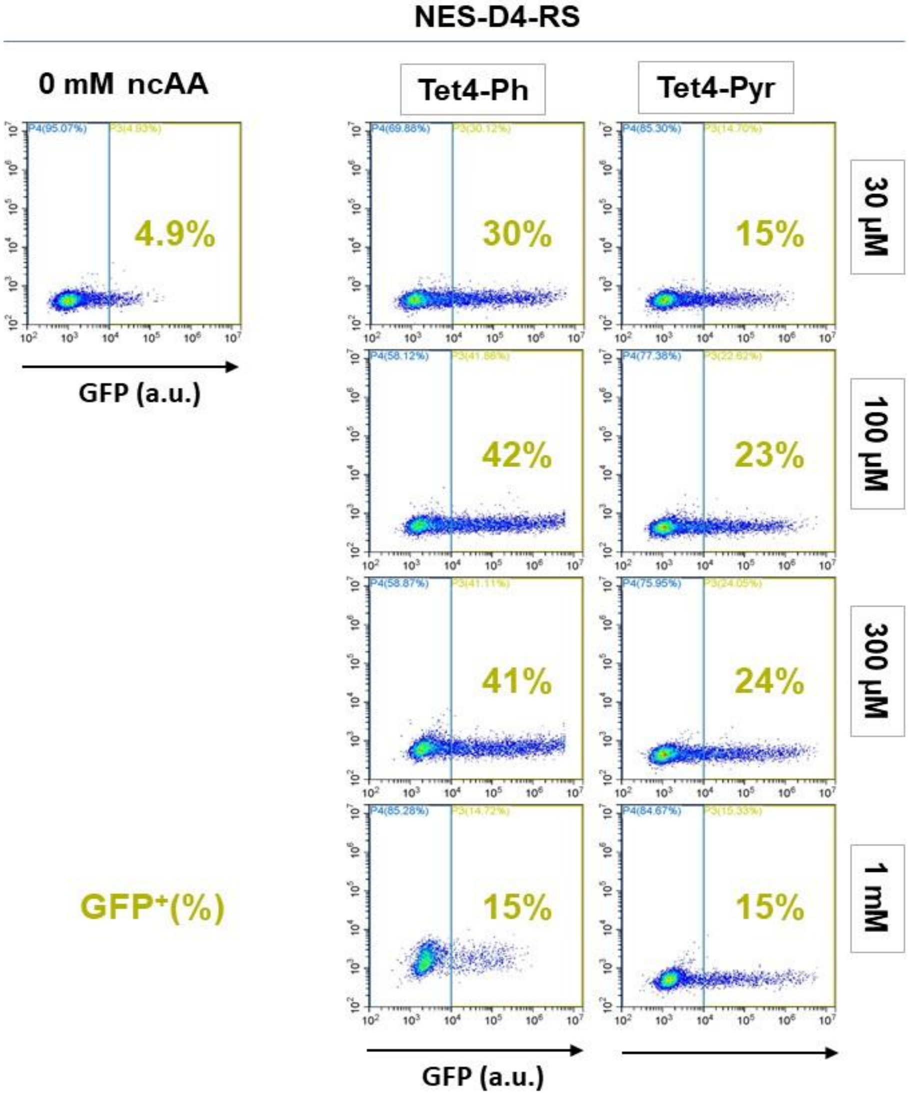

**Figure S18.** Flow cytometry analysis of the figure S17, GFP expression efficiency in presence of Tet4 in *HEK293T* cells with pAcBac1-NES-D4-RS at 30 μM – 1 mM concentration.

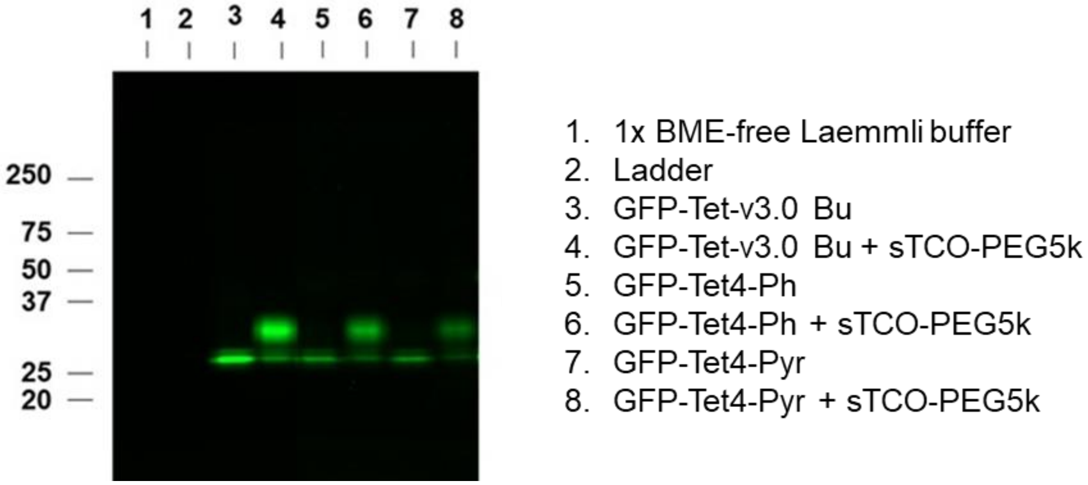

**Figure S19. *In vitro* mobility shift analysis of Tet4 protein in mammalian cell lysates**. A gel mobility shift assay was performed to verify the reactivity of Tet4 on protein expressed in HEK293T cells. HEK293T cells were transfected with pAcBac1-R2-84-RS and pAcBac1-GFP-150TAG in the presence of 30 µM Tet-v3.0Bu (lanes 3, 4), pAcBac1-NES-E1-RS and pAcBac1-GFP-150TAG in the presence of 30 µM Tet4-Ph (lanes 5, 6), or pAcBac1-NES-D4-RS and pAcBac1-GFP-150TAG in the presence of 30 µM Tet4-Pyr (lanes 7, 8). Cells were washed three times with culture media and once with PBS. Crude cell lysates were prepared in non-reducing Laemmli buffer and incubated for 30 min with 91 µM sTCO-PEG5k (lanes 4, 6, 8) or with vehicle (lanes 3, 5, 7). The lysates were separated by SDS-PAGE and fluorescent gel images were captured using Bio-Rad ChemiDoc system.

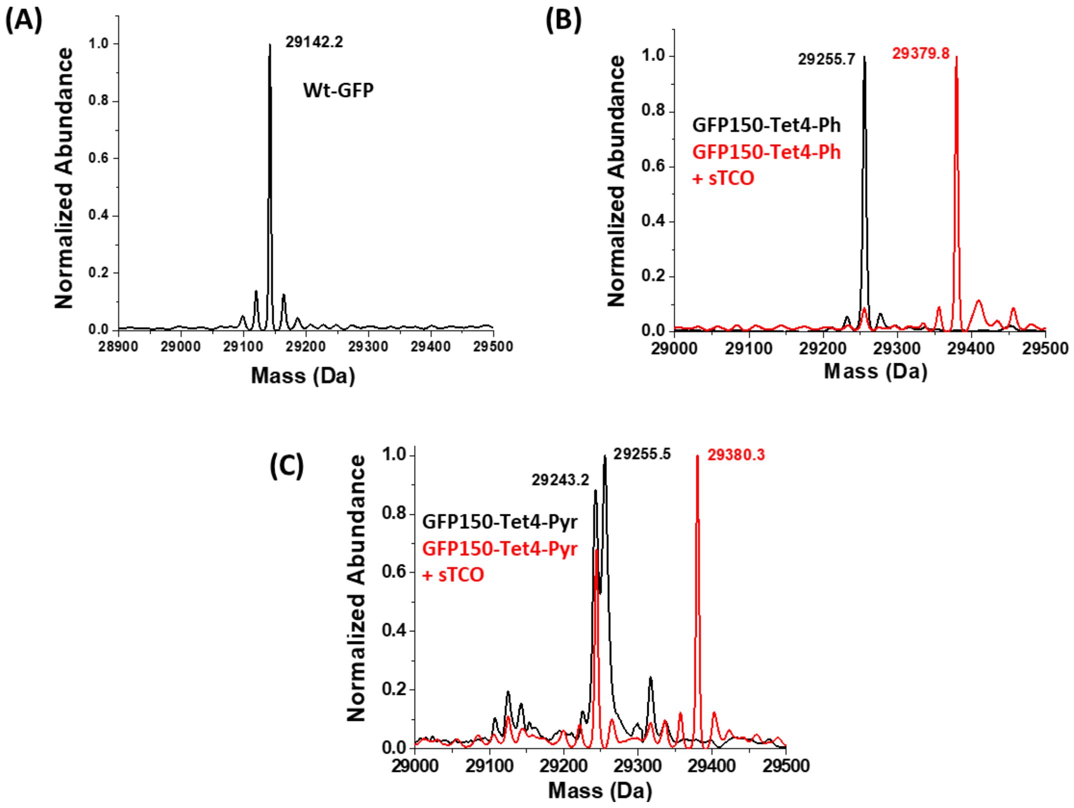

**Figure S20.** ESI-Q-TOF mass spectrometry of purified GFP-Tet4 from HEK293T cells confirms high fidelity encoding of single Tet-4.0 amino acid into GFP and labeling with sTCO. (A) Wt-GFP shows average mass 29142.2 Da (predicted 29141.49 Da). (B) GFP-Tet4-Ph observed: 29255.7 Da avg, (expected: 29254.4 Da avg); GFP-Tet4-Ph + sTCO observed: 29379.8 Da avg, (expected: 29378.3 Da avg); (C) GFP-Tet4-Pyr observed: 29243.2 and 29255.5 Da avg, (expected: 29255.4 Da avg); GFP-Tet4-Pyr + sTCO observed 29243.2 and 29380.3 Da avg, (expected: 29379.2 Da avg). (see above Table S1 for analysis).

**Tet4-Pyr degradation in *HEK293* cells expressed GFP150-Tet4-Pyr.**

The MS analysis of *HEK293T* cells expressed GFP150-Tet4-Pyr (Table S1) shows two peaks: one at 29255.5 Da avg. for single site Tet4-Pyr incorporation and a second at 29243.2 Da avg. which is 12 Da avg. unit less than expected mass. When the sample was exposed to sTCO the GFP150-Tet4-Pyr, major peak 29255.5 Da avg. completely shifted to 29380.3 Da Avg. due to the addition of sTCO mass 124.2 Da avg. and loss of $N_2$ but the protein at 29243.2 Da remains unreactive.

While electron withdrawing groups on 1,2,4,5-tetrazines increases their reactivity with strained alkenes but it also makes them more electrophilic and susceptible to nucleophilic attack. Hunter et al. have reported that under basic conditions the tetrazines are converted to a stable and unreactive oxadiazol derivatives which molecular mass 12 unit less than the parent molecule.[11] The MS data for Tet4-Pyr suggested that the peak at 29243.2 Da avg. corresponds to a portion of GFP150-Tet4-Pyr that gets converted to its oxadiazole derivative and is therefore not reactive to the sTCO reagent (Figure S21).

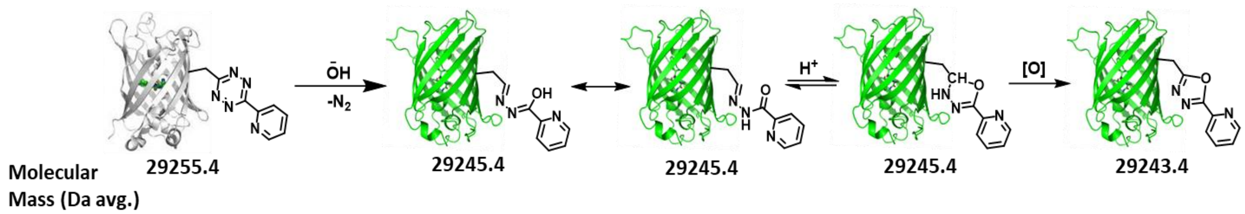

**Figure S21:** Proposed mechanism of Tet4-Pyr side reaction.

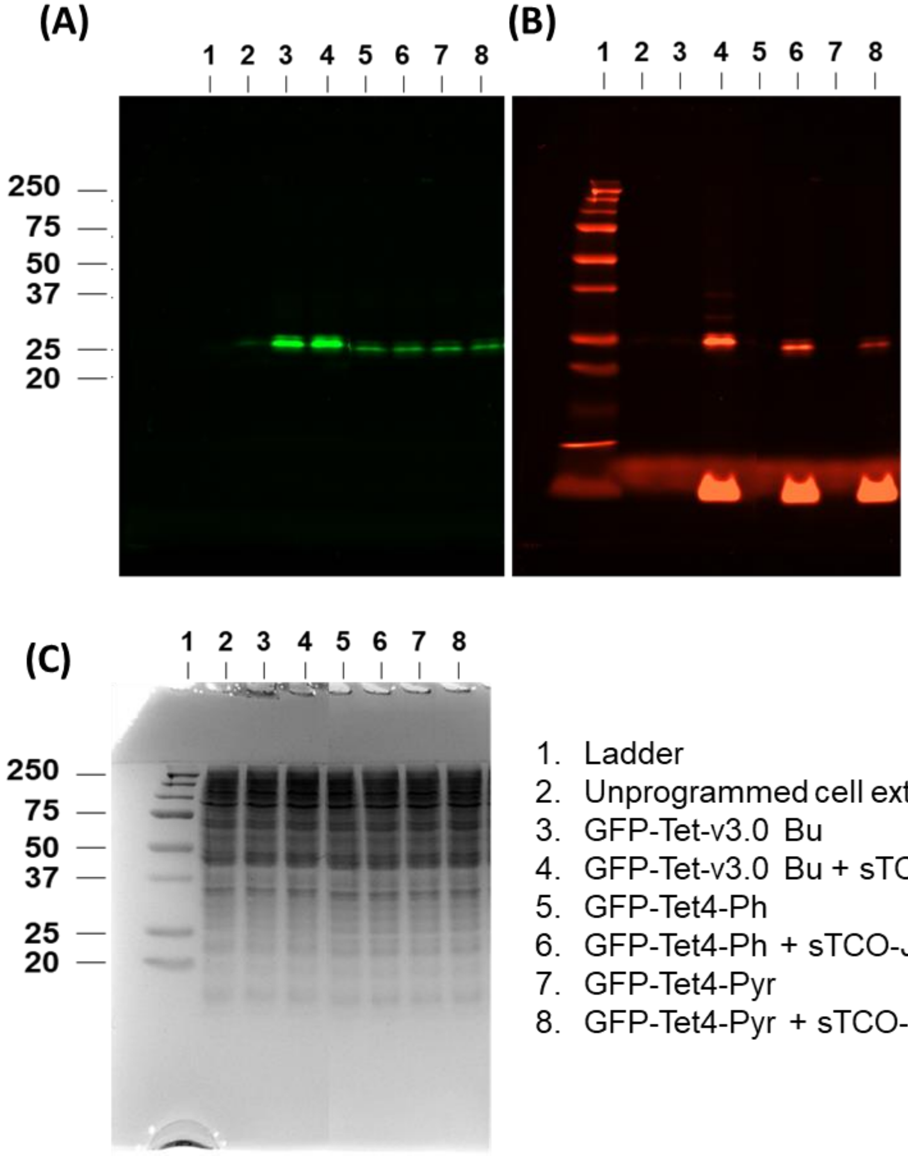

1. Ladder
2. Unprogrammed cell extract
3. GFP-Tet-v3.0 Bu
4. GFP-Tet-v3.0 Bu + sTCO-JF646
5. GFP-Tet4-Ph
6. GFP-Tet4-Ph + sTCO-JF646
7. GFP-Tet4-Pyr
8. GFP-Tet4-Pyr + sTCO-JF646

**Figure S22**: ***In vivo* labeling of GFP-Tet protein with sTCO-JF646**. HEK293T cells were transfected as described in figure S19. Cells were washed three times with culture media and incubated for 30 min with 1 µM sTCO-JF646. Unreacted labeling agent was quenched by excess Tet-v2.0Me for 10 min. Cell lysates were prepared in non-reducing Laemmli buffer and analyzed by SDS-PAGE. The GFP (A) and JF646 (B) fluorescence gel images were scanned on Bio-Rad ChemiDoc imaging system. The gel was stained with Coomassie Blue to visualize the protein (C).

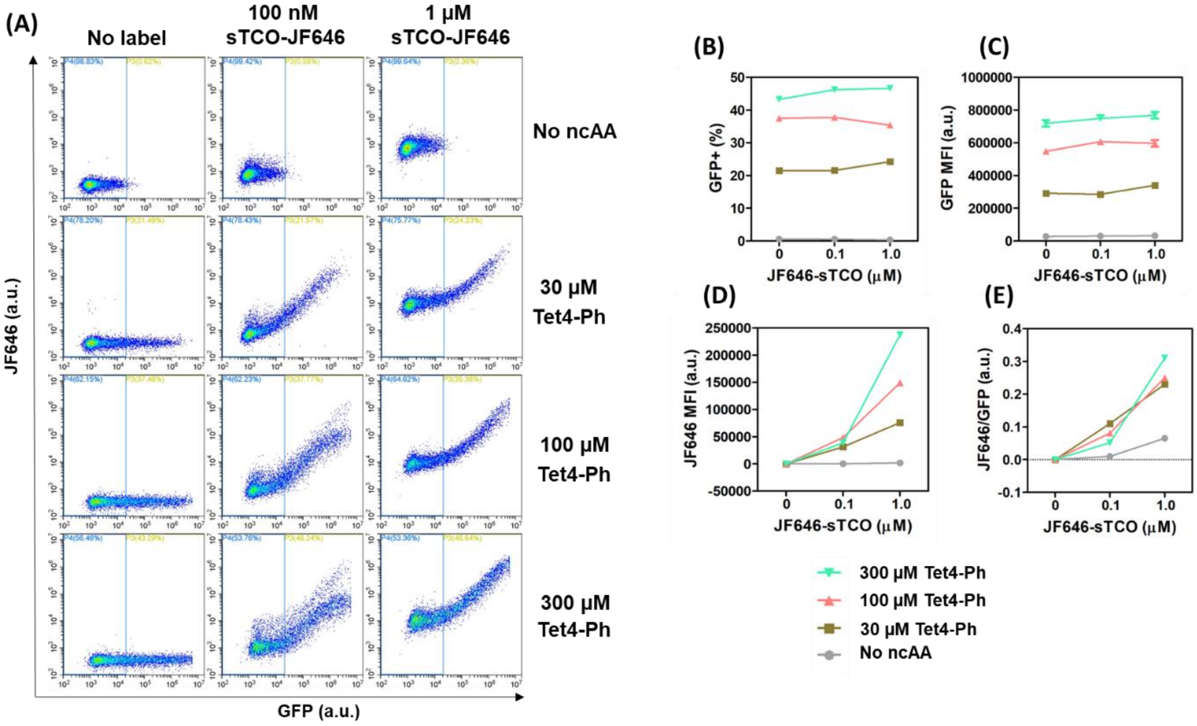

**Figure S23**: **Flow cytometry analysis of *in cellulo* labeling of Tet4-Ph protein**. HEK293T cells were transfected with pAcBac1-GFP-150TAG and pAcBac1-NES-E1-RS for 24 h in the presence of Tet4-Ph. Cells were washed three times and labeled with 100 nM sTCO-JF646, 1 uM sTCO-JF646, or 0.1% DMSO for 30 min at 37 $^{o}$C. The unreacted label was quenched by 3 µM Tet-v2.0Me for 10 min. Cells were washed once with PBS and analyzed by flow cytometry (A). A threshold (blue line in panel A) was arbitrarily set to split the cell population into two: GFP-negative and GFP-positive. The GFP-positive population is the one that has GFP fluorescence greater than the arbitrary threshold (A). The GFP-positive cell population increased as more Tet4- was added to the cells (B). The GFP mean fluorescence intensity (MFI) increased as more Tet4-Ph was added (C). The specific labeling was calculated by subtracting the background JF646 staining of GFP-negative population from JF646 staining of GFP-positive population The specific labeling was proportional to GFP MFI when labeled with 1 uM sTCO-JF646 (D).  The specific labeling was normalized by GFP MFI (E).

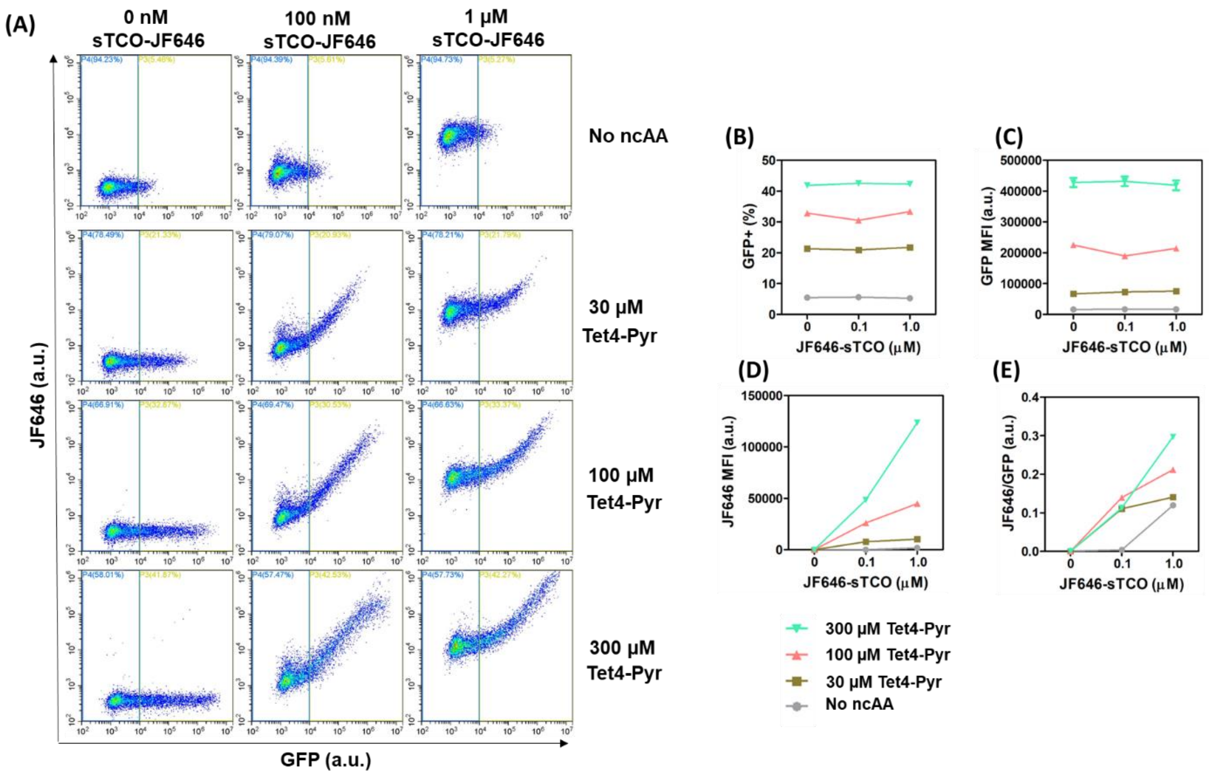

**Figure S24**: **Flow cytometry analysis of *in celluo* labeling of Tet4-Pyr protein**. HEK293T cells were transfected with pAcBac1-GFP-150TAG and pAcBac1-NES-D4 RS for 24 h in the presence of Tet4-Pyr. Cells were labeled and analyzed by flow cytometry as described in Figure S23 (A). The GFP-positive population (B) and the average GFP expression level per cell (C) increased as more Tet4-Pyr was added. The specific labeling was proportional to both GFP MFI and the label concentration (D). The specific labeling was normalized by GFP MFI (E).

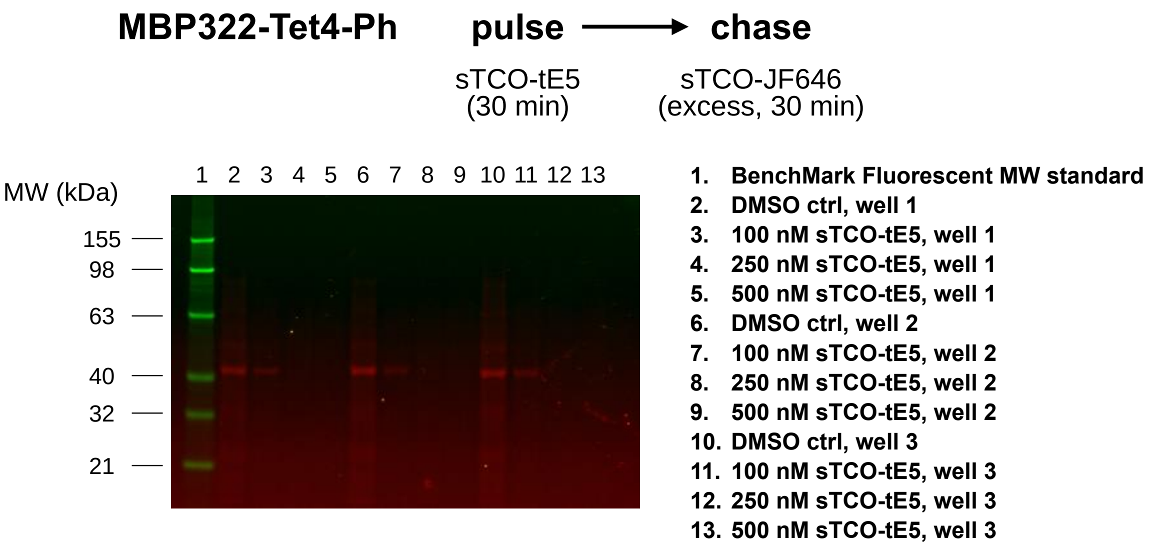

**Figure S25**: Spin-labeling of MBP322-Tet4-Ph in HEK293T cells measured by pulse-chase in-gel fluorescence assay. Each lane is from a separate well of HEK293T cells grown and transfected in 6-well plates. Washed cells were incubated with sTCO-tE5 at the specified concentration in DMEM + 10% FBS at room temperature for 30 minutes. Reactions were chased by addition of 0.5 μM sTCO-JF646 for 30 minutes at r.t. and quenched with 10 μM Tet4-Ph. Cells were lysed and non-spin-labeled protein was quantified by SDS-PAGE with JF-646 fluorescence detection.

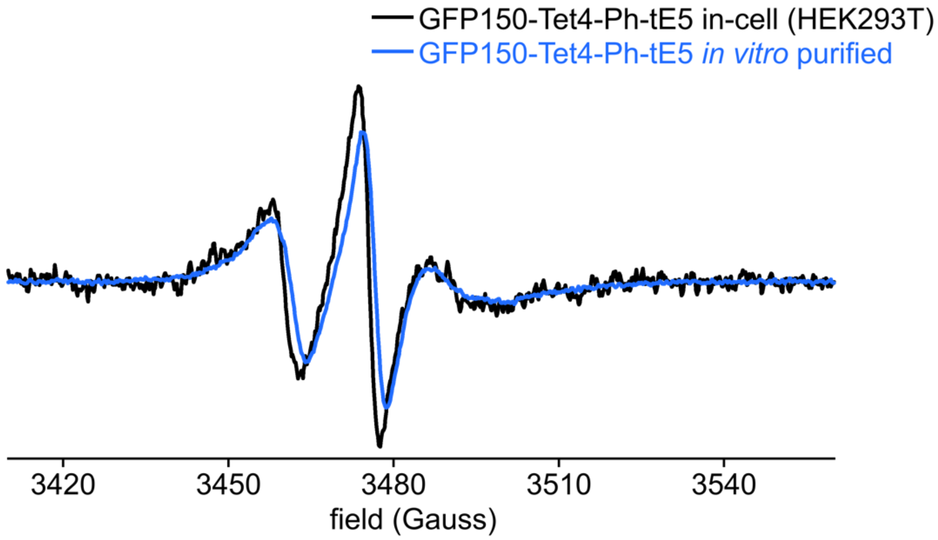

**Figure S26**: Room temperature X-band CW EPR spectra of GFP150-Tet4-Ph spin-labeled with sTCO-tE5 nitroxide. Similar lineshapes are observed for protein expressed, labeled, and measured directly in HEK293T cells (black) and protein purified and spin-labeled *in vitro* (blue). Spectra are normalized by total spin concentrations calculated by double integration of the baselined EPR signal.

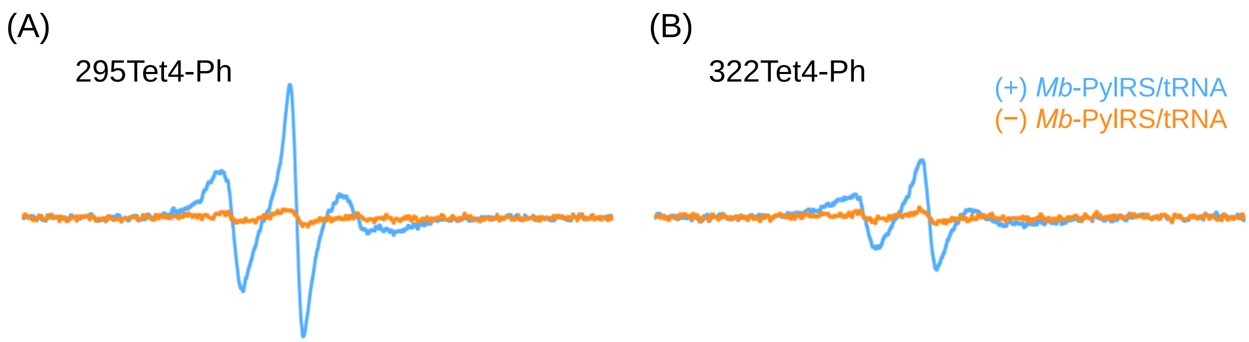

**Figure S27**: In-cell X-band CW EPR spectra (r.t.) of MBP295-Tet4-Ph (A) and MBP322-Tet4-Ph (B) constructs in HEK293T cells spin-labeled with 200 nM sTCO-tE5 in DMEM growh medium for 30 minutes (r.t.). *Blue* traces are spectra recorded from expresions containing all GCE components (MBP-TAG DNA, *Mb*-PylRS/tRNA-E1 DNA, and 100 µM Tet4 ncAA), whereas *orange* traces are from cells in which the plasmid encoding the orthogonal PylRS/tRNA pair was omitted.

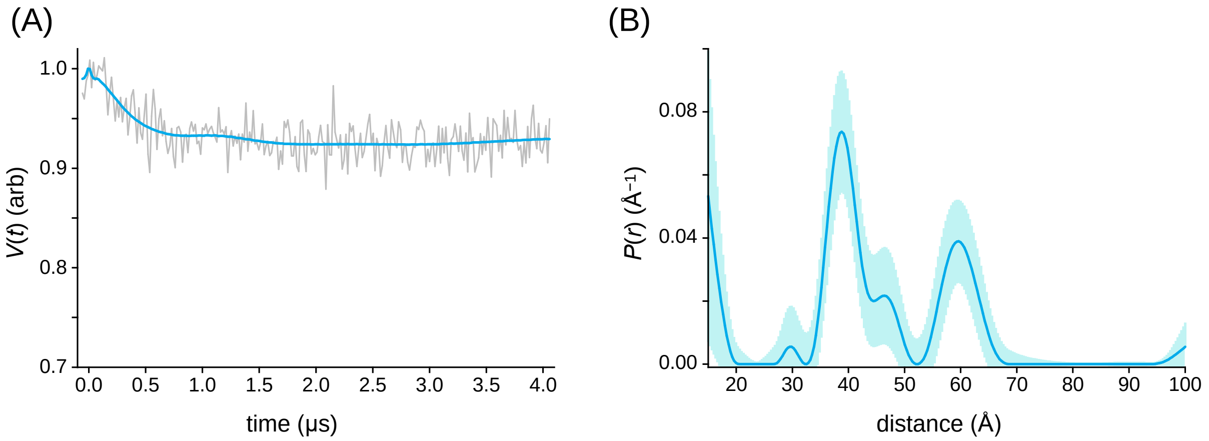

**Figure S28: In-cell DEER of MBP W340A mutant** (A) Background-corrected 4-pulse DEER time-domain data (Q-band, 45 K) and corresponding fit from HEK293T cells expressing MBP211/295(W340A)-Tet4-Ph and spin-labeled with sTCO-tE5. (B) Calculated distance distribution obtained by non-parametric fitting of the time-domain signal with Tikhonov regularization using the LongDistances software package. Error bands on the distribution were generated by the Error function of LongDistances using default values.

**$^{1}H$ and $^{13}C$ NMR spectra of synthesized compounds**:

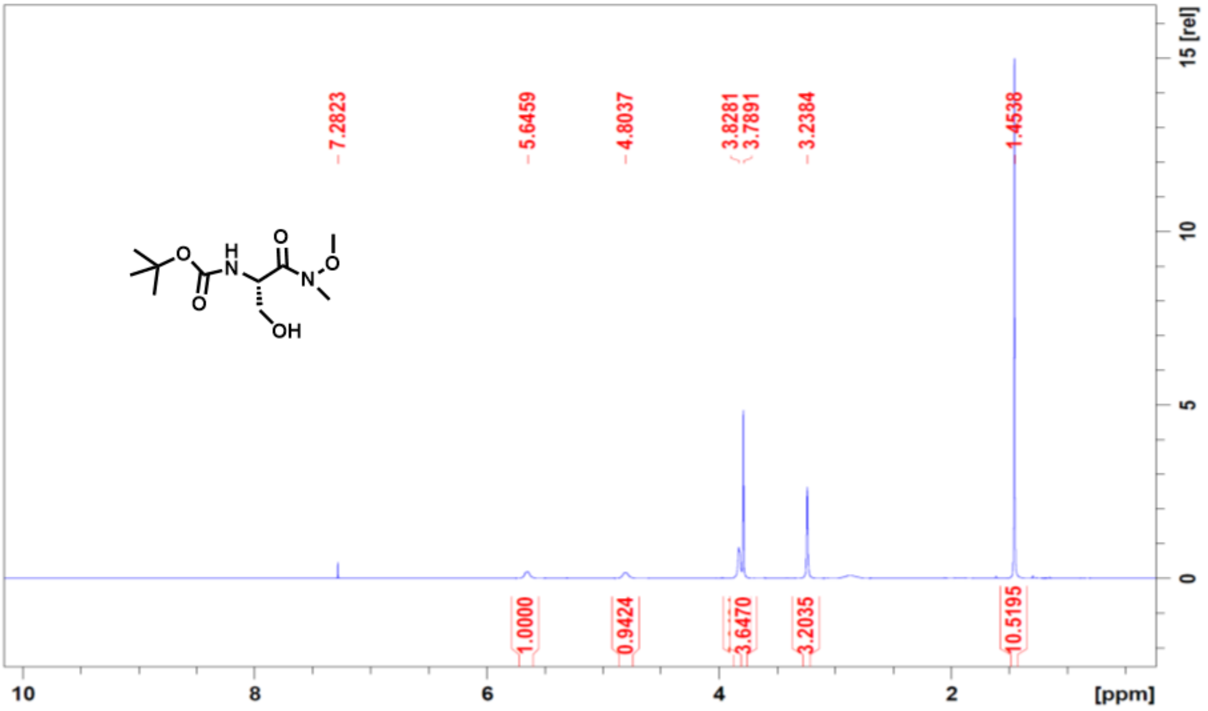

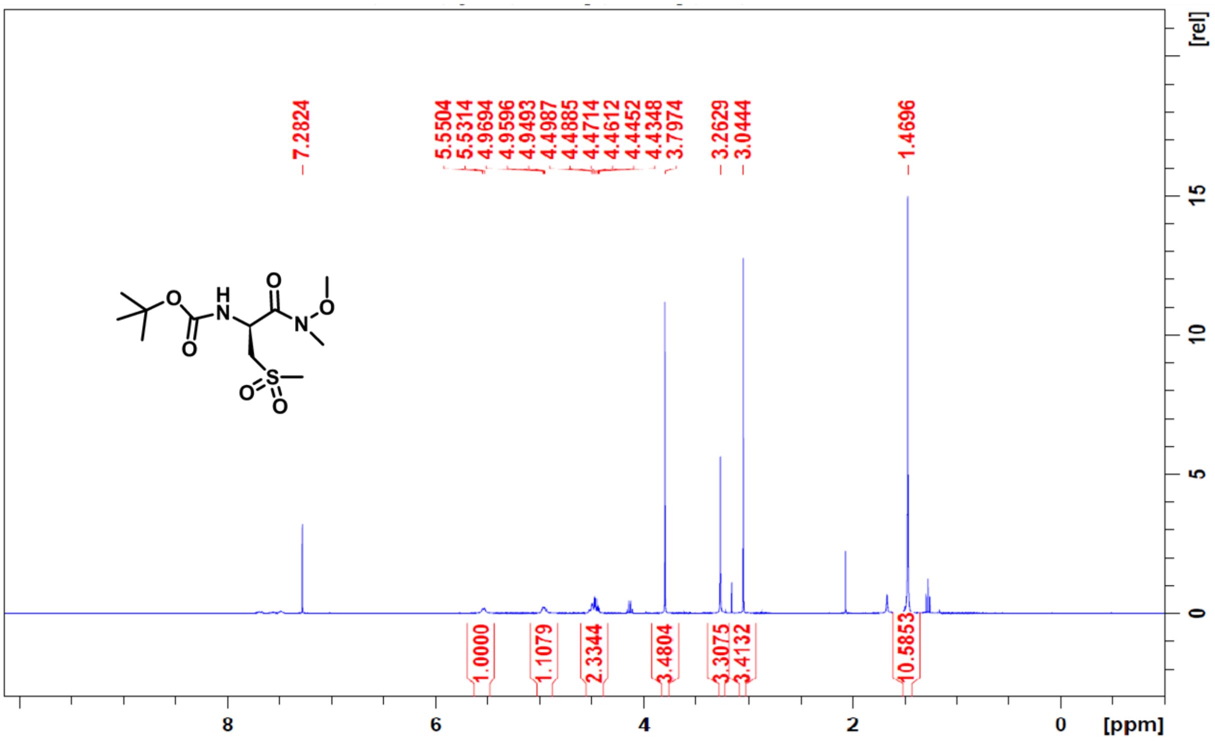

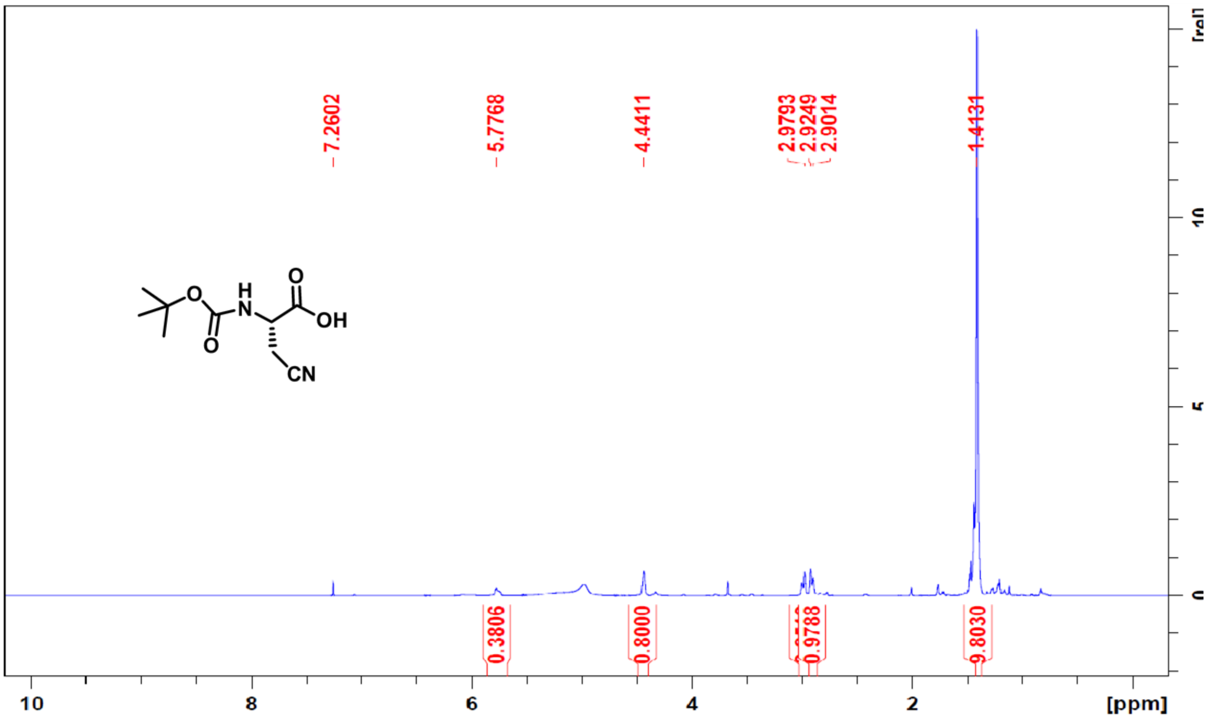
7.2602
5.7768
4.4411
2.9793
2.9249
2.9014
1.4131
0.3806
0.8000
0.9788
9.8030
10
8
6
4
2
[ppm]
O
H
N
O
OH
O
CN

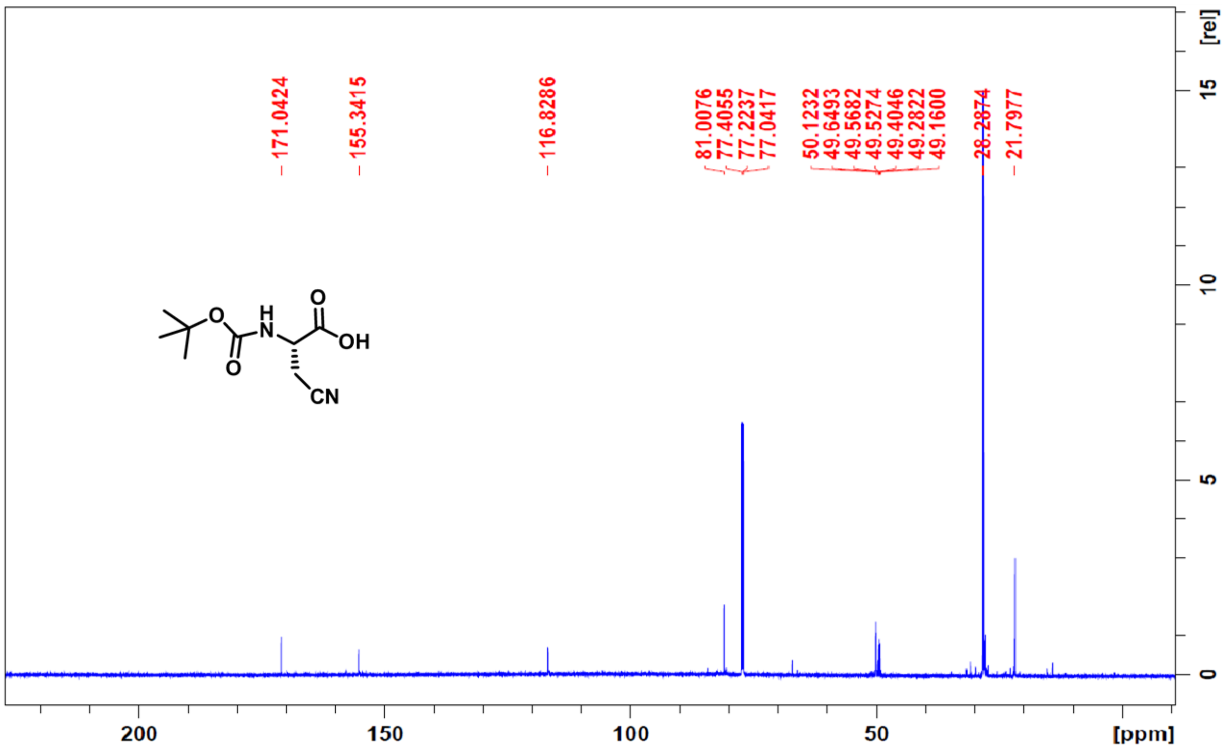
171.0424
155.3415
116.8286
81.0076
77.4055
77.2237
77.0417
50.1232
49.6493
49.5682
49.5274
49.4046
49.2822
49.1600
28.2874
21.7977
[rel]
15
10
5
0
200
150
100
50
[ppm]
O
H
N
O
OH
O
CN

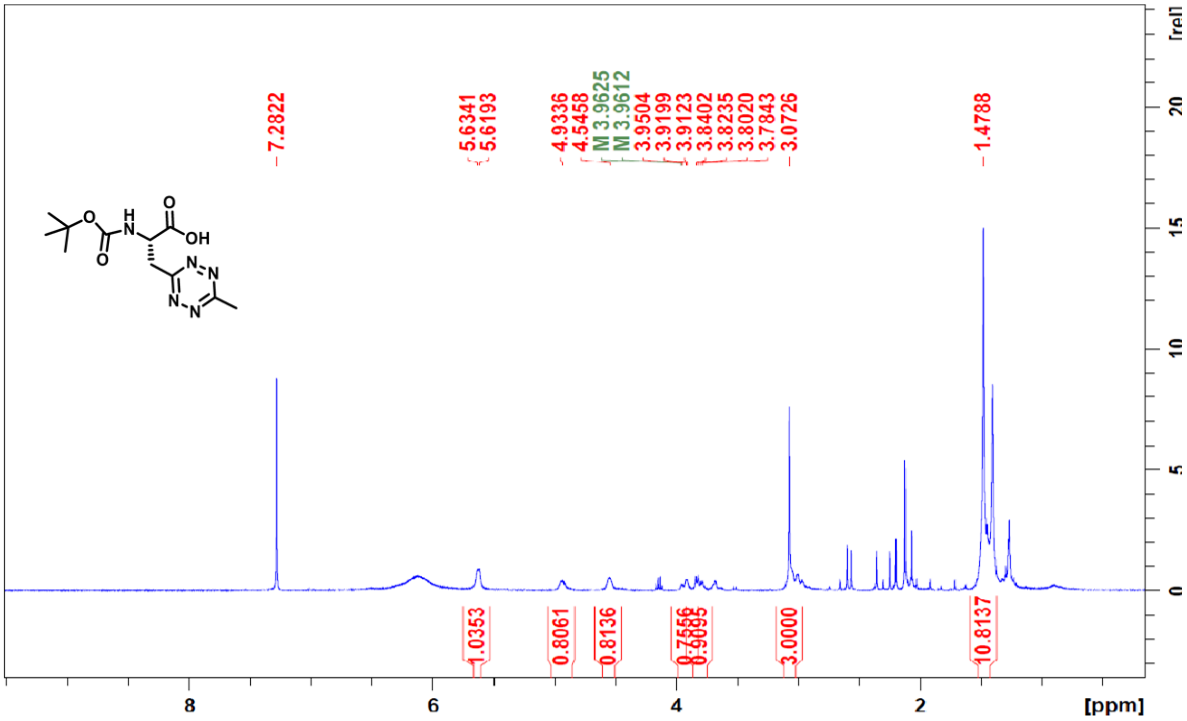
7.2822
5.6341
5.6193
4.9336
4.5458
M 3.9625
M 3.9612
3.9504
3.9199
3.9123
3.8402
3.8235
3.8020
3.7843
3.0726
1.4788
1.0353
0.8061
0.8136
0.7556
0.9095
3.0000
10.8137
[rel]
[ppm]

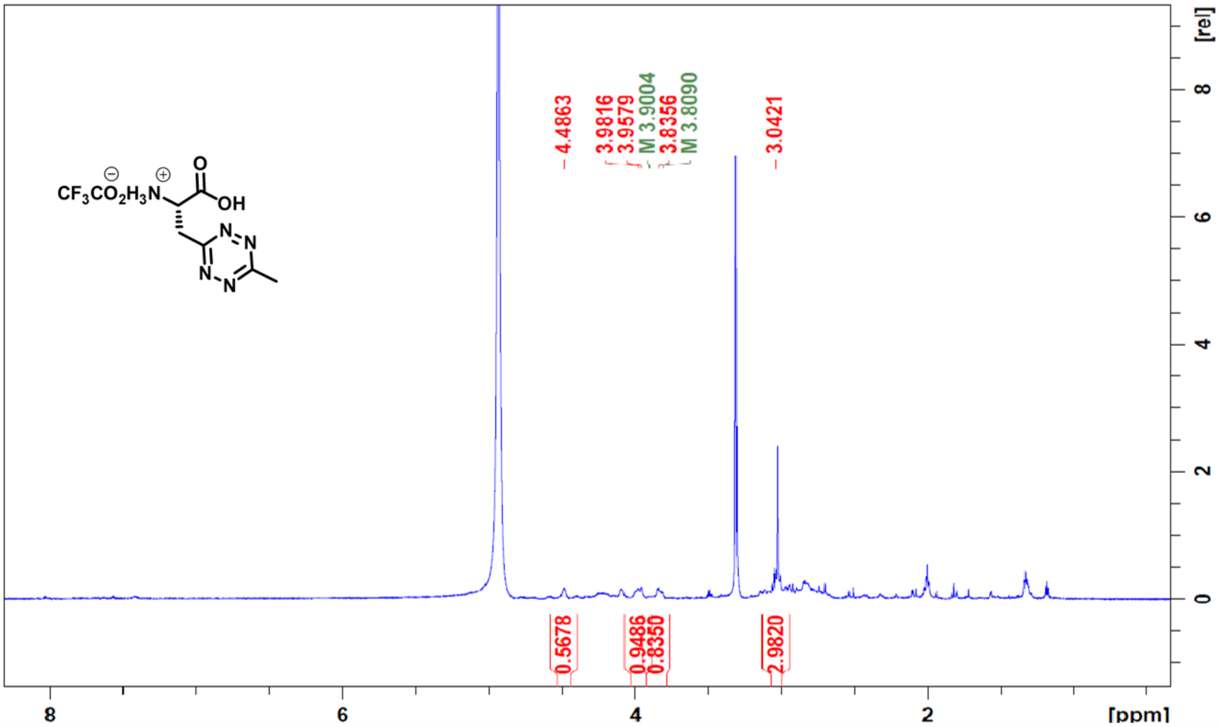
4.4863
3.9816
3.9579
M 3.9004
3.8356
M 3.8090
3.0421
0.5678
0.9486
0.8350
2.9820
[rel]
[ppm]

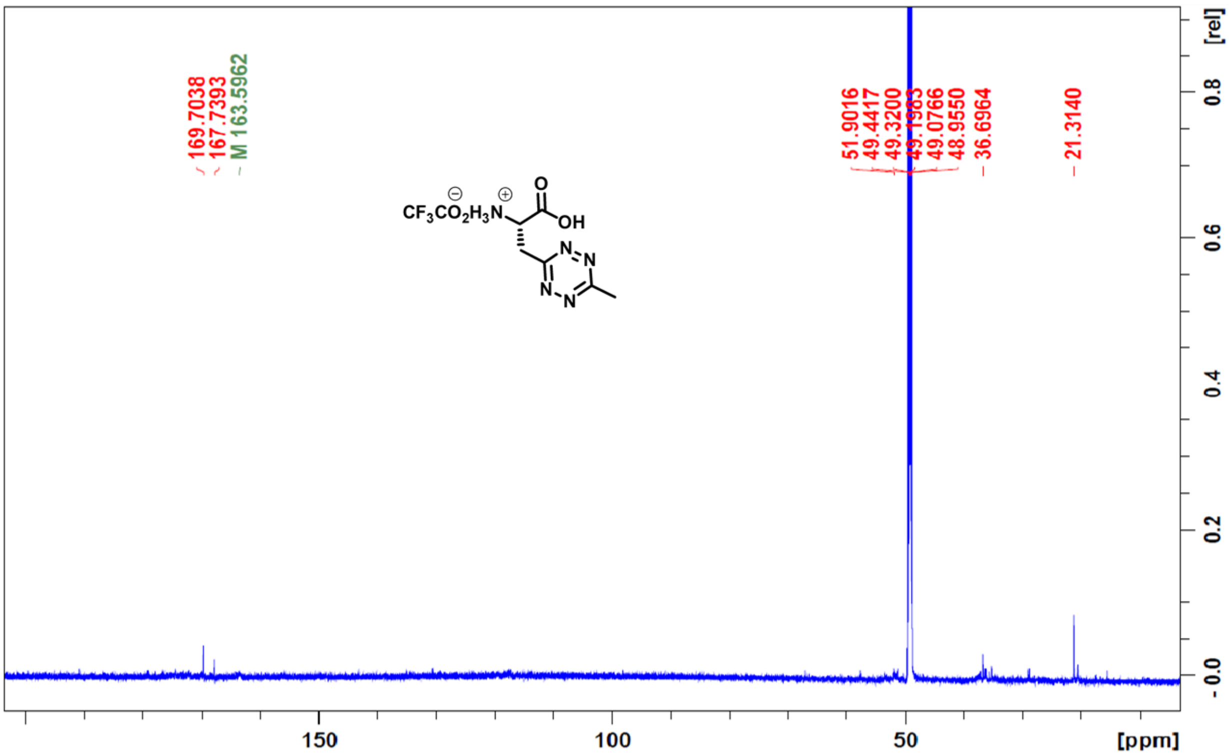
169.7038
167.7393
M 163.5962
51.9016
49.4417
49.3200
49.1983
49.0766
48.9550
36.6964
21.3140
[rel]
0.8
0.6
0.4
0.2
-0.0
150
100
50
[ppm]

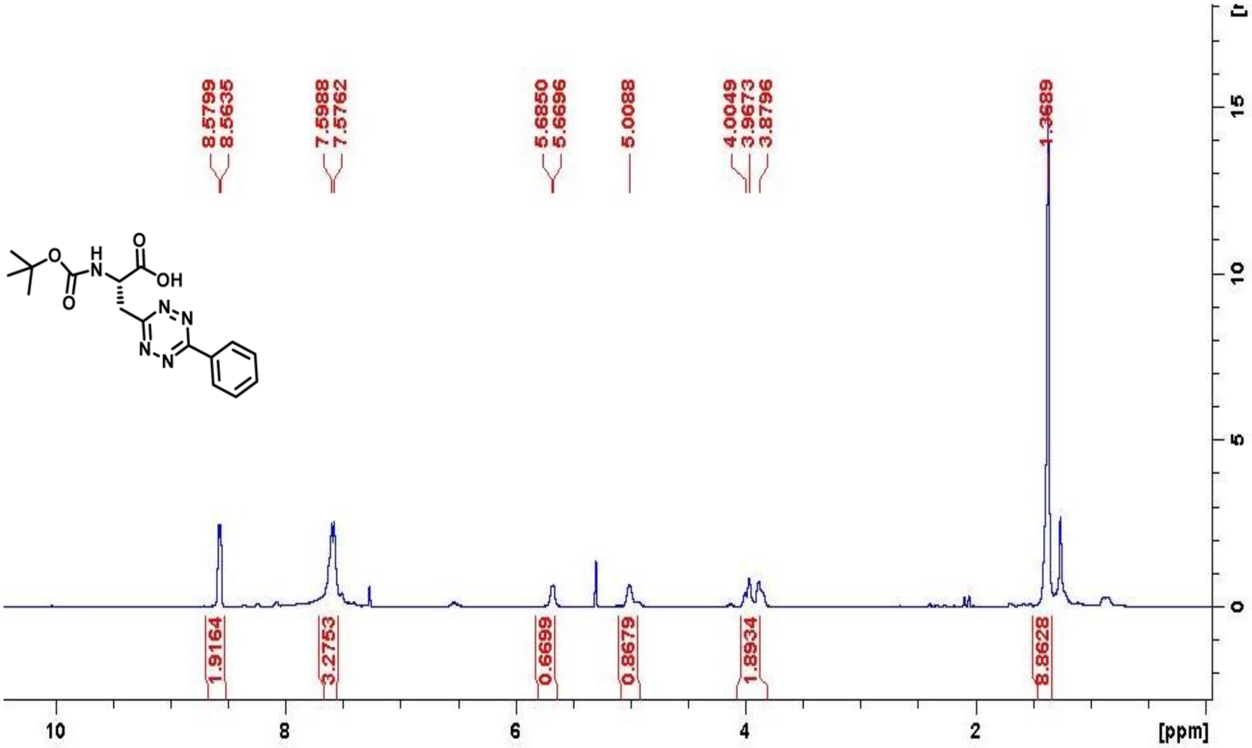
8.5799
8.5635
7.5988
7.5762
5.6850
5.6696
5.0088
4.0049
3.9673
3.8796
1.3689
1.9164
3.2753
0.6699
0.8679
1.8934
8.8628
[ppm]

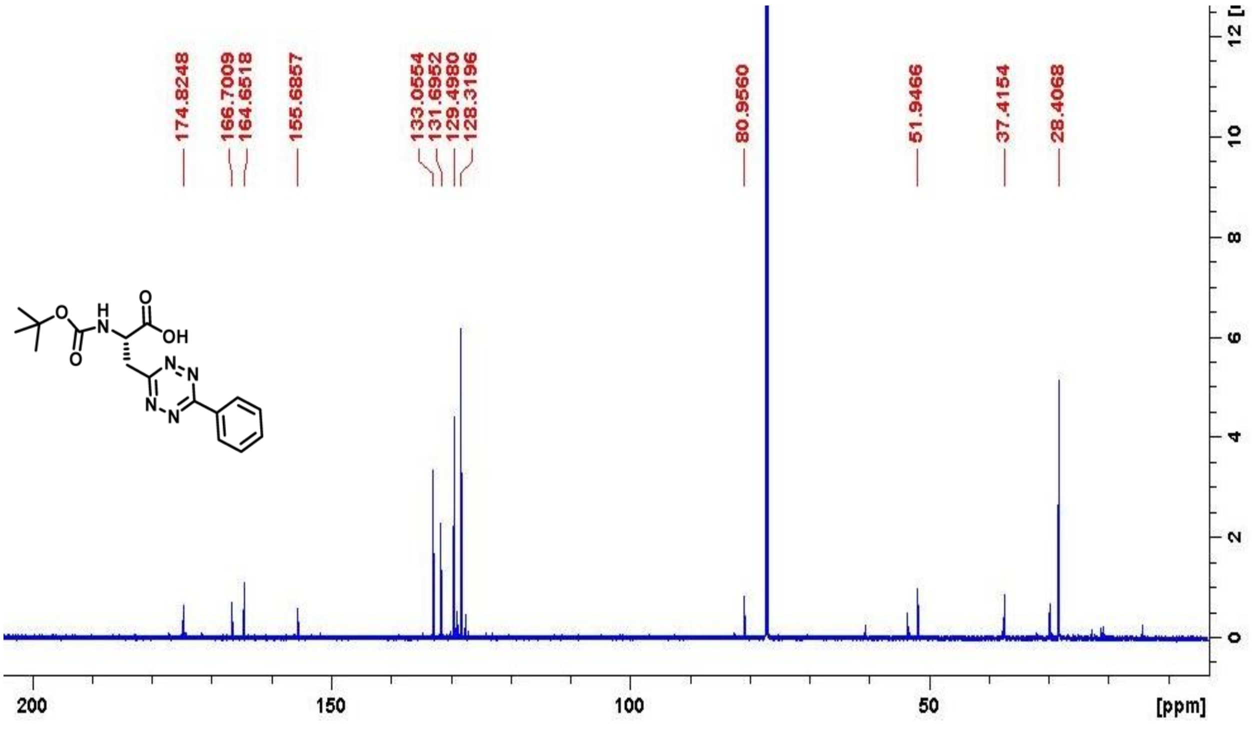
174.8248
166.7009
164.6518
155.6857
133.0554
131.6952
129.4980
128.3196
80.9560
51.9466
37.4154
28.4068
[ppm]

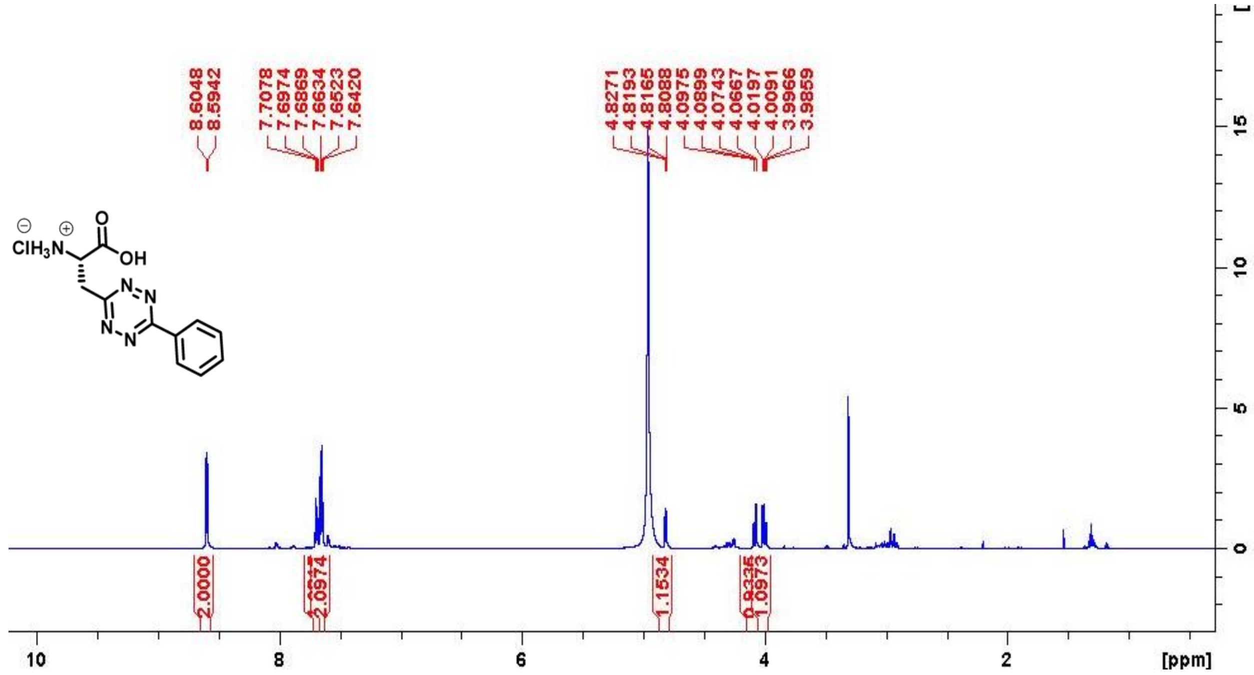

8.6048
8.5942
7.7078
7.6974
7.6869
7.6634
7.6523
7.6420
4.8271
4.8193
4.8165
4.8088
4.0975
4.0899
4.0743
4.0667
4.0197
4.0091
3.9966
3.9859
2.0000
2.0974
1.1534
1.0973
10
8
6
4
2
[ppm]

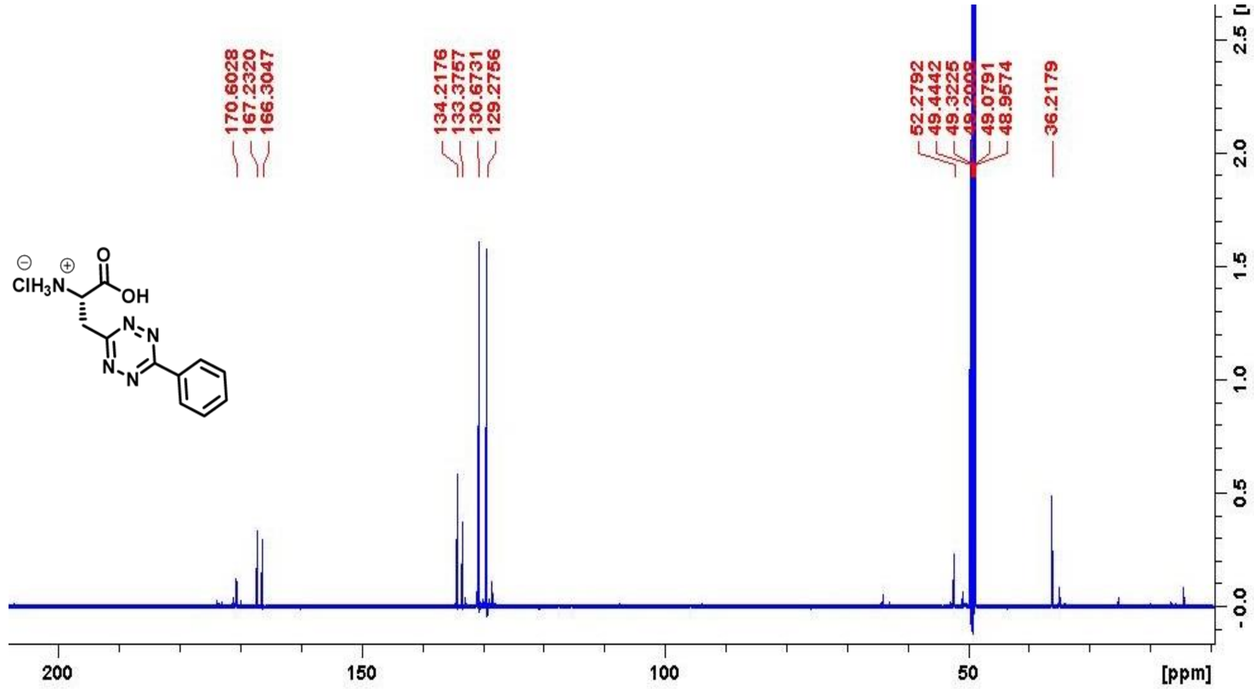

170.6028
167.2320
166.3047
134.2176
133.3757
130.6731
129.2756
52.2792
49.4442
49.3225
49.0791
48.9574
36.2179
200
150
100
50
[ppm]

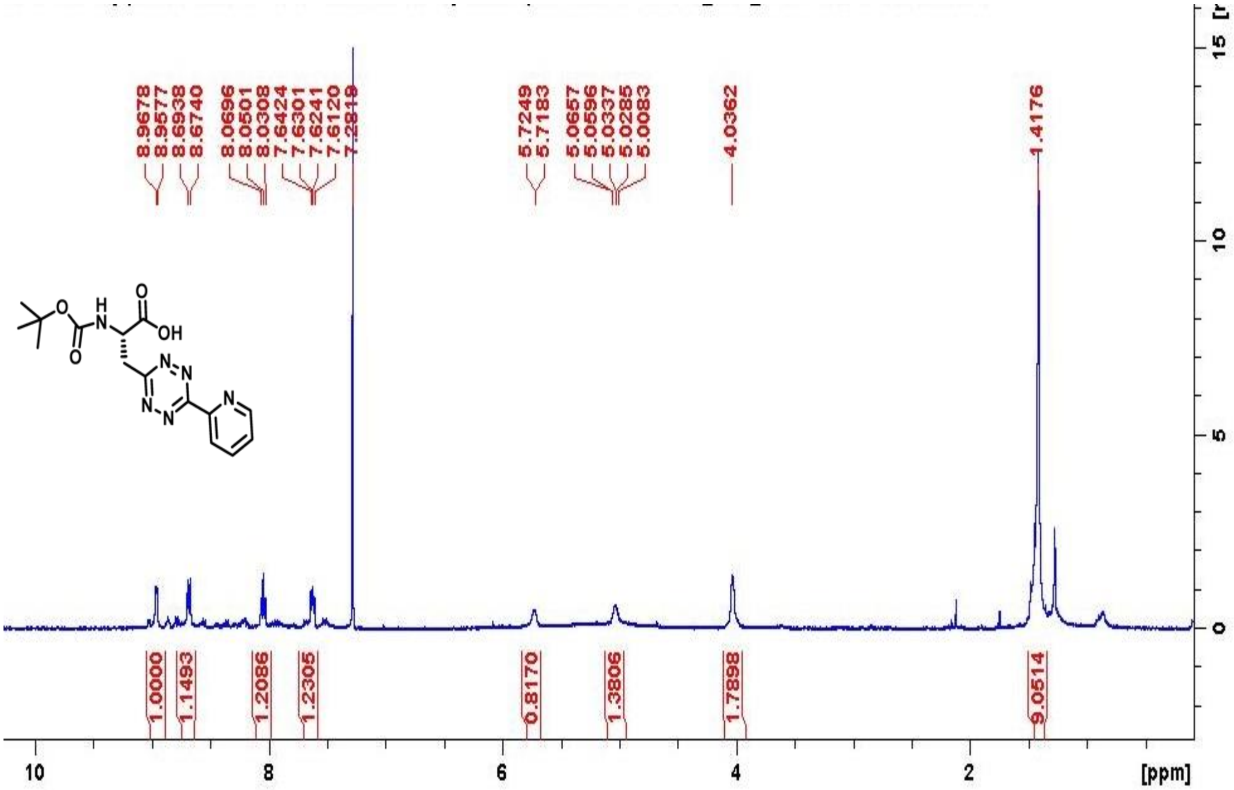

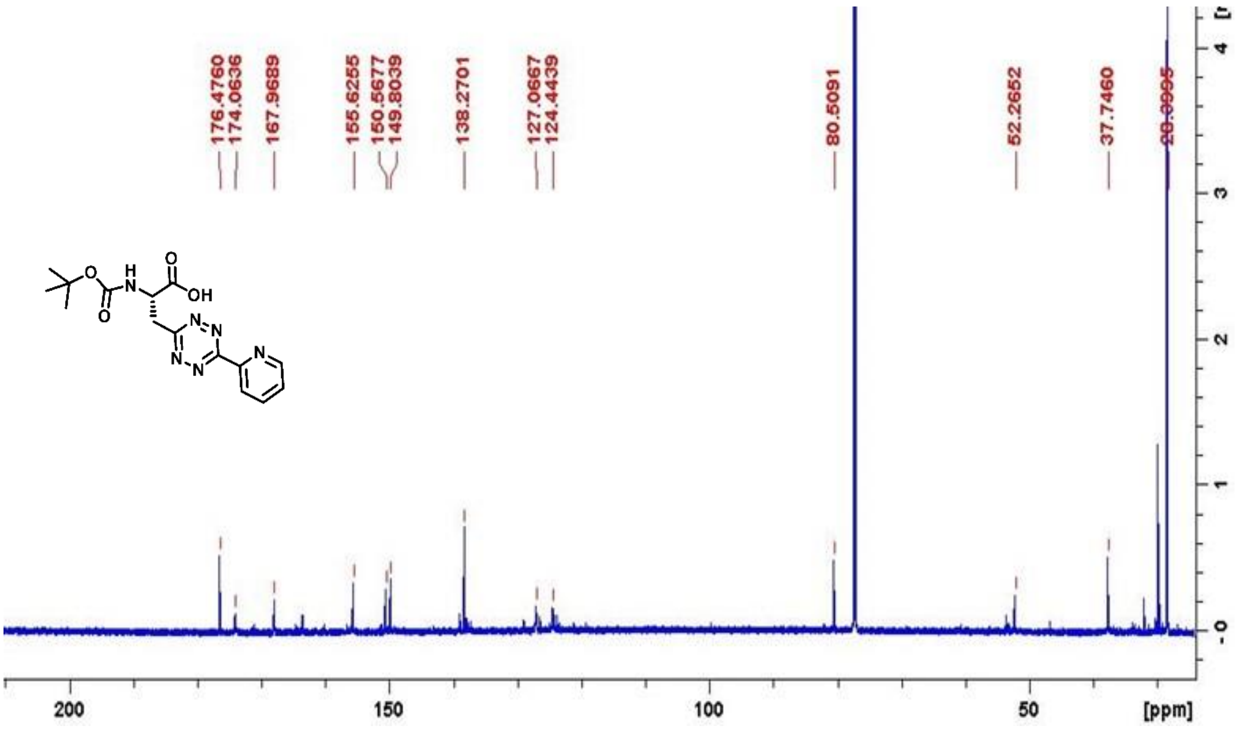

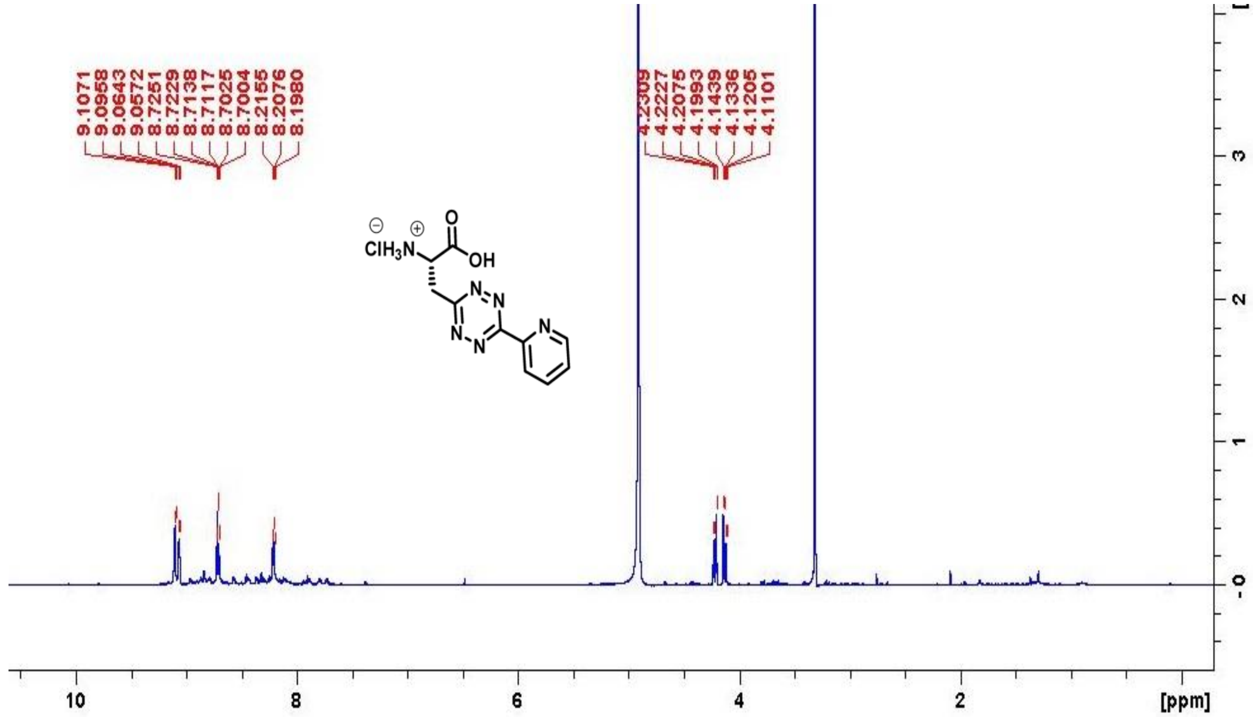

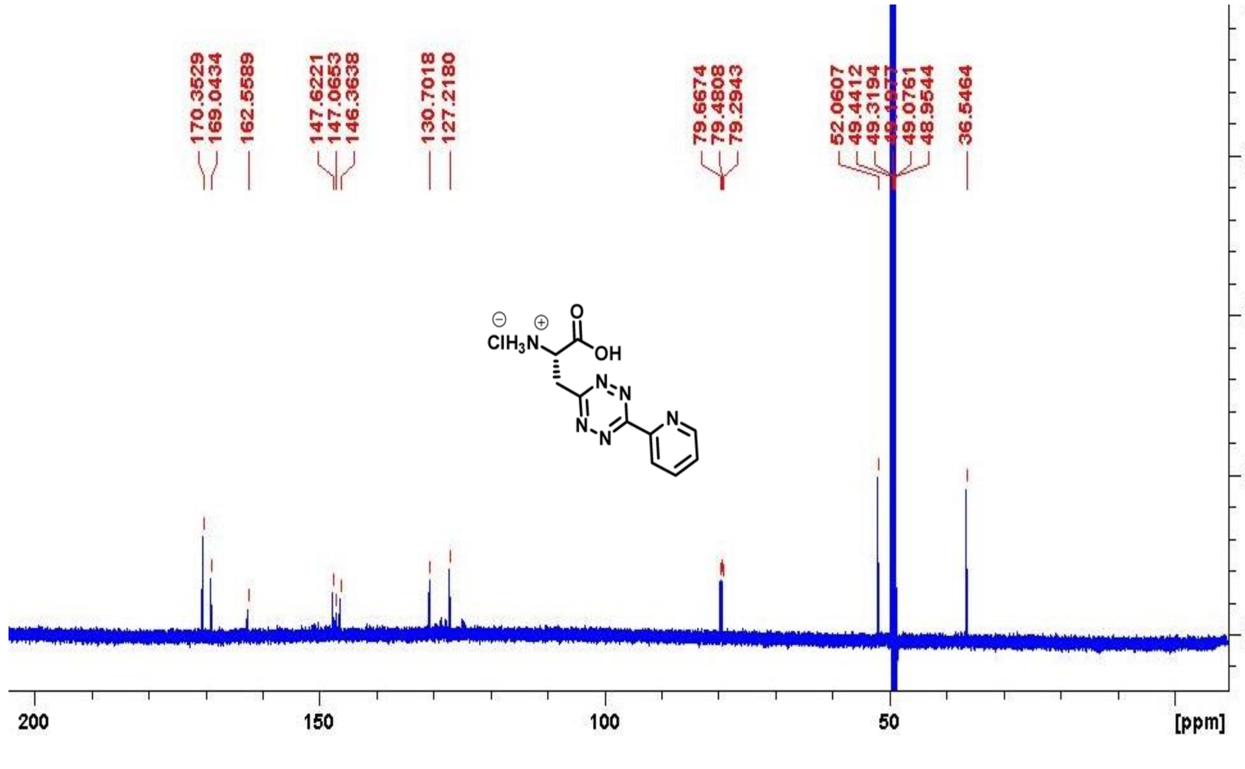

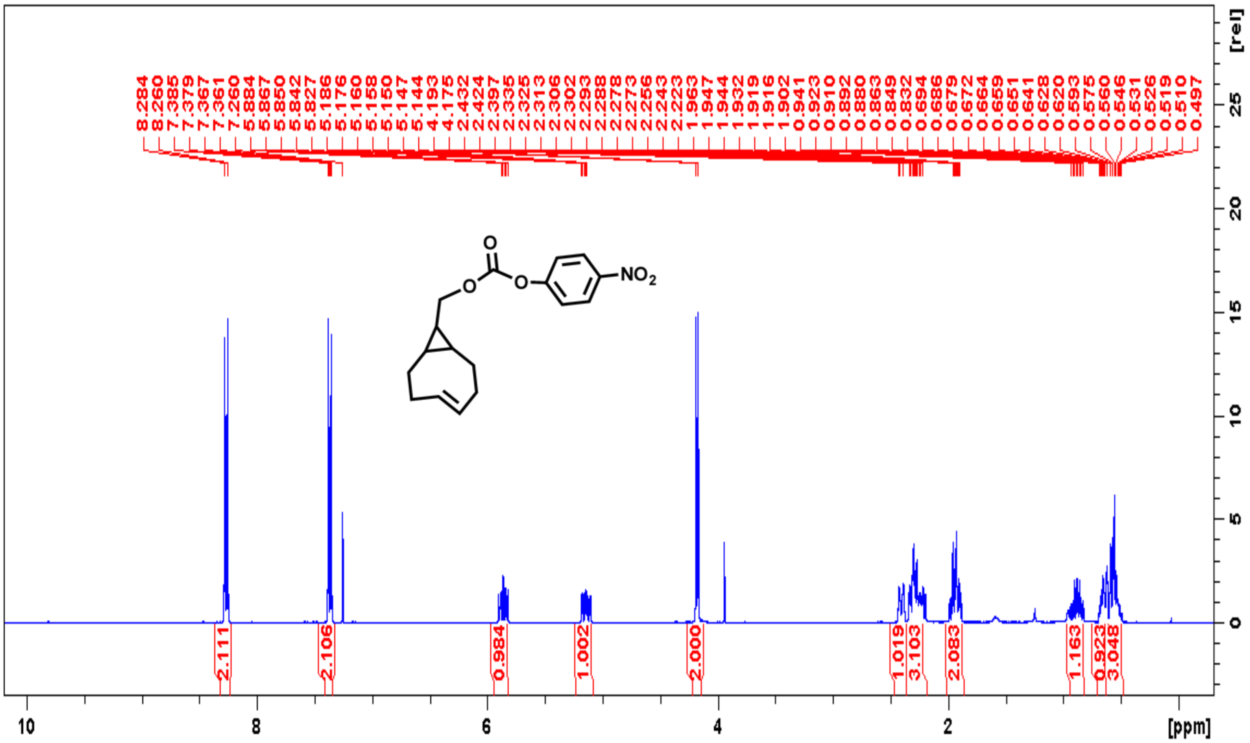
8.284 8.260 7.385 7.379 7.367 7.361 7.260 5.884 5.867 5.850 5.842 5.827 5.186 5.176 5.160 5.158 5.150 5.144 4.193 4.175 2.432 2.424 2.397 2.335 2.325 2.313 2.306 2.302 2.293 2.288 2.278 2.273 2.256 2.243 2.223 1.963 1.947 1.944 1.932 1.919 1.916 1.902 0.941 0.923 0.910 0.892 0.880 0.863 0.849 0.832 0.694 0.686 0.679 0.672 0.664 0.659 0.651 0.641 0.628 0.620 0.593 0.575 0.560 0.546 0.531 0.526 0.519 0.510 0.497
NO2
2.111 2.106 0.984 1.002 2.000 1.019 3.103 2.083 1.163 0.923 3.048
[rel]
[ppm]

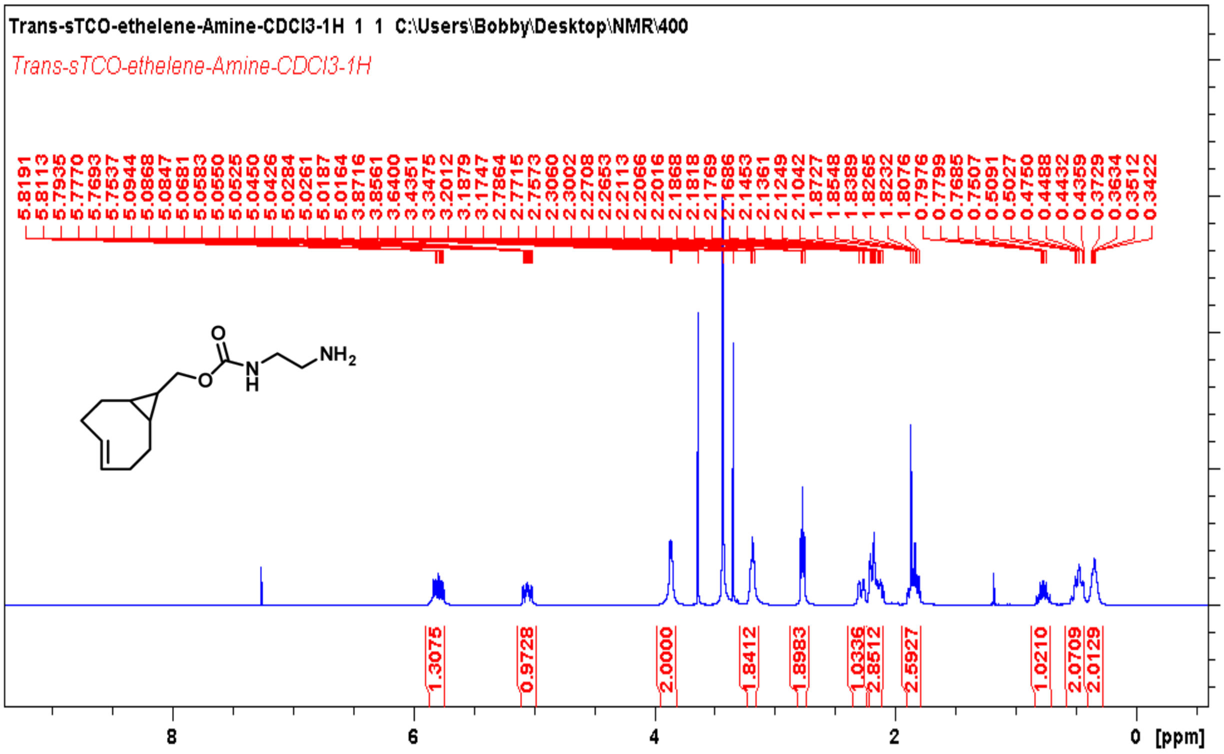
Trans-sTCO-ethelene-Amine-CDCl3-1H 1 1 C:\Users\Bobby\Desktop\NMR\400
Trans-sTCO-ethelene-Amine-CDCl3-1H
5.8191 5.8113 5.7935 5.7770 5.7693 5.7537 5.0944 5.0868 5.0847 5.0681 5.0583 5.0550 5.0525 5.0450 5.0426 5.0284 5.0261 5.0187 5.0164 3.8716 3.8561 3.6400 3.4351 3.3475 3.2012 3.1879 3.1747 2.7864 2.7715 2.7573 2.3060 2.3002 2.2708 2.2653 2.2113 2.2066 2.2016 2.1868 2.1818 2.1769 2.1686 2.1453 2.1361 2.1249 2.1042 1.8727 1.8548 1.8389 1.8265 1.8232 1.8076 0.7976 0.7799 0.7685 0.7507 0.5091 0.5027 0.4750 0.4488 0.4432 0.4359 0.3729 0.3634 0.3512 0.3422
NH2
1.3075 0.9728 2.0000 1.8412 1.8983 1.0336 2.8512 2.5927 1.0210 2.0709 2.0129
[rel]
[ppm]

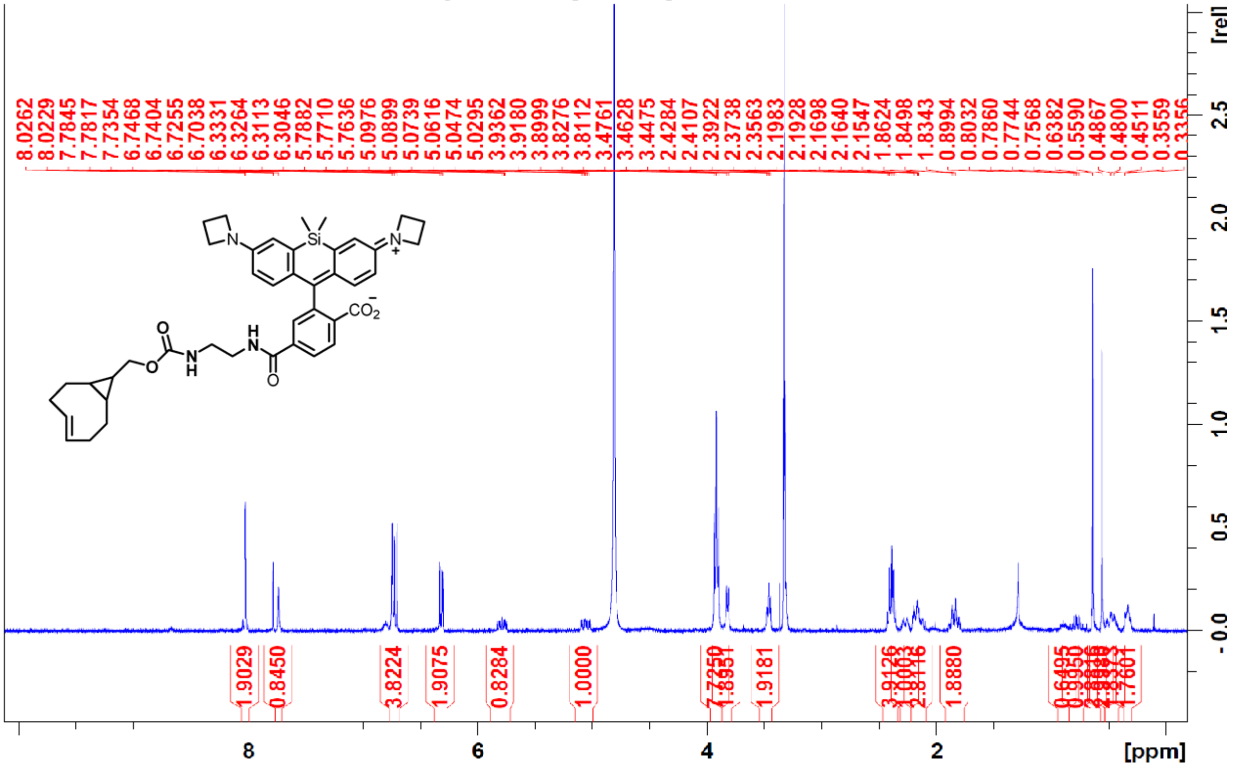
8.0262
8.0229
7.7845
7.7817
7.7354
6.7468
6.7404
6.7255
6.7038
6.3331
6.3264
6.3113
6.3046
5.7882
5.7710
5.7636
5.0976
5.0899
5.0739
5.0616
5.0474
5.0295
3.9362
3.9180
3.8999
3.8276
3.8112
3.4761
3.4628
3.4475
2.4284
2.4107
2.3922
2.3738
2.3563
2.1983
2.1928
2.1698
2.1640
2.1547
1.8624
1.8498
1.8343
0.8994
0.8032
0.7860
0.7744
0.7568
0.6382
0.5590
0.4867
0.4800
0.4511
0.3559
1.9029
0.8450
3.8224
1.9075
0.8284
1.0000
1.9181
1.8880
[ppm]
[rel]
Si
N
O
H

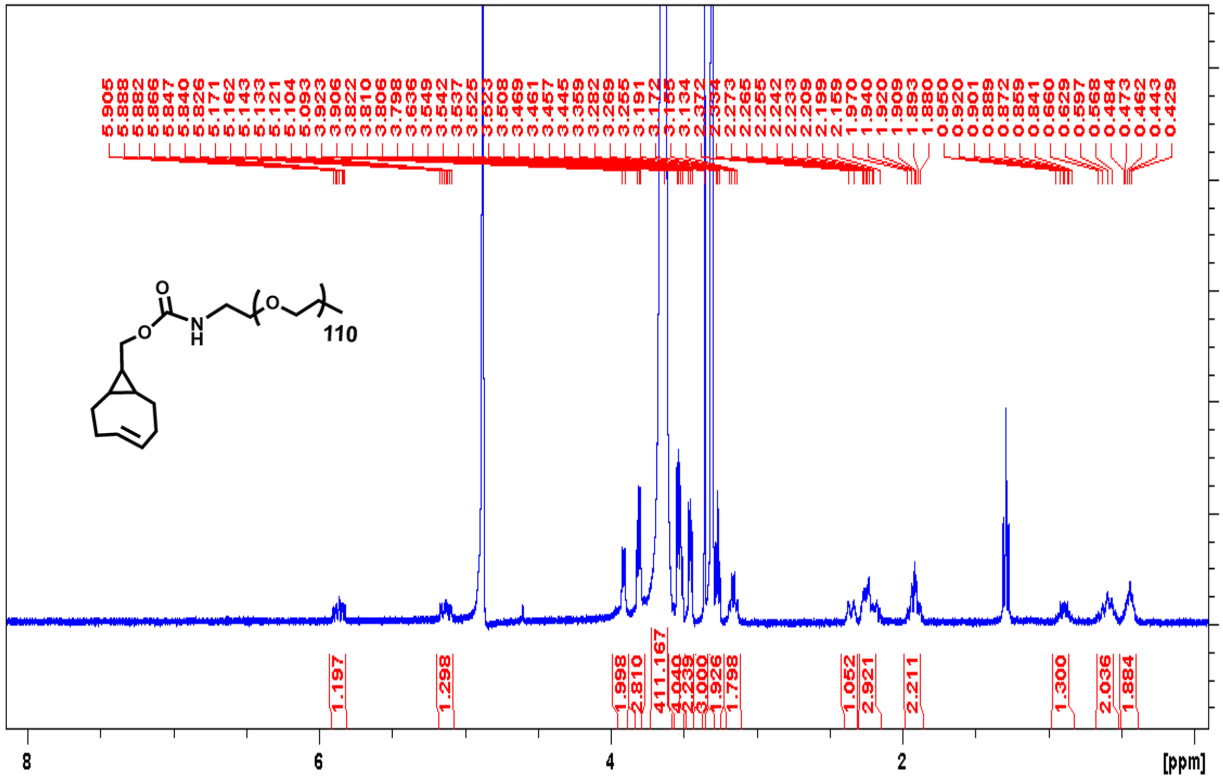
5.905
5.888
5.882
5.866
5.847
5.840
5.826
5.171
5.162
5.143
5.121
5.104
5.093
3.923
3.906
3.822
3.810
3.806
3.798
3.636
3.549
3.542
3.537
3.525
3.508
3.469
3.461
3.457
3.445
3.359
3.282
3.269
3.255
3.191
3.172
3.155
3.134
2.273
2.265
2.255
2.242
2.233
2.209
2.199
2.159
1.970
1.940
1.920
1.909
1.893
1.880
0.950
0.920
0.901
0.889
0.872
0.859
0.841
0.660
0.629
0.597
0.568
0.484
0.473
0.462
0.443
0.429
1.197
1.298
1.998
2.810
411.167
4.040
2.239
3.000
1.926
1.798
1.052
2.921
2.211
1.300
2.036
1.884
[ppm]
[rel]
110
O
N
H

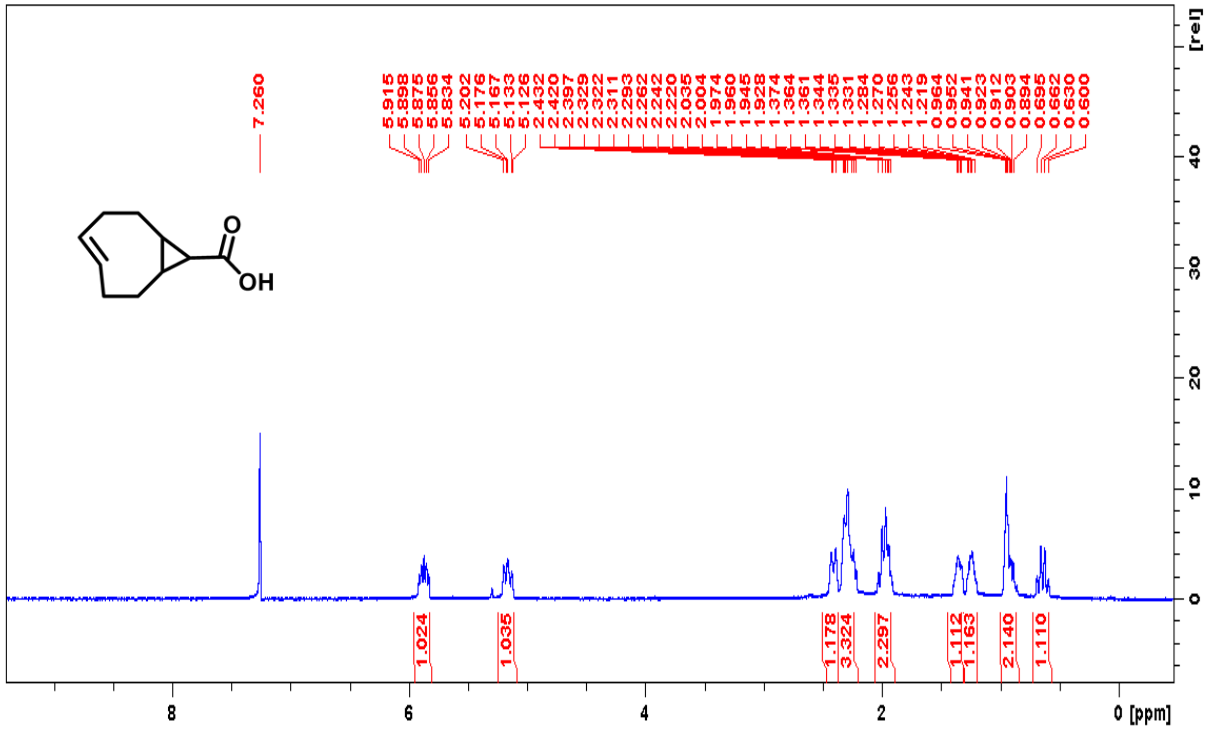

O
OH
7.260
5.915
5.898
5.875
5.856
5.834
5.202
5.176
5.167
5.133
5.126
2.432
2.420
2.397
2.329
2.322
2.311
2.293
2.262
2.242
2.220
2.035
2.004
1.974
1.960
1.945
1.928
1.374
1.364
1.361
1.344
1.335
1.331
1.284
1.270
1.256
1.243
1.219
0.964
0.952
0.941
0.923
0.912
0.903
0.894
0.695
0.662
0.630
0.600
1.024
1.035
1.178
3.324
2.297
1.112
1.163
2.140
1.110
[rel]
8
6
4
2
0 [ppm]

**ESI-MS spectrum of synthesized sTCO-spin labels**

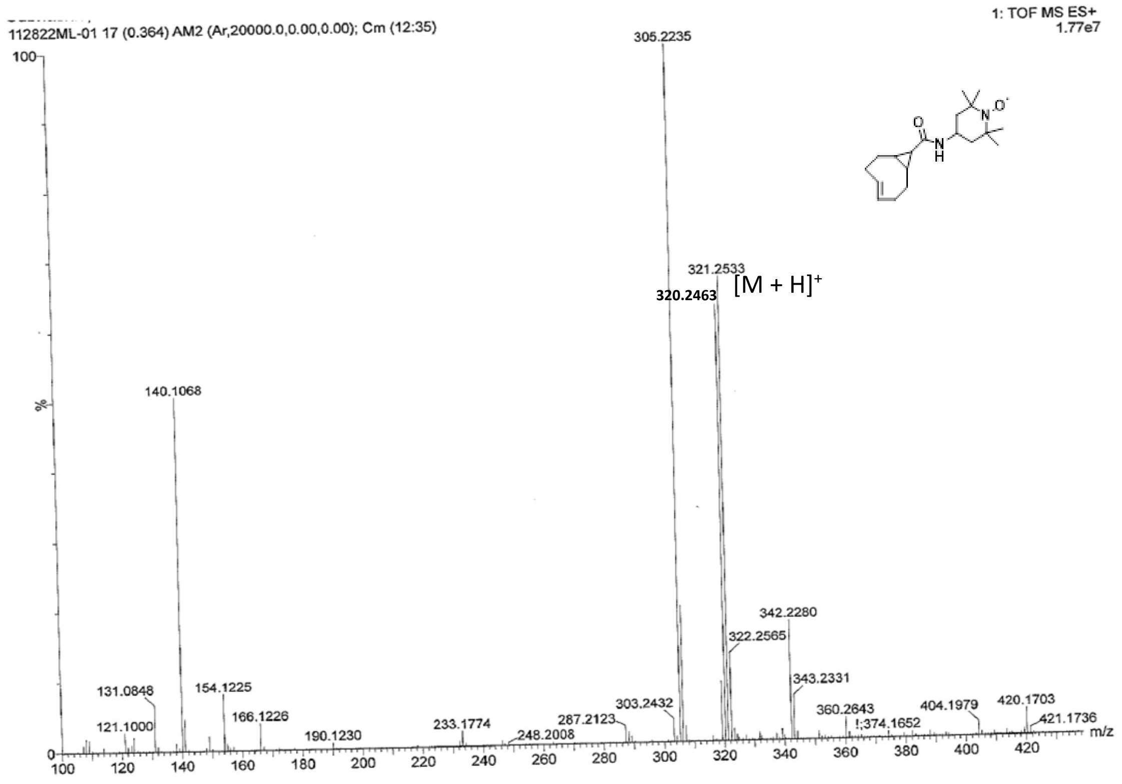

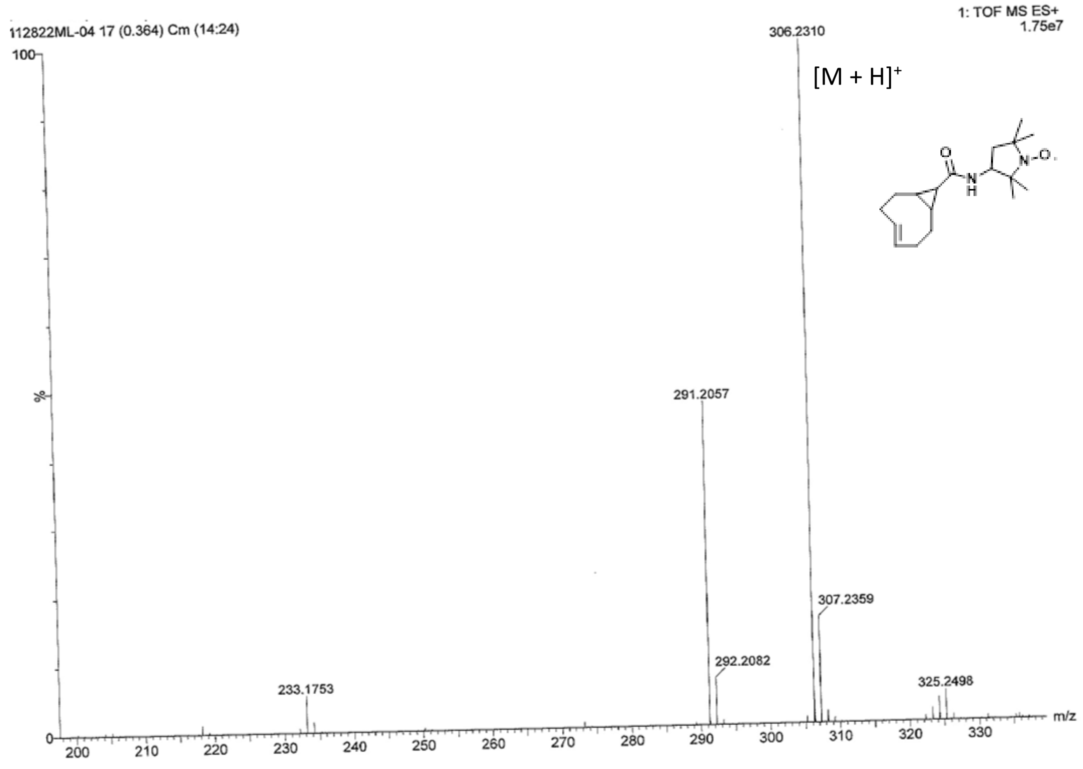

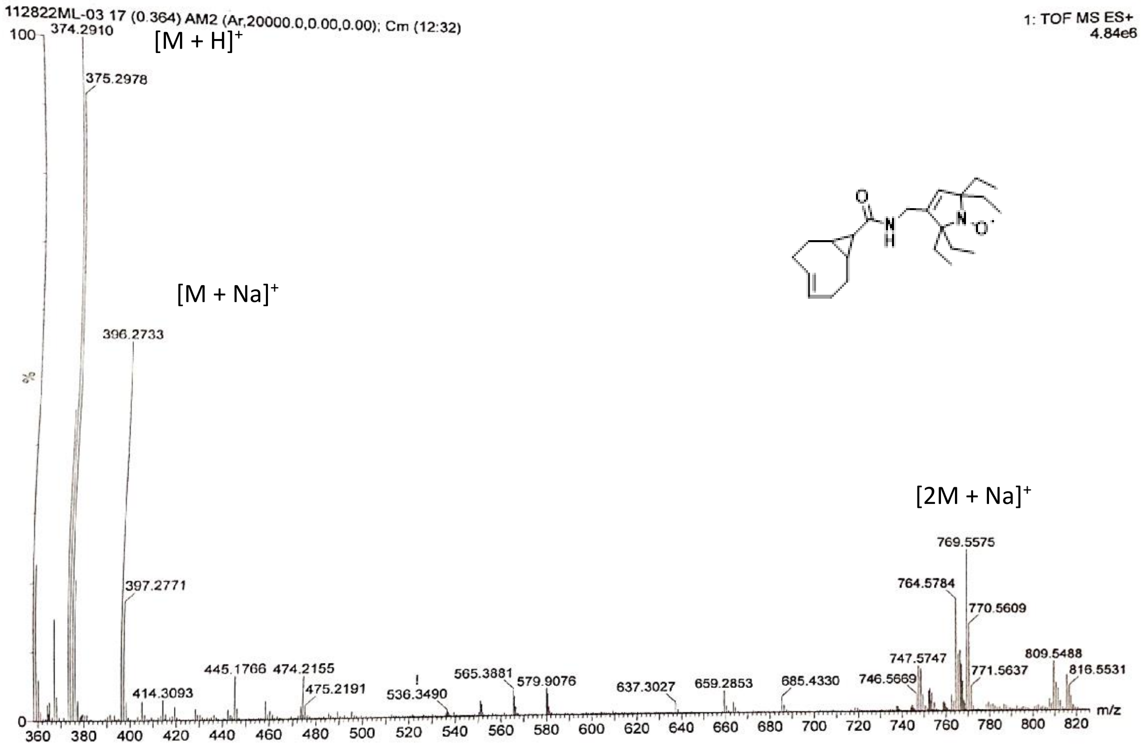
112822ML-03 17 (0.364) AM2 (Ar,20000.0,0.00,0.00); Cm (12:32)
1: TOF MS ES+
4.84e6
[M + H]+
374.2910
375.2978
[M + Na]+
396.2733
397.2771
[2M + Na]+
769.5575
764.5784
770.5609
809.5488
816.5531
771.5637
747.5747
746.5669
685.4330
659.2853
637.3027
579.9076
565.3881
536.3490
475.2191
474.2155
445.1766
414.3093
%
m/z
